# Collateral effects of lethal management operations: biologging reveals multiscale responses in non-target vervet monkeys

**DOI:** 10.64898/2026.07.23.739798

**Authors:** Loïc Brun, Josefien A. Tankink, Erica van de Waal

## Abstract

Lethal wildlife management operations are typically evaluated through their direct effects on target species, yet their disturbance footprint may extend to non-target animals sharing the same landscape. We opportunistically used a large-scale helicopter-assisted culling event targeting ungulates at Mawana Game Reserve, South Africa, as a natural experiment to examine how such an operation affected the behaviour and space use of non-target vervet monkeys (*Chlorocebus pygerythrus*). Using GPS and tri-axial accelerometry from 21 collared individuals across seven habituated social groups, we quantified behavioural and spatial responses around the culling event, including changes in activity budgets, circadian activity patterns, short-term walking and running responses, daily space use, and habitat composition. Vervets showed fear-consistent responses that differed among behaviours, social groups, and temporal scales. Foraging declined at the population level, with the strongest reduction occurring in the morning, when helicopter and shooting activity were the most intense. Inactivity increased during culling and remained elevated afterwards, suggesting both disturbance-related change and seasonal background effects, while social behaviour showed a transient increase consistent with a possible social-buffering response. Walking and running showed little consistent change in daily averages across the culling period, but finer-scale analyses revealed short-lived deviations concentrated in specific groups and time windows. Spatially, groups remained within existing ranges but showed divergent responses, from contraction and increased use of dense vegetation to greater movement in more open areas, highlighting how population-level inference can obscure context-dependent responses when heterogeneity among groups, individuals, or habitats is not explicitly considered. Together, our findings show that lethal management can generate multiscale, context-dependent responses in non-target wildlife, and that overlooking these effects may lead to systematic underestimation of their ecological consequences.

## Introduction

Human activities are among the most pervasive sources of disturbance for wildlife, reshaping ecosystems through habitat fragmentation and exploitation, climate change, infrastructure, vehicles, artificial light, noise, human presence, hunting, and wildlife management operations (Díaz et al., 2019; Venter et al., 2016). Understanding how animals respond to these pressures is therefore central to conservation biology and behavioural ecology, because behaviour is often the most immediate and flexible way in which animals adjust to rapid environmental change and adapt in human-altered landscapes (Berger-Tal et al., 2011; Tuomainen & Candolin, 2011; Wong & Candolin, 2015). Yet disturbance effects are difficult to quantify because responses may occur simultaneously across behavioural, physiological, temporal, and spatial scales, and because their expression depends on the interaction between disturbance type, environmental and demographic context, and the individual state (Pirotta et al., 2018; Tablado & Jenni, 2017).

This challenge is especially pronounced for lethal management operations such as hunting and culling, which are intense, often short-lived, and usually assessed through their direct effects on target species. These operations create mortality risk for target animals but also generate sensory cues, such as gunfire, helicopter and other vehicle noise, and sudden human presence, that can propagate beyond the immediate location of killing and signal danger to non-target animals (Beale & Monaghan, 2004; Erbe et al., 2022; Frid & Dill, 2002). The lethal anthropogenic disturbance literature has focused primarily on target species showing that culling disrupts spatial organisation and alters movement patterns in badgers (Ham et al., 2019; Woodroffe et al., 2006), elevates vigilance and physiological stress in fallow deer (Pecorella et al., 2016) and recent work on wild pigs suggests that intensive culling can alter space use and contact patterns (Chalkowski et al., 2025). In contrast, collateral responses in non-target species remain poorly documented. The available evidence comes mainly from birds, where shooting operations can cause spatial displacement or incidental mortality of non-target species (Marchowski et al., 2025; Ozsanlav-Harris et al., 2024). In mammals, evidence remains comparatively scarce and mostly concerns ungulates, particularly deer, with studies showing that hunting can alter refuge and habitat use, movement, or activity timing in non-target species or non-target demographic groups exposed to hunting disturbance (Brown et al., 2020; Grignolio et al., 2011; Peters et al., 2025). Yet these studies largely concern spatial or temporal avoidance, leaving broader activity-budget responses in non-target mammals, especially outside ungulates, poorly documented - a gap explicitly identified in a recent systematic review calling for studies to broaden their focus beyond target species and targeted demographic classes (Mysterud, 2026).

Despite this gap, predictions about how non-target animals may respond can be guided by theoretical frameworks linking perceived threat to behaviour. The risk-disturbance hypothesis proposes that human-induced stimuli are processed through the same threat-detection systems as natural predator cues, triggering comparable behavioural responses even when animals are not directly targeted (Frid & Dill, 2002). Empirical evidence from non-lethal human disturbance supports this: elk reduce foraging more strongly under human presence than under wolf predation risk, and pumas substantially shorten feeding time when exposed to human vocal cues, consistent with the view that humans act as a ’super predator’ whose cues elicit fear responses exceeding those of natural predators (Ciuti et al., 2012; Darimont et al., 2015; Smith et al., 2017). These shifts are expected to vary across both space and time. The landscape of fear predicts that animals adjust their behaviour according to spatial variation in perceived danger, contracting space use, avoiding risky areas, or shifting towards safer habitat (Gaynor et al., 2021; Laundré et al., 2010), while the risk allocation hypothesis predicts that riskier activities such as foraging should shift temporally into periods of lower perceived threat (Lima & Bednekoff, 1999). Such temporal reorganisation is widely documented, with mammals globally increasing nocturnality in response to human presence (Gaynor et al., 2018). For non-target species sharing the landscape with culled animals, culling-associated cues may function as threat signals, producing behavioural trade-offs between resource acquisition and safety, even in animals that are never directly pursued or shot.

A large-scale helicopter-assisted culling operation targeting multiple ungulate species at Mawana Game Reserve, a private game reserve in South Africa, provided a natural experiment to examine how non-target vervet monkeys (*Chlorocebus pygerythrus*) responded to an active lethal management event. The operation lasted one week, during which approximately 600-700 animals, ranging from impala to giraffe, were killed for their meat. Although such operations are usually planned by reserve managers, opportunities for independent researchers to document them are rare. Fine-scaled behavioural data are difficult to acquire because direct observation is typically curtailed for safety reasons once active culling begins. In this case, a concurrent biologging project provided high-resolution GPS and accelerometry data across the event and surrounding weeks, allowing us to quantify movement, space use, and activity budgets of vervet monkeys throughout. Animal-borne accelerometers capture fine-scale body movements associated with behavioural states, while GPS tracking provides records of spatial position and movement. Together, these tools can resolve responses at temporal scales that would be difficult to capture through intermittent observation alone (Nathan et al., 2012; Shepard et al., 2008). This proved particularly valuable here because disturbance responses may range from sustained shifts across the full culling period to brief, acute responses concentrated in periods of likely heightened disturbance intensity. Yet, despite the rapid expansion of tracking and biologging technologies and their clear management relevance, their use in applied conservation settings remains comparatively limited, particularly for documenting behavioural responses to management operations as they unfold (Katzner & Arlettaz, 2020).

Vervets at Mawana are monitored by the Inkawu Vervet Project (IVP), an independent long-term research programme that has followed wild but habituated groups since 2010, providing detailed knowledge of group composition, life histories, and ecological context. Vervet monkeys exhibit predator-specific antipredator responses, adjust their activity budgets flexibly across ecological and social conditions and live in stable multi-male, multi-female groups within established home ranges (McFarland et al., 2014; Seyfarth et al., 1980). These characteristics make them a particularly relevant model for examining whether non-target disturbance produces behavioural and spatial reorganisation without necessarily causing large-scale displacement. In territorial primates, abandoning familiar areas incurs high costs, including reduced access to known food resources and refuges, increased exposure to predators, and greater risk of conflict with neighbouring groups (Isbell et al., 1993; Willems & Hill, 2009). Under short-term disturbance, spatial responses may therefore be more likely to involve adjustments within existing home ranges than displacement into unfamiliar areas. Their sociality also allows us to examine a complementary dimension of response, namely whether the culling disturbance was associated with changes in affiliative behaviour within the group. Under the social buffering hypothesis, affiliative interactions are expected to attenuate physiological stress responses, and in primates, behaviours such as grooming are associated with reduced glucocorticoid responses under challenging conditions, particularly among strongly bonded individuals (Sanchez et al., 2015; Young et al., 2014). Consistent with this, primates may increase grooming rates or broaden social connections following acute stressors, including natural disasters and anthropogenic disturbance (Semple et al., 2013; Testard et al., 2021).

Building on these predictions, we examined whether the culling operation altered vervet behaviour and space use across temporal and spatial scales. Specifically, we asked whether responses were restricted to the culling period or persisted afterwards, whether they were sustained across the disturbance window or concentrated at particular times of the day, and whether they were consistent across social groups. We predicted that culling-associated disturbance would reduce foraging and increase inactivity, reflecting a safety-resource trade-off under perceived threat (Frid & Dill, 2002). Because helicopter and shooting activity were reported to be most intense in the morning, we expected this behavioural reallocation to be strongest early in the day, consistent with the risk allocation hypothesis (Lima & Bednekoff, 1999). Given the role of affiliative behaviour in primate stress regulation, we further predicted a transient increase in social behaviour during the culling period, consistent with the social buffering hypothesis (Sanchez et al., 2015; Young et al., 2014). Spatially, we expected groups to remain within familiar ranges while reducing daily movement, contracting space use, or shifting toward denser vegetation that could provide refuge. Finally, because groups differed in habitat structure and refuge availability, and likely also in their exposure to disturbance, we primarily expected movement and spatial responses to be heterogeneous across groups, with brief locomotor responses potentially concentrated around periods of heightened disturbance rather than expressed uniformly across the full culling period. By combining continuous biologging with a rare natural experiment created by an active lethal management operation, this study shows how acute human disturbance can reshape the behaviour and space use of a non-target primate, and assesses whether the ecological footprint of culling extends beyond target-species mortality to affect wider wildlife communities.

## Methods

### Study site and culling event

This study was conducted at Mawana Game Reserve (28°00.327′S, 31°12.348′E), a mesic savannah of approximately 11,000 ha in KwaZulu-Natal, South Africa, characterized by a cold and dry winter and hot summer with important rainfall. The vegetation mostly consists of Acacia woodland (mainly *Vachellia tortilis* and *Vachellia nilotica*), spiny shrubs, and open grass patches (Mucina & Rutherford, 2006). Vegetation structure varies considerably across the reserve, ranging from open savannah to dense riverine areas, resulting in a heterogeneous landscape.

A large-scale culling event took place from 15 to 21 May 2023, authorised by Ezemvelo KZN Wildlife (Permit OP 1538/2023) and targeting multiple ungulate species for commercial meat harvesting (planned quotas: kudu *n* = 500, impala *n* = 1,000, blue wildebeest *n* = 1,000, zebra *n* = 200, blesbok *n* = 400, giraffe *n* = 100, nyala *n* = 300, warthog *n* = 100, waterbuck *n* = 50). Culling involved helicopter-assisted herding and direct aerial shooting across the reserve, with carcasses processed on site and transported in refrigerated trucks. The operation was interrupted before reaching these targets, with around 661 animals removed. Culling intensity was higher in the northern sector encompassing the IVP research area, and helicopter and shooting activity was concentrated mostly in the morning. However, because the culling was not anticipated as part of this study, we had limited information on the precise spatial and temporal distribution of disturbance. We therefore treated it as a discrete seven-day event and defined three analysis phases: *before*, *during*, and *after* culling. To balance statistical power against potential seasonal confounding, we chose a temporal window spanning 15 days on either side of the culling period to define the after and before phase.

### Study system and animals

At the time of the study, seven habituated groups (Fig.1), each comprising approximately 20–60 individuals, were monitored daily by trained field assistants as part of the IVP. As direct observations were reduced for safety during culling, we relied exclusively on concurrent biologging data to ensure continuous behavioural coverage across groups and days. Although not originally designed to study disturbance responses, this dataset provided continuous behavioural and movement information across the culling event. Collars (e-obs GmbH, Germany) combined GPS and tri-axial accelerometer sensors, enabling space use and behaviour to be quantified from the same individuals. Between two to seven monkeys were collared per group, resulting in 21 tagged individuals (6 females and 15 males; Table S1). All tagged individuals were confirmed residents of their respective groups throughout the study window, so analyses reflected established group members rather than dispersing or newly integrated individuals.

**Figure 1.**
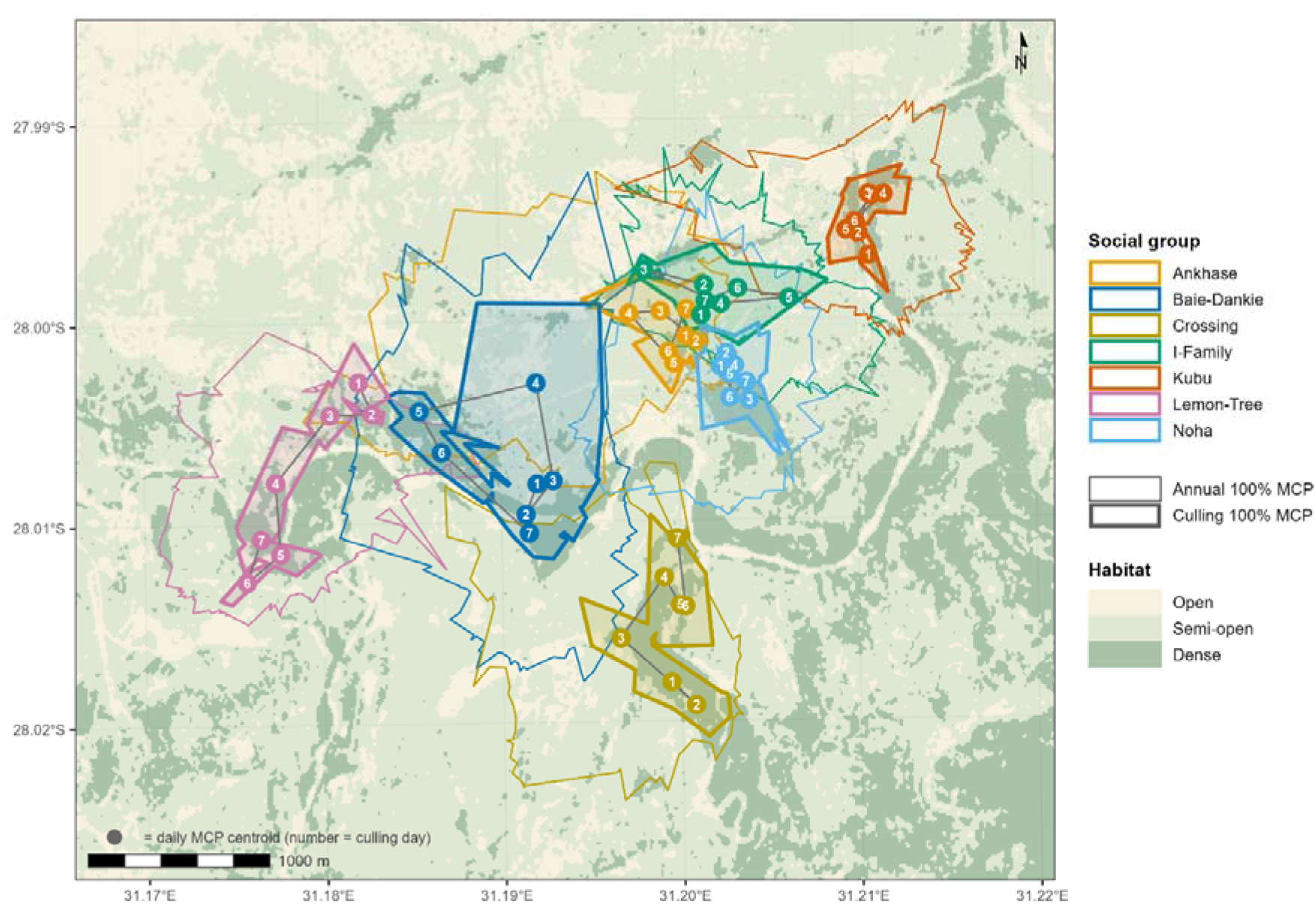
Study area, group ranges, and culling-period space use. Thin coloured outlines show each group’s observed home range over one year of GPS monitoring, approximated as the union of daily 100% minimum convex polygons (MCPs). Thicker outlines show the union of daily 100% MCPs calculated during the culling period, and numbered points indicate daily MCP centroids for each culling day. Background shading represents habitat classes derived from NDVI thresholds: open, semi-open, and dense vegetation.

### Ethical Statement

Captures for collar deployment were conducted by a certified veterinarian, following protocols approved by the University of KwaZulu-Natal (UKZN) Animal Ethics Committee and by the European Research Council, in accordance with Horizon Europe regulations. EvdW is affiliated to UKZN and collaborates with Prof. Collen Downs who has been granted animal ethics permission to capture, fit loggers and release vervet monkeys with the assistance of a veterinarian (application AREC/00005320/2023(00020574), Project title: Ecosystem health and biodiversity: The ecology, physiology, behaviour and conservation of selected southern African vertebrates). Only individuals (6 females, 31 males) for whom the collar mass represented < 3% of total body mass were selected for deployment. At the end of the study, another capture session took place to remove the collars.

### GPS data and spatial metrics

GPS collars recorded locations every 2 h for females and every 4 h for males. Locations were filtered prior to analysis to remove low-quality fixes (SM1). From these data, we derived two spatial metrics: daily path length and daily space use. Daily path length was calculated for each individual as the sum of step lengths between successive locations within a day. Because sampling interval differed between sexes and directly influences movement estimates, trajectories were downsampled to 4 h for each individual, and only days with 5-6 fixes per individual were retained, ensuring comparable movement estimates across individuals (SM2).

Daily space use was quantified using 100% minimum convex polygons (MCPs), computed using the *amt* R package (Signer et al., 2019). MCPs were preferred over kernel-based estimators, which produced unstable estimates at the daily scale (SM3). MCPs were computed at the group-day level from a single focal female per group using the higher-resolution 2-h dataset. We used one single female individual per group for three reasons. First, vervet groups are cohesive and maintain relatively constrained group spread (Noë & Laporte, 2014; Struhsaker, 1967), meaning a single individual represent a consistent proxy for the group, especially females who typically occupy more central positions than the more peripheral males (Isbell et al., 1993). Second, their 2-h sampling schedule provided finer spatial resolution than the 4-h schedule used for males (SM2). Third, MCP area is sensitive to the number of individuals contributing locations, so this approach avoided overrepresenting groups with more collared individuals. One group (I-Family) was excluded from MCP-based analyses because no collared female was available. To maximize the number of days per group with representative MCPs, we only kept days with 7-12 fixes per female, which delivered relatively stable estimates of daily space use areas (SM2). Within each daily MCP, habitat composition was quantified from an NDVI-derived vegetation layer, with pixels classified as open (NDVI < 0.25), semi-open (0.25 ≤ NDVI < 0.33), or dense (NDVI ≥ 0.33).

Analyses retained the proportional cover of dense and semi-open vegetation only, because open habitat proportions were rare across daily MCP areas and mathematically connected to the two other categories (SM4).

### Behavioural classification from accelerometry

Tri-axial accelerometer data were recorded at 10 Hz in 13.8-s bursts every 90 s. Behavioural states were inferred using two supervised deep-learning classifiers, HydraROCKET and TabPFN, both trained on labelled vervet monkey accelerometry data (Brun et al., 2026; preprint). The two classifiers operated at different temporal resolutions and showed complementary strengths: HydraROCKET was applied to the full 13.8-s burst and was better suited to sustained behaviours, whereas TabPFN was applied to four shorter sub-bursts of approximately 3.4 s each, derived by dividing the original burst into four segments, and improved the detection of shorter-lived or postural behaviours. The full classification and consensus procedure are described in Supplementary M5.

Both classifiers independently predicted eight initial behavioural categories: eating, walking, running, resting, sleeping, grooming actor, grooming receiver, and self-scratching, although the latter category was discarded from subsequent analyses (Supplementary M5.2). Foraging was defined as active food-searching and captured bursts combining locomotor and feeding components, specifically cases in which HydraROCKET predicted eating or walking and TabPFN identified a mixture between eating and walking across the four sub-bursts. This category was kept distinct from eating either when both classifiers predicted eating or when TabPFN identified eating mixed with resting sub-bursts. Eating therefore primarily corresponded to stationary feeding, often in bushes or trees, and was expected to differ from ground-based foraging in its exposure to disturbance. Resting and sleeping were merged into inactive because these states were difficult to distinguish reliably from accelerometry alone and were expected to have similar relevance for the disturbance analyses. Grooming actor and grooming receiver were merged into social behaviour because both represented affiliative social interactions and were expected to carry similar biological meaning in response to the disturbance. Bursts showing complete disagreement between classifiers, or falling below the minimum confidence threshold, were excluded following the consensus procedure described in SM5. All analyses were restricted to daytime bursts.

### Spatial responses to culling

To assess spatial responses to the culling event, we analysed daily path length, daily MCP area, and vegetation composition within daily MCPs across the three disturbance phases. Daily path length and MCP area were modelled using GLMMs with a Gamma error distribution and log link, whereas vegetation proportions were analysed using beta GLMMs with a logit link. Daily path length analyses included all 21 collared individuals. Standardising trajectories to a 4-h interval resulted in 5–6 fixes per individual per day, equalising sampling effort across individuals and removing the need to include fix number as a covariate. Models included individual identity and date as random intercepts. Sensitivity analyses using the raw data with up to 13 GPS fixes and fix number included as a covariate confirmed that effect directions were consistent across datasets (SM2).

Daily MCP area and vegetation metrics were analysed at the group-day level using the 2-h female dataset, with date included as a random intercept when supported by model fit. For all responses, we compared interaction, additive, population-level, and null model structures using Akaike’s Information Criterion (AIC; Table S2). For daily path length, inference followed the best-supported structure. For MCP and vegetation metrics, interaction models were retained regardless of AIC ranking for three reasons. First, groups occupied distinct areas of the reserve and were expected to differ in their exposure to culling disturbance, making group-specific responses the biologically meaningful target. Second, with only one observation per group per day, replication was inherently low, limiting the reliability of AIC selection. Finally, model selection favoured an interaction structure for daily path length, which was expected to correlate at least partially with MCP area. Thus, retaining the same structure for space-use metrics ensured consistency across related responses.

### Daily behavioural responses from accelerometry

Behavioural activity was quantified as the daily proportion of daytime accelerometry bursts assigned to each behavioural category for each individual and was calculated relative to the total number of bursts per day. Analyses were based on 20 individuals only as one female from Crossing group (CR), Nihau, was excluded due to a reduced accelerometer sampling rate that produced atypical behavioural patterns. Responses were modelled using beta GLMMs with a logit link, with event phase as the primary fixed effect. For each behaviour, we compared three candidate model structures: a population-level model with phase as a fixed effect and group as a random effect, an additive model with phase and group as fixed effects, and an interaction model allowing group-specific phase responses. All models included individual identity and date as random intercepts. Unlike the MCP-based analyses, model selection was applied as usual here, as the greater replication provided by multiple individuals and bursts per day gave AIC sufficient sensitivity to distinguish among structures. Importantly, the selected model structure was itself biologically informative: a population-level model indicated a response shared across groups, whereas an interaction model indicated group-specific variation in the direction or magnitude of the response. The best-supported structure for each behaviour is reported in Table S3.

### Statistical analyses

All models were fitted in R using the *glmmTMB* package (Brooks et al., 2017). Model fit was evaluated using simulation-based diagnostics implemented in *DHARMa* (Hartig, 2024; Figs. S1 and S2), including tests of residual uniformity, dispersion, and outliers. Statistical inference was based on estimated marginal means and pairwise contrasts between phases computed on the response scale using the *emmeans* package (Lenth & Piaskowski, 2025), with p-values adjusted for multiple testing using the Benjamini–Hochberg false discovery rate (FDR) procedure (Benjamini & Hochberg, 1995). Model selection across all analyses was based on AIC (Burnham & Anderson, 2002).

### Temporal trajectories and circadian organisation

Phase contrasts summarise average behavioural change across broad multi-day windows, but may miss finer-scale responses and make it difficult to distinguish disturbance-induced shifts from gradual background changes over the study period. We therefore complemented phase contrasts with daily trajectories and circadian analyses. A culling-induced response was expected to show stronger temporal alignment with the disturbance window, whereas background drift was expected to produce a smoother progressive change across the full study period. Daily trajectories assessed whether behavioural changes were centred on the culling window, while circadian analyses identified whether phase differences were concentrated at particular times of day or reflected broader deviations from baseline.

Population-level daily trajectories were modelled using generalised additive models (GAMs) implemented in the *mgcv* package (Wood, 2017), with daily behavioural proportions averaged across individuals, and fitted as smooth functions of time relative to the start of culling under a beta distribution with a logit link. The smoothing parameter *k* was constrained to a maximum of 10 to avoid overfitting. At the group level, where daily means were based on ess data and fewer individuals, *LOESS* smoothers were used instead. Group-level trajectories are presented in the supplementary material to illustrate heterogeneity among groups and short-lived responses that may be diluted in population-level summaries.

For circadian analyses, accelerometry bursts were aligned relative to sunrise rather than clock time to reduce seasonal variation in daylight timing, and behavioural activity was summarised in 30-minute bins across the daytime period. Bursts recorded at dawn before sunrise and at dusk after sunset were retained as part of the active period, resulting in curves that extend slightly beyond the sunrise and sunset reference points. Smooth curves of behavioural proportion as a function of hours relative to sunrise were compared among phases, first at the group level and then at the population level.

### Fine-scale locomotor responses during culling

Daily averages and smooth trajectories may obscure short-lived responses to acute disturbance. We therefore quantified sub-daily deviations from baseline during the culling period for all six behavioural categories (Fig. S6). Behaviours with higher baseline frequencies (i.e. eating, foraging, inactive, and to a lesser extent social) were already well captured by the daily and circadian analyses and did not reveal additional patterns. We therefore focus here on walking and running, which have lower baseline frequencies, making proportional departures harder to detect with GAMs and phase contrasts, yet are the behaviours most likely to reflect immediate responses to disturbance cues such as helicopter flights or gunshots.

Bursts were aligned relative to sunrise and deviations were quantified across three consecutive time windows spanning the first 10.5 hours of the active day: morning (0–3.5 h), midday (3.5–7 h), and late afternoon (7–10.5 h) after sunrise. For each group, behaviour, and time window, expected activity during culling was estimated from the corresponding baseline proportion in the before phase. Deviations were quantified in two complementary ways. First, as Pearson-type standardised residuals, r = (O − E) / √(E(1 − p)), where O is the observed burst count, E = np the expected count under baseline probability p, and n the total bursts in that group, day, and time window. Second, we calculated raw percentage-point differences between observed and expected proportions. Standardised residuals indicate how unusual a deviation was relative to baseline variability, whereas percentage-point differences capture its absolute magnitude.

These analyses required sufficient baseline coverage to estimate stable group-specific reference distributions. Crossing (CR) was therefore excluded because, following the removal of Nihau, only one collared individual remained in that group, providing insufficient data to estimate a reliable baseline, particularly for lower-frequency behaviours.

## Results

### Daily path length

Daily path length responses varied among groups and no before-during difference remained statistically supported after FDR correction (Fig. 2a; Table S4). Nevertheless, four groups showed directional reductions during culling relative to baseline, ranging from approximately 15% to 21% (AK: −21%, CR: −19%, NH: −19%, KB: −15%). Evidence for these reductions was strongest in AK, NH, and CR, with adjusted p-values close to the significance threshold (p = 0.06, 0.05, and 0.09, respectively). By contrast, the remaining groups (BD, IF and LT) showed estimated changes close to 0 or slight increases in daily path length. A general rebound in movement after culling was suggested by the during-after contrasts, which was strongest and only statistically supported in LT (+40%, p = 0.009), with KB showing a borderline increase (+28%, p = 0.06). Before−after contrasts were more heterogeneous. LT showed the only significant increase relative to baseline (+32%, p = 0.007), while most other groups showed lower and non-significant changes ranging from −12% in CR to +22% in IF.

**Figure 2.**
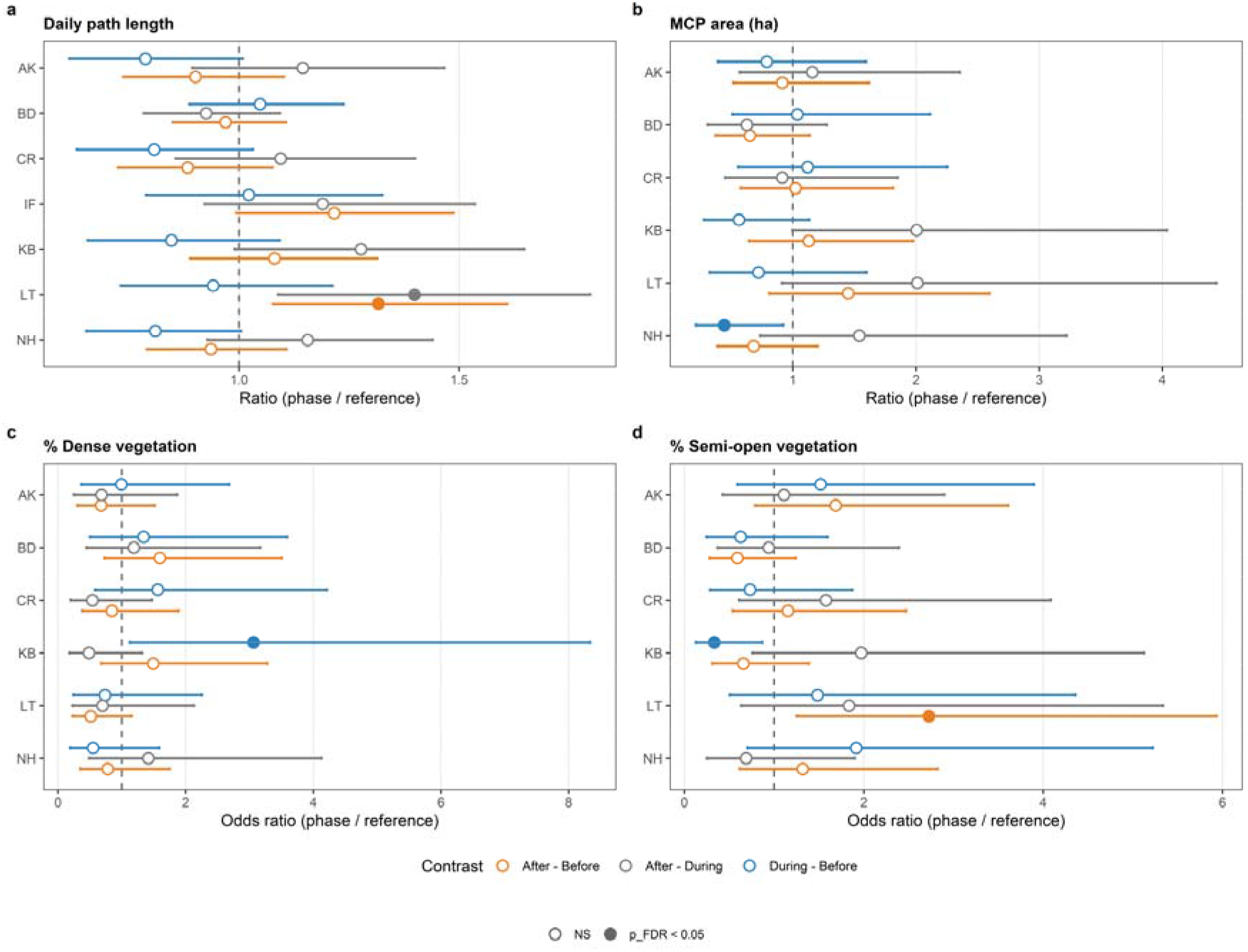
Estimated effects of the culling event on (a) daily path length, (b) daily MCP area, (c) dense vegetation within daily MCPs, and (d) semi-open vegetation within daily MCPs. Points represent model-estimated pairwise contrasts between phases, expressed as ratios (daily path length and MCP area; Gamma GLMMs) or odds ratios (vegetation proportions; beta GLMMs), with horizontal lines indicating 95% confidence intervals. The vertical dashed line indicates no effect (ratio = 1). Colours denote phase contrasts (blue: during − before; grey: after − during; orange: after − before), and filled symbols indicate statistically significant effects after FDR correction (p < 0.05). Estimates are shown separately for each group.

### Daily space use and habitat composition

Daily MCP area responses were generally stronger than those observed for daily path length, though similarly heterogeneous across groups (Fig. 2b; Table S4). During culling, NH showed the only statistically supported reduction in space use relative to baseline (−56%, p = 0.03), with comparable but non-significant decreases in KB (−44%, p = 0.11) and LT (−28%, p = 0.42), whereas BD and CR showed little change. Post-culling contrasts indicated a rebound in space use in several groups, with KB and LT approximately doubling their MCP area after culling relative to during culling (+101% each; p = 0.05 and p = 0.08 respectively), while NH also showed a weaker increase (+54%, p = 0.25). BD showed a further decrease after culling relative to during culling (−37%, p = 0.20). Before−after contrasts showed no consistent direction across groups and remained statistically unsupported.

Habitat composition within daily MCPs differed most clearly in KB, which showed a significant increase in dense vegetation use during culling (+25 percentage points, p = 0.03) and a corresponding reduction in semi-open vegetation (−24 percentage points, p = 0.02). LT showed a statistically supported increase in semi-open vegetation after culling relative to baseline (+22 percentage points, p = 0.01). Remaining contrasts were not supported after correction (Fig. 2c-d; Table S4).

Daily temporal trajectories revealed substantial day-to-day variability in path length, MCP area, and habitat composition across the culling period, with marked fluctuations between consecutive days and several isolated extreme values (Fig. S3). In some groups, phase-level estimates appeared to be strongly influenced by single days, most notably in BD where an isolated increase on 18 May appeared to drive the overall culling-phase pattern. These trajectories suggest that spatial responses were not sustained uniformly across the full culling week but concentrated on particular days.

### Acceleration-based behaviour

Behavioural responses to the culling event varied across groups and behaviours, with model selection supporting population-level responses for foraging and social behaviour, and group-specific responses for walking, eating, inactive behaviour, and running (Table S3; Figs. 3–4). Estimated marginal means and phase contrasts are reported in Table S5. Foraging showed the clearest and most consistent response. Mean foraging activity declined from 19.8% before culling to 17.7% during culling (−2.1%, p = 0.02) and remained lower after culling than before (18.0%, −1.7%, p = 0.02), whereas the during-after contrast was not supported (p = 0.67). The population GAM confirmed this pattern, showing a decline clearly centred on the culling window with a partial recovery after day 7 that did not return to pre-culling levels by the end of the study period (Fig. 3b).

**Figure 3.**
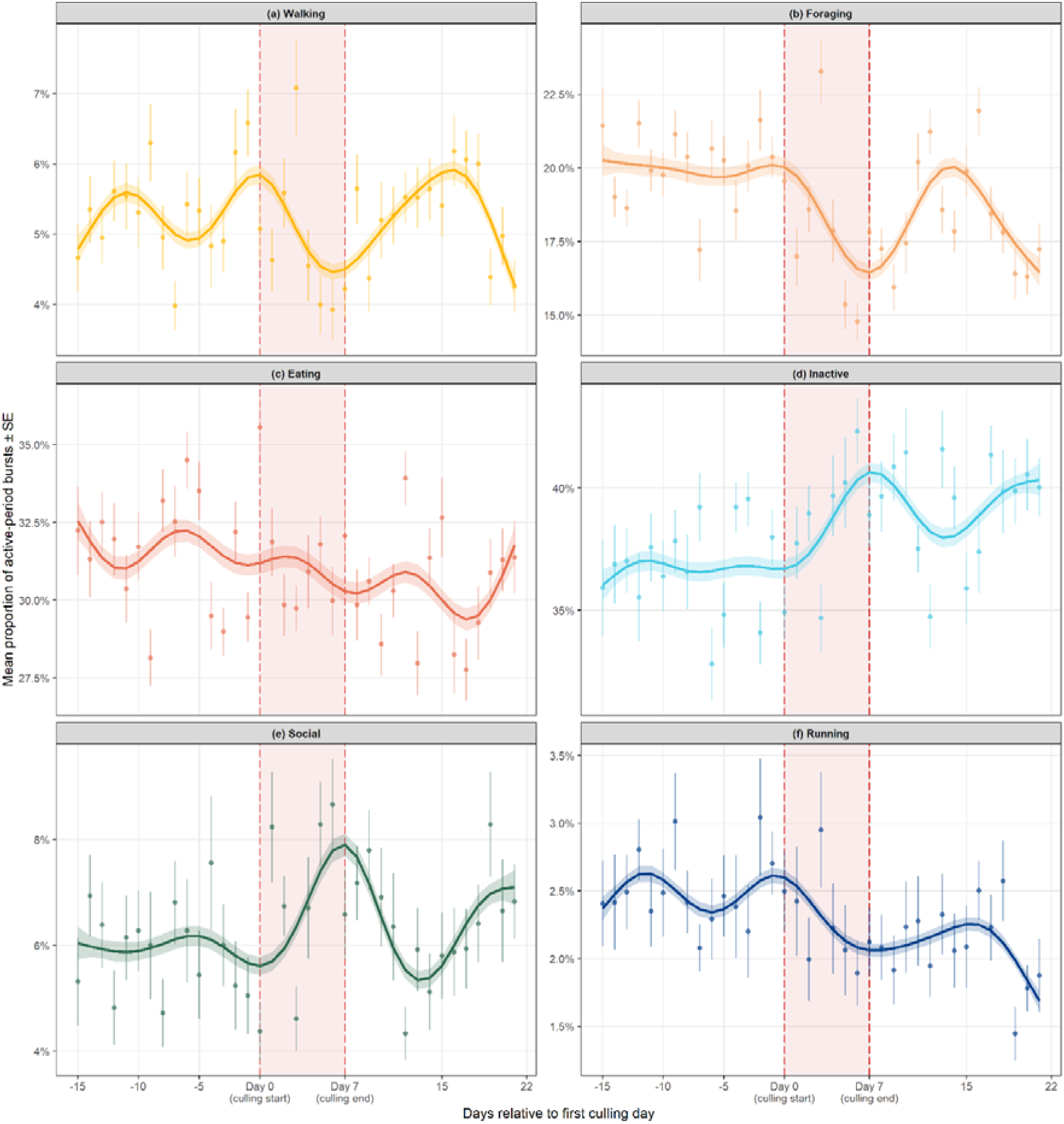
Population-level GAM trajectories of behavioural activity across the culling event for (a) walking, (b) foraging, (c) eating, (d) inactive behaviour, (e) social behaviour, and (f) running. Points show daily population means ± SE and solid lines show fitted beta GAM trajectories ± 95% CI ribbons. The x-axis shows days relative to the first culling day (day 0); red shading and dashed red vertical lines indicate the culling period (days 0–7).

**Figure 4.**
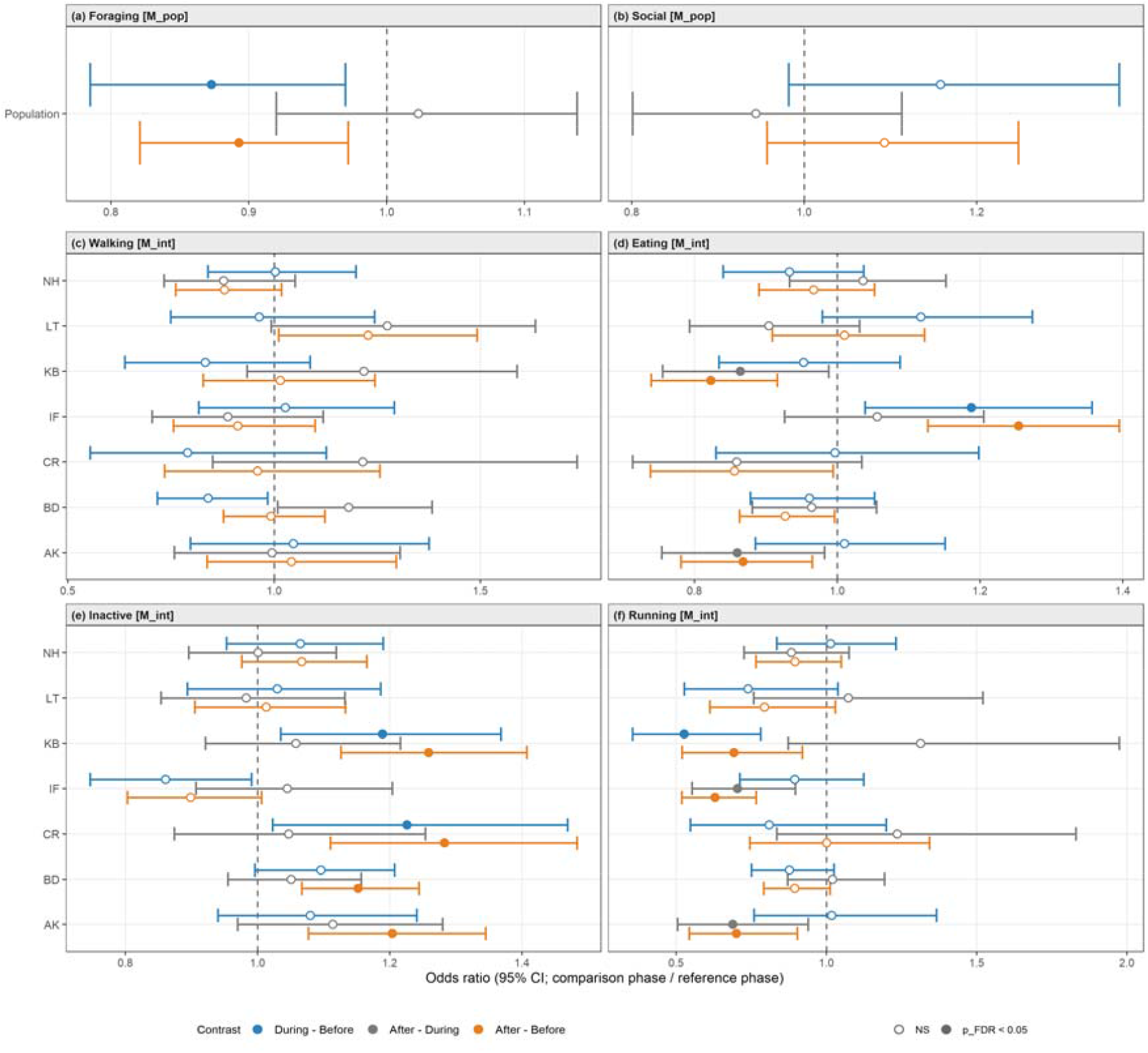
Phase contrasts for accelerometry-derived behavioural responses to culling for (a) foraging, (b) social behaviour, (c) walking, (d) eating, (e) inactive behaviour, and (f) running. Points represent model-estimated pairwise contrasts between phases, expressed as odds ratios (OR) from beta GLMMs with logit link; horizontal lines indicate 95% confidence intervals. The vertical dashed line marks no effect (OR = 1). Colours denote phase contrasts: blue = during − before, grey = after − during, and orange = after − before. Filled symbols indicate contrasts that were statistically supported after FDR correction (p < 0.05). Population-level estimates are shown in the top row for foraging and social behaviour, whereas group-specific estimates are shown for walking, eating, inactive behaviour, and running, in line with model selection results (Table S3).

Social behaviour showed a weaker parallel pattern. Mean social activity increased from 5.4% before to 6.2% during culling (+0.8%, p = 0.25) and remained slightly elevated after culling (5.9%, +0.5%, p = 0.28), though neither contrast was statistically. The population GAM revealed an important peak clearly centred on the culling window, reflecting the sharpest increase relative to baseline across all behaviours and followed by a rapid decline to pre-culling levels within approximately five days after the end of culling. This pattern being consistent across most groups in the group-level trajectories (Figs. 3e, S4e).

Inactive behaviour showed a directionally consistent increase across groups. During culling, statistically supported increases were observed in CR (+5.0%, p = 0.04) and KB (+4.1%, p = 0.02), while most other groups showed directional but non-significant increasing trends, and only IF showed a decreasing trend. Before-after contrasts revealed stronger and more widespread elevations, with significant increases in AK (+4.5%, p = 0.003), BD (+3.3%, p < 0.001), CR (+6.2%, p = 0.002), and KB (+5.5%, p < 0.001), while during-after contrasts were not supported in any group. The population GAM showed a steady increase in inactivity from the onset of culling that continued beyond day 7 without clear recovery and visible in most groups (Figs. 3d, S4d).

Eating showed no consistent response across groups. Phase contrasts were not supported during culling in any group except IF, which showed a directional increase (+3.6%, p = 0.02). The strongest contrasts emerged in the before-after comparison but reflected opposing patterns: KB and AK showed a significant decrease (−4.1%, p = 0.001; −3.0%, p = 0.03 respectively) while IF increased markedly (+4.8%, p < 0.001). During-after contrasts were similarly mixed, with lower eating after culling in AK (−3.3%, p = 0.04) and KB (−3.0%, p = 0.05). The population GAM showed no clear culling-centred structure, and group-level trajectories confirmed that changes in eating were often not aligned with the culling window (Figs. 3c, S4c).

Walking phase contrasts were small and non-significant across all groups, with the largest before-during decrease in BD (−1.0%, p = 0.06). The population GAM showed a modest decrease during culling but with a lot of oscillations around the disturbance, consistent with the heterogeneity visible in the group-level trajectories. Nevertheless, several groups, notably CR, NH, and BD, showed short-lived declines perfectly centered around the culling and most group showed a clear rebound shortly afterwards (Figs. 3a, S4a).

Running generally showed a directional decrease relative to baseline. The only supported during−before contrast was in KB (−0.7%, p = 0.004), while most other groups showed non-significant decreasing trends. Before-after contrasts indicated significant reductions in IF (−1.4%, p < 0.001), AK (−0.6%, p = 0.02), and KB (−0.4%, p = 0.02), with during-after contrasts also showing lower running in IF (−1.0%, p = 0.007) and AK (−0.6%, p = 0.03). The population GAM suggested a slight elevation immediately before culling followed by a decline during and after the event. This pattern was further confirmed to be shared across several groups, notably IF, CR, and NH, which all showed a peak just before or at the start of culling followed by a sharp decline (Figs. 3f, S4f). Notably, walking and running were the only two behaviours showing a pre-culling signal, with an increasing trend in the days immediately before the event followed by a decline at or shortly after its onset — a pattern visible in several groups for both behaviours (walking: CR, NH; running: IF, CR; Figs. S4a, f).

### Circadian organisation of behaviour

Circadian analyses revealed that, in addition to changes in total daily activity budgets, culling altered the temporal distribution of behaviour within days, with the strongest phase differences concentrated within the first two to four hours after sunrise (Fig. 5; Fig. S5). In line with the phase contrast, foraging showed the strongest circadian signal. The during phase was markedly reduced relative to before within the first four hours after sunrise. Confidence intervals did not overlap during this period, indicating a reduction reaching approximately 7%, which is substantially larger than the 2% reduction observed at the daily scale, before the three phases gradually converged towards midday. Comparison of the during and after curves further showed that the during phase remained slightly lower for longer into the morning, consistent with a response more closely tied to the culling period than to seasonal drift alone (Fig. 5b).

**Figure 5.**
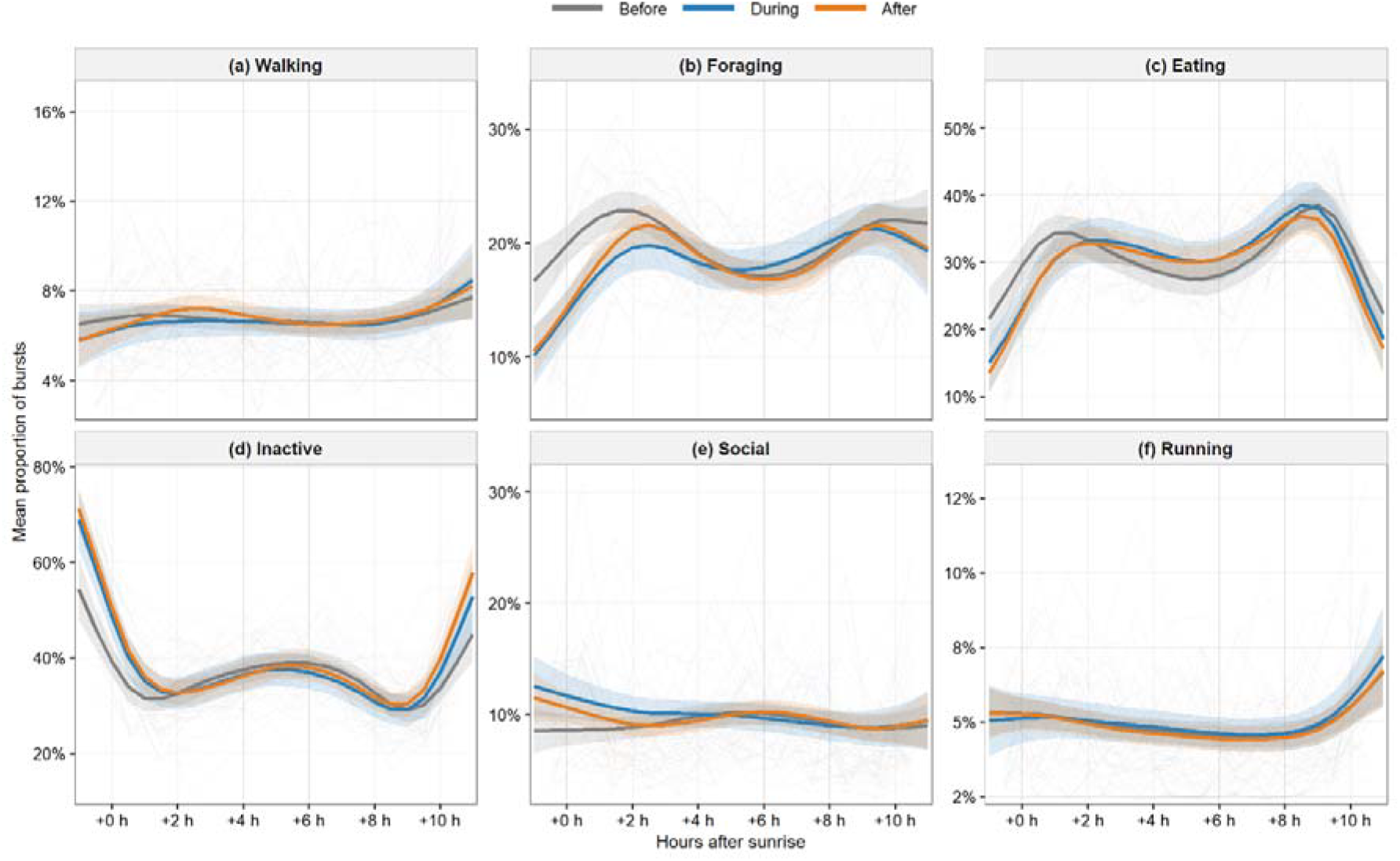
Population-level circadian organisation of behaviour across disturbance phases. Behavioural proportions were aligned relative to sunrise and compared across phases before, during, and after the culling event. Solid lines show phase-specific beta GAM smooth trajectories, and shaded ribbons indicate 95% confidence intervals.

Eating showed a broadly parallel pattern, with a visible reduction in the during phase in the morning relative to before, although the during and after phases were very similar across the whole day, slightly contrasting with foraging. Both phases also showed a directional increase in the afternoon relative to before, although confidence intervals overlapped in this case (Fig. 5c). Social behaviour showed a similar morning deviation across phases, although the during phase was more clearly separated from both before and after compared to both eating and foraging, before the three phases converged later in the day here again (Fig. 5e). Inactive behaviour was elevated in the first two hours after sunrise in the during phase relative to before, mirroring the morning reduction in foraging and eating. The during and after phases were mostly similar across the day, consistent with the more sustained elevation observed in the daily phase contrasts (Fig. 5d).

Walking and running showed greater overlap among phases at the population level, indicating weaker redistribution of activity across the day or at least that no patterns were clearly captured when averaging across days (Figs. 5a, f). Overall, at the group level, the before phase was relatively homogenous in shape across groups for most behaviours, whereas the during and after phases showed greater divergence, with some groups showing patterns opposite to the population trend (Fig. S5).

### Fine-scale locomotor responses during culling

Walking and running showed weak or inconsistent phase contrasts and circadian patterns, yet fine-scale analyses revealed substantial within-day deviations from baseline with interesting patterns emerging across days and time windows (Fig. 6; Fig. S6).

**Figure 6.**
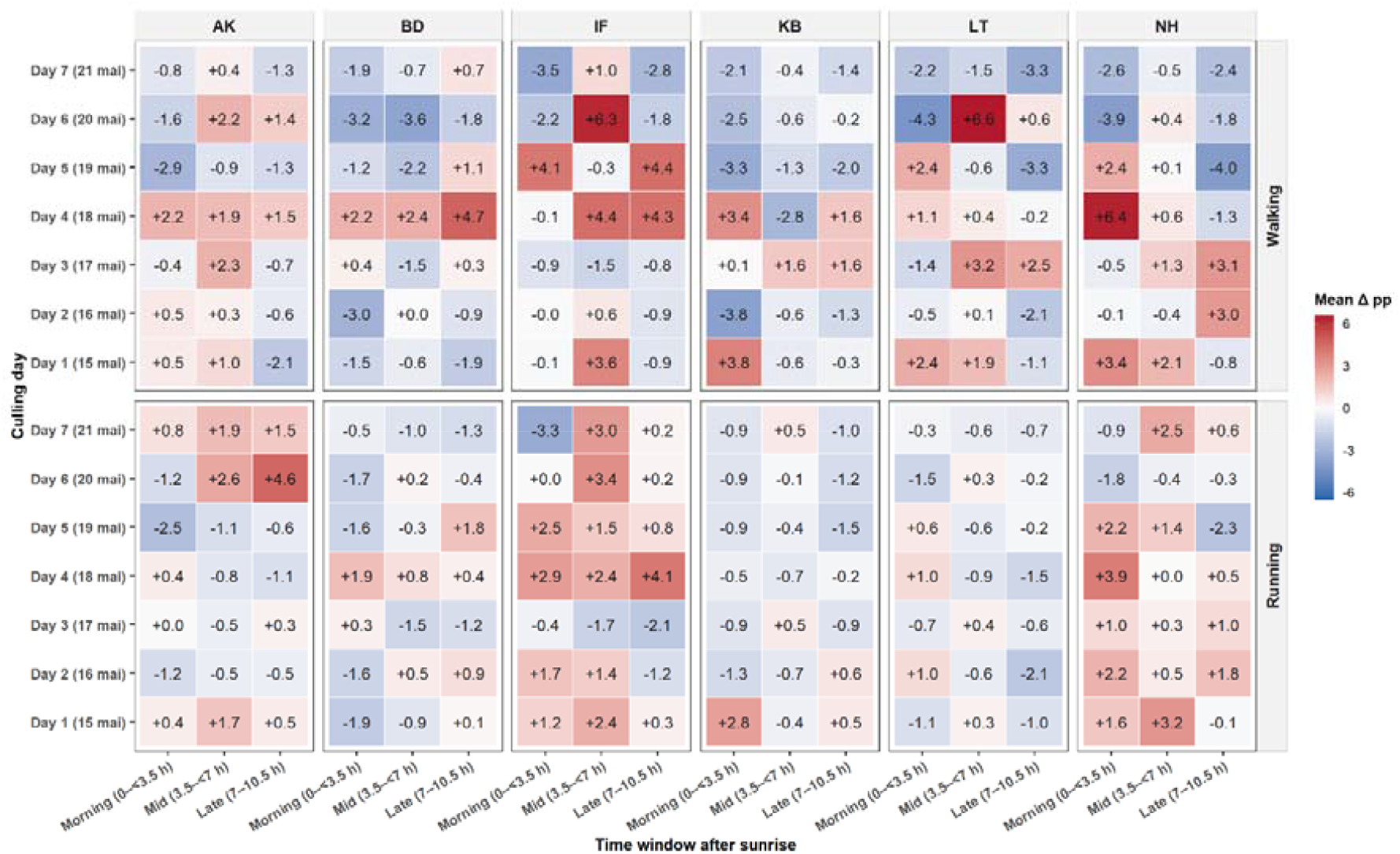
Acute within-day deviations from baseline during the culling period. Heatmaps show mean percentage-point deviations from the group-specific baseline expected at the same time of day for walking and running during each culling day. Deviations are summarised within three periods after sunrise (morning, mid-day, and late day). Positive values indicate higher-than-expected activity relative to baseline, and negative values indicate lower-than-expected activity.

Walking showed the largest fine-scale deviations, with several group × day × time-window combinations reaching approximately +6 percentage points above baseline, particularly in IF, LT and NH (Fig. 6). Given baseline walking levels of approximately 4–6% in these groups (Table S5), such deviations could represent a doubling or more of expected walking activity at the corresponding time of day. These short-lived increases were much larger than the phase-averaged contrasts, which did not exceed approximately 1 percentage point across groups. Two temporal patterns were apparent. In some cases, positive deviations were spread across all three time windows, suggesting a more sustained increase in locomotion across the day, as observed in AK and BD on day 4. In other cases, deviations were concentrated within a single time window and accompanied by near-zero or negative deviations in adjacent windows, suggesting brief locomotor responses rather than sustained increases across the full day. Examples include IF and LT on day 6 and NH on day 4 (Fig. 6). The strongest deviations also tended to cluster across groups on particular culling days. Positive walking deviations were especially apparent on day 1 and between days 4 to 6. On day 6, two of the three strongest positive deviations occurred around midday in LT (+6.6 percentage points) and IF (+6.3 percentage points). Negative deviations also co-occurred on the same day, with two of the strongest morning reductions observed in LT (−4.3 percentage points) and NH (−3.9 percentage points; Fig. 6).

Running showed smaller absolute deviations, though standardised residuals were of comparable magnitude to those for walking given its lower baseline frequency of approximately 1% to 3%. Unlike walking, running showed more heterogeneity across groups than across days, with IF, NH, and to a lesser extent AK displaying most of the increases relative to their baseline (Fig. S6a). Deviations in running were moreover not consistently correlated with those in walking, indicating that the two behaviours could respond differently on the same day and within the same time window.

## Discussion

In this study, we investigated how a large-scale helicopter-assisted culling operation targeting ungulates impacted the behaviour and space use of vervet monkeys. Responses were heterogeneous across behaviours and groups, but several patterns nevertheless pointed to fear-consistent behavioural and spatial reorganisation. Foraging declined during the disturbance, whereas inactivity and social behaviour showed an increasing tendency. Spatial responses were expressed as adjustments within existing home ranges rather than large-scale displacement, while acute locomotor deviations only emerged at finer temporal resolution, together highlighting that responses varied across both spatial and temporal scales. That such patterns emerged in a non-target species highlights the potential for lethal anthropogenic disturbance to generate effects well beyond its intended scope.

The clearest behavioural response was a decline in foraging. This decrease began around the onset of culling and recovered partly afterwards, suggesting that it was more closely linked to the disturbance window than to a gradual background trend. In addition, circadian analyses revealed that this reduction was concentrated in the morning, when helicopter activity was reported to be the most intense. This pattern points toward a cue-driven response rather than a general seasonal decline in activity. Our results are therefore consistent with the risk allocation hypothesis (Lima & Bednekoff, 1999), suggesting that animals can partition risky activities to avoid periods of heightened perceived threat (Gaynor et al., 2018; Kohl et al., 2018). More broadly, the response of vervets is consistent with broader evidence that anthropogenic cues can induce strong behavioural shift in wildlife, as was previously shown in elk and pumas (Ciuti et al., 2012; Smith et al., 2017). Our study adds to this literature by showing that a lethal management operation can similarly reduce and restructure foraging activity in a non-target primate.

By contrast, eating showed weaker and less consistent alignment with the culling period, suggesting that the event did not simply reduce all feeding-related activity. This distinction is biologically plausible because the derived foraging category captured active food-searching involving movement through the environment, whereas eating reflected more stationary feeding that may occur from comparatively sheltered positions, such as trees or dense vegetation. Active food-searching may therefore have exposed individuals more directly to acoustic, aerial, or visual disturbance cues, making it more sensitive to perceived threat than food consumption itself.

Inactivity showed a more complex trajectory than foraging. Whereas foraging was most clearly reduced during the culling period, inactivity increased more gradually and remained more elevated after the operation. This pattern likely reflects both disturbance-related change and seasonal background effects. The study window coincided with the onset of winter, when cooler conditions and shorter days are expected to reduce activity in primates and other mammals (Brivio et al., 2016; McFarland et al., 2014). However, seasonal drift alone is unlikely to explain the full pattern. Mean temperature was lower during culling than afterwards (14.4°C vs. 15.4°C; Fig. S8), yet inactivity remained more elevated after the operation, indicating that the trajectory did not simply track temperature. In addition, inactivity began to increase around the start of the culling period and showed its clearest circadian deviation in the early morning, partly mirroring the morning reduction in foraging during the window of reported peak disturbance. This suggests that apparent inactivity reflected both a broader seasonal component and a disturbance-aligned shift during culling. Interpretation of this response is also constrained by the behavioural resolution of accelerometry: we cannot distinguish true rest from vigilance, which share similar low-movement postural signatures but have different ecological meanings. Within the ecology-of-fear framework, vigilance is expected to increase at the expense of foraging (Brown, 1999).

Part of the apparent inactivity response may therefore reflect heightened alertness rather than reduced arousal. Distinguishing between these possibilities matters because vigilance-like inactivity would imply a safety–resource trade-off, whereas true rest could also reflect seasonal or thermal constraints. Direct observation or physiological measures would be needed to test this interpretation further.

We similarly observed an increasing trend in social behaviour during the culling period. Although this increase was not statistically supported after correction, this may partly reflect the limited power of the phase contrast analyses inherent to an opportunistic design with several groups, few individuals, multiple behavioural tests, and a relatively short temporal window, rather than a true absence of effect. Supporting this interpretation, both population-and group-level trajectories showed a peak centred around the disturbance window followed by a rapid return to baseline, with an effect size proportionally stronger than that of foraging given the lower baseline occurrence of affiliative behaviours. The circadian analysis further suggested that social behaviour was most elevated in the morning compared to both the before and after phases, when the strongest foraging reduction was also observed. These patterns are consistent with the social buffering hypothesis, whereby affiliative interactions with conspecifics can attenuate physiological stress responses under challenging conditions (Sanchez et al., 2015; Young et al., 2014). However, the exact nature of this social response remains unresolved. After Hurricane Maria, rhesus macaques showed persistent increases in social connections, especially among previously peripheral individuals, suggesting that severe and prolonged environmental disruption can reshape social networks beyond the immediate stressor (Testard et al., 2021). By contrast, experimental work on vervets at our study site showed that social allocation can also change rapidly and reversibly when ecological constraints are manipulated (Gareta García et al., 2021). The short duration of the social peak in our study is more consistent with a temporary adjustment to a bounded disturbance than with a lasting social reorganisation. However, because accelerometry captures social time but not partner identity, proximity structure, or grooming direction, we cannot determine whether this increase reflected stronger investment in established partners, broader redistribution of affiliation and increased group cohesion, or passive aggregation under perceived threat. Pairing biologging with proximity sensors or direct observations would allow future work to test what kind of social buffering, if any, occurs during lethal management disturbance.

Locomotor responses were more difficult to detect at the daily scale, but the finer temporal analyses revealed clear short-lived deviations in walking and running. This contrast between weak phase-level effects and stronger acute responses is informative. It suggests that locomotor reactions to culling were brief, unevenly distributed across days, and therefore easily diluted when averaged across the full disturbance period. This is consistent with the idea that the temporal scale of analysis must match the temporal grain of the behavioural process being studied (Gaillard et al., 2010). Because antipredator behaviours are often transient and context dependent, relying solely on daily mean activity can mask finer-scale behavioural plasticity and subsequent temporal reorganisation (Brivio et al., 2016, 2024). Critically, the strongest locomotor deviations were not randomly distributed across the culling period. Some days concentrated large deviations across several groups, suggesting that particular windows may have involved more intense or more proximal disturbance, such as helicopter overflights or nearby shooting episodes. Notably, on the sixth day of culling, LT and IF showed two of the strongest positive walking deviations, whereas BD and NH showed two of the strongest negative deviations on the same day (Fig. 6). This contrast suggests that groups may have experienced a common external trigger but may have expressed different behavioural responses depending on local context. Without continuous accelerometry, these responses would probably have proved difficult to detect. Instead, the fine-scale analyses suggest that the same culling operation produced both sustained changes in behavioural time allocation and brief event-driven locomotor deviations, in line with short-lived antipredator responses to acute threat cues detected in other species (Clinchy et al., 2013).

The heterogeneity in acute locomotor responses was mirrored by heterogeneous spatial responses. Across groups, however, these responses consistently reflected adjustments within existing ranges rather than larger-scale displacement (Fig. 1). No group appeared to abandon its existing home range, which is consistent with the high costs that range abandonment can impose on group-living primates (Willems & Hill, 2009). KB provided the clearest refuge-oriented response, combining a directional reduction in space use with a significant increase in dense vegetation during culling. This combination most closely matches the prediction that animals should concentrate activity in safer or more sheltered areas under elevated perceived risk (Gaynor et al., 2021; Laundré et al., 2010). Other groups showed different spatial strategies. AK and NH showed evidence of spatial contraction without a corresponding shift toward denser vegetation, and both were associated with relatively strong running deviations. IF, for which MCP-based inference was unavailable because no collared female was present, also showed marked locomotor responses in running but no comparable evidence for contraction.

Several non-exclusive processes may account for this divergence. First, exploratory NDVI summaries of daily ranging areas suggested that these groups, particularly IF, occupied comparatively more open areas, supporting the interpretation that dense vegetation may not have been equally available, accessible, or effective as a refuge habitat across home ranges (Fig. S9). Second, more proximal or intense disturbance may have made concealment less viable than repeated short-distance flight. In this sense, the same underlying response to perceived threat may have produced different behavioural and spatial outcomes depending on local habitat structure and exposure. Finally, prior experience with human-associated threat may also have contributed to group-level heterogeneity. In primates, antipredator responses can vary with local hunting pressure: Diana monkeys (*Cercopithecus diana*) exposed to hunting show more cryptic responses to humans and finer discrimination of human threat cues (Bshary, 2001), while predator-relevant information can also spread socially through alarm calls and learned avoidance responses (Griffin, 2004; León et al., 2022). KB provides a plausible example of how such prior exposure may shape group-level responses. This group occupied the area closest to a village with regular human activity and reported poaching incidents, and field assistants consistently reported that this group often moved rapidly into arboreal cover in response to nearby shooting or dogs while remaining silent, whereas other groups more often fled and gave alarm calls (unpublished data). Although this mechanism remains untested, repeated exposure to poaching and related anthropogenic threats may have further shaped a group-level concealment response in KB. More broadly, the diversity of responses observed here likely reflects a combination of habitat constraints, differential exposure, and group-specific histories of threat highlighting the value of moving beyond population-level summaries to consider how different groups, and potentially individuals, react to the same lethal management operation.

Nevertheless, several limitations bear on how these results should be interpreted. The primary constraint is the absence of independent exposure data. Because helicopter activity, shooting locations, and carcass processing were not quantified in space and time, differences among groups cannot be directly attributed to variation in the type, intensity, or magnitude of exposure. This limits inferences for the observed heterogeneity but does not undermine the conclusion that behaviour was altered around the culling event. Second, because biologging data were available for only one year, the study lacked both an unaffected comparison group and equivalent data from the same calendar period in other years. To limit seasonal and phenological confounding, we restricted the main before and after phases to approximately two-week windows around the culling period, but co-occurring background trends still cannot be formally excluded, particularly for gradual responses such as inactivity. This is why temporal structure was central to interpretation: responses centred on the culling window, concentrated during reported peak-disturbance hours, or visible only at fine temporal scales provide stronger evidence for a disturbance-related process than broad before-after differences alone.

Furthermore, spatial analyses were constrained by sampling resolution and the choice of space-use estimator. Daily path length estimates standardised to a 4-h GPS fix schedule likely underestimated tortuous or back-and-forth movements (Nathan et al., 2022). MCPs provided a simple and comparable measure of daily spatial extent, but they are sensitive to peripheral fixes and sampling effort and may overestimate space use by enclosing unused areas or conversely, underestimate it when sparse fixes fail to capture the full daily range (Burgman & Fox, 2003; Fieberg & Börger, 2012). Moreover, MCPs describe spatial extent rather than intensity of use, and therefore do not capture core areas or repeated use refuges, which may have captured complementary aspects of the response. However, given our limited number of fixes available, other utilization-based estimators proved too unstable for daily comparisons. Finally, because MCPs were derived from a single collared female per group-day, they may not fully represent group space use on days when group cohesion was potentially reduced.

More broadly, this study focused primarily on group-level responses and therefore leaves interindividual variation only partly resolved. Given the strong spatial cohesion of vervet groups (Noë & Laporte, 2014; Struhsaker, 1967), this limitation is unlikely to strongly affect broad spatial inference, but it may be more relevant for behavioural or physiological responses to disturbance. Even within cohesive groups, individuals may differ in exposure to threat according to sex, social position, and within-group spatial positioning (Rhine et al., 1981). In vervets, females form the stable social core of the group, whereas adult males often occupy more exposed or leading positions during risky group progressions (Isbell et al., 1993; Tankink et al., 2025, preprint). Individual variation may therefore be expressed more through vigilance, locomotion, affiliation, or stress physiology than through major differences in ranging. Exploratory analyses of sex differences provided no clear evidence for sex-specific behavioural responses, and models including sex interactions were not retained. Individual-level phase contrasts were also generally consistent in direction within groups, especially for foraging, although some variation was visible across behaviours (Fig. S8).

Building on recent calls to evaluate the collateral effects of lethal management beyond target species and demographic groups (Mysterud, 2026), our study shows that such effects can extend to a non-target primate and involve multiple dimensions of behaviour and space use. Existing evidence remains concentrated on birds, while mammalian studies have mostly focused on ungulates and on spatial or temporal avoidance responses (Mysterud, 2026). Our study broadens this perspective by showing that culling was associated not only with within-range spatial adjustment, but also with changes in behavioural time allocation, circadian organisation and acute locomotor responses.

The marked variation among groups further suggests that collateral effects are unlikely to be uniform across a population. Responses were likely shaped by local habitat structure, uneven exposure, and group-specific context, with some groups showing stronger spatial contraction or shifts toward denser vegetation, and others showing more pronounced locomotor or behavioural changes. This heterogeneity matters because population-level summaries can mask biologically relevant within-population variation and may therefore underestimate disturbance effects when responses differ among groups, individuals, or behavioural dimensions (Harding et al., 2019).

Whether the responses documented here carried physiological or fitness costs remains unknown. However, reductions in foraging, changes in activity timing, altered social behaviour, and repeated acute locomotor responses could become costly if similar disturbances are frequent or prolonged. This possibility is supported by evidence that hunting-related disturbance can elevate glucocorticoids in wild primates (Kaisin et al., 2021), and that sustained non-lethal fear effects can reduce reproduction and survival at scales comparable to direct predation (Zanette & Clinchy, 2019). Physiological responses may also occur even when overt behavioural responses are weak or absent, meaning that behaviour-only assessments may underestimate the full cost of disturbance (Ellenberg et al., 2013; Wascher, 2021). Our results therefore suggest that assessments focused only on mortality, displacement, or target-species responses are likely to underestimate the broader ecological footprint of lethal management. Continuous biologging provides a practical way to detect these effects as they unfold, especially when paired with direct measures of exposure, physiological stress, and longer-term demographic monitoring.

## Author contributions

L.B.: Conceptualization, Methodology, Software, Validation, Formal Analysis, Investigation, Data Curation, Writing - Original Draft, Writing - Review & Editing, Visualization; J.A.T.: Conceptualization, Methodology, Validation, Investigation, Data Curation, Writing - Review & Editing; Visualization E.vd.W.: Conceptualization, Funding Acquisition, Resources, Supervision, Writing - Review & Editing

## Acknowledgements

We first and foremost want to thank all people going into the field to collect demographic and behavioural data during the culling: Anumita Samanta, Eleanor Matthews, Jérémy Perez, Lou Coudurier, Marisa Nicol, Marta Lago, Noah de Jong, Nokubonga Dlamini, Omar el Mecky, Sarah Mendes and Veronicca Khosana. We are very grateful for your hard work and preservation during that difficult week. We thank the van der Walt family for permission to conduct this study on their land. We are deeply grateful for all on-site managers that contributed to the collaring that provided the GPS and accelerometry data, including Michael Henshall, Siboniso Thela and Zonke Mbutho. We thank Michel Halbwax for his veterinary assistance during collar deployment and removal. We are also grateful to Ebi Anthony George and Elham Nourani for valuable discussions and comments.

## Data availability

The data and R code supporting this study will be archived on Zenodo and made publicly available upon publication.

## Supporting Information

This manuscript contains one additional supplementary file containing Supplementary Methods (SM1-SM5); Supplementary Tables S1-S5; Supplementary Figures S1-S9.

## Declaration of generative AI and AI-assisted technologies

During the preparation of this work the authors used Claude in order to assist in refining code associated with this manuscript and in language editing. After using this tool, the authors reviewed and edited the content as needed and take full responsibility for the content of the published article.

## References

Beale, C. M., & Monaghan, P. (2004). Human disturbance: people as predation-free predators? Journal of Applied Ecology, 41(2), 335–343. 10.1111/J.0021-8901.2004.00900.X

Benjamini, Y., & Hochberg, Y. (1995). Controlling the False Discovery Rate: A Practical and Powerful Approach to Multiple Testing. Journal of the Royal Statistical Society: Series B (Methodological), 57(1), 289–300. 10.1111/J.2517-6161.1995.TB02031.X

Berger-Tal, O., Polak, T., Oron, A., Lubin, Y., Kotler, B. P., & Saltz, D. (2011). Integrating animal behavior and conservation biology: a conceptual framework. Behavioral Ecology, 22(2), 236–239. 10.1093/BEHECO/ARQ224

Brivio, F., Apollonio, M., Anderwald, P., Filli, F., Bassano, B., Bertolucci, C., & Grignolio, S. (2024). Seeking temporal refugia to heat stress: Increasing nocturnal activity despite predation risk. Proceedings of the Royal Society B: Biological Sciences, 291(2015). 10.1098/RSPB.2023.1587/104328

Brivio, F., Bertolucci, C., Tettamanti, F., Filli, F., Apollonio, M., & Grignolio, S. (2016). The weather dictates the rhythms: Alpine chamois activity is well adapted to ecological conditions. Behavioral Ecology and Sociobiology, 70(8), 1291–1304. 10.1007/S00265-016-2137-8/FIGURES/5

Brooks, M. E., Kristensen, K., van Benthem, K. J., Magnusson, A., Berg, C. W., Nielsen, A., Skaug, H. J., Mächler, M., & Bolker, B. M. (2017). glmmTMB balances speed and flexibility among packages for zero-inflated generalized linear mixed modeling. R Journal, 9(2), 378–400. 10.32614/rj-2017-066

Brown, C. L., Smith, J. B., Wisdom, M. J., Rowland, M. M., Spitz, D. B., & Clark, D. A. (2020). Evaluating Indirect Effects of Hunting on Mule Deer Spatial Behavior. Journal of Wildlife Management, 84(7), 1246–1255. 10.1002/jwmg.21916

Brown, J. S. (1999). Vigilance, patch use and habitat selection: Foraging under predation risk. Evolutionary Ecology Research, 1(1), 49–71.

Brun, L., Rothrock, J., van de Waal, E., & George, E. A. (2026). Classifier architecture and data preprocessing jointly shape accelerometer-based behavioural inference. In bioRxiv (p. 2026.02.16.706143). Cold Spring Harbor Laboratory. 10.64898/2026.02.16.706143

Bshary, R. (2001). Diana monkeys, Cercopithecus diana, adjust their anti-predator response behaviour to human hunting strategies. Behavioral Ecology and Sociobiology, 50(3), 251–256. 10.1007/S002650100354/METRICS

Burgman, M. A., & Fox, J. C. (2003). Bias in species range estimates from minimum convex polygons: implications for conservation and options for improved planning. Animal Conservation Forum, 6(1), 19–28. 10.1017/S1367943003003044

Burnham, K. P., & Anderson, D. R. (2002). Model selection and multimodel inference: a practical information-theoretic approach. Springer.

Chalkowski, K., Snow, N. P., Feuka, A. B., Leland, B. R., VerCauteren, K. C., Miller, R. S., & Pepin, K. M. (2025). Wildlife movement and contact responses to intensive culling: implications for disease control. 10.1101/2025.06.13.659516

Ciuti, S., Northrup, J. M., Muhly, T. B., Simi, S., Musiani, M., Pitt, J. A., & Boyce, M. S. (2012). Effects of Humans on Behaviour of Wildlife Exceed Those of Natural Predators in a Landscape of Fear. PLOS ONE, 7(11), e50611. 10.1371/JOURNAL.PONE.0050611

Clinchy, M., Sheriff, M. J., & Zanette, L. Y. (2013). Predator-induced stress and the ecology of fear. Functional Ecology, 27(1), 56–65. 10.1111/1365-2435.12007

Darimont, C. T., Fox, C. H., Bryan, H. M., & Reimchen, T. E. (2015). The unique ecology of human predators. Science, 349(6250), 858–860. 10.1126/science.aac4249

Díaz, S., Settele, J., Brondízio, E. S., Ngo, H. T., Agard, J., Arneth, A., Balvanera, P., Brauman, K. A., Butchart, S. H. M., Chan, K. M. A., Lucas, A. G., Ichii, K., Liu, J., Subramanian, S. M., Midgley, G. F., Miloslavich, P., Molnár, Z., Obura, D., Pfaff, A., … Zayas, C. N. (2019). Pervasive human-driven decline of life on Earth points to the need for transformative change. In Science (Vol. 366, Issue 6471). American Association for the Advancement of Science. 10.1126/science.aax3100

Ellenberg, U., Mattern, T., & Seddon, P. J. (2013). Heart rate responses provide an objective evaluation of human disturbance stimuli in breeding birds. Conservation Physiology, 1(1). 10.1093/conphys/cot013

Erbe, C., Dent, M. L., Gannon, W. L., McCauley, R. D., Römer, H., Southall, B. L., Stansbury, A. L., Stoeger, A. S., & Thomas, J. A. (2022). The effects of noise on animals. In Exploring Animal Behavior Through Sound: Volume 1: Methods (pp. 459–506). Springer International Publishing. 10.1007/978-3-030-97540-1_13

Fieberg, J., & Börger, L. (2012). Could you please phrase “home range” as a question? Journal of Mammalogy, 93(4), 890–902. 10.1644/11-MAMM-S-172.1

Frid, A., & Dill, L. (2002). Human-caused disturbance stimuli as a form of predation risk. Ecology and Society, 6(1). 10.5751/es-00404-060111

Gaillard, J. M., Hebblewhite, M., Loison, A., Fuller, M., Powell, R., Basille, M., & Van Moorter, B. (2010). Habitat–performance relationships: finding the right metric at a given spatial scale. Philosophical Transactions of the Royal Society B: Biological Sciences, 365(1550), 2255–2265. 10.1098/RSTB.2010.0085

Gareta García, M., Farine, D. R., Brachotte, C., Borgeaud, C., & Bshary, R. (2021). Wild female vervet monkeys change grooming patterns and partners when freed from feeding constraints. Animal Behaviour, 181, 117–136. 10.1016/j.anbehav.2021.08.027

Gaynor, K. M., Cherry, M. J., Gilbert, S. L., Kohl, M. T., Larson, C. L., Newsome, T. M., Prugh, L. R., Suraci, J. P., Young, J. K., & Smith, J. A. (2021). An applied ecology of fear framework: linking theory to conservation practice. In Animal Conservation (Vol. 24, Issue 3, pp. 308–321). John Wiley & Sons, Ltd. 10.1111/acv.12629

Gaynor, K. M., Hojnowski, C. E., Carter, N. H., & Brashares, J. S. (2018). The influence of human disturbance on wildlife nocturnality. Science, 360(6394), 1232–1235. 10.1126/SCIENCE.AAR7121/SUPPL_FILE/AAR7121-GAYNOR-SM.PDF

Griffin, A. S. (2004). Social learning about predators: A review and prospectus. Learning and Behavior, 32(1), 131–140. 10.3758/BF03196014/METRICS

Grignolio, S., Merli, E., Bongi, P., Ciuti, S., & Apollonio, M. (2011). Effects of hunting with hounds on a non-target species living on the edge of a protected area. Biological Conservation, 144(1), 641–649. 10.1016/j.biocon.2010.10.022

Ham, C., Donnelly, C. A., Astley, K. L., Jackson, S. Y. B., & Woodroffe, R. (2019). Effect of culling on individual badger Meles meles behaviour: Potential implications for bovine tuberculosis transmission. Journal of Applied Ecology, 56(11), 2390–2399. 10.1111/1365-2664.13512

Harding, H. R., Gordon, T. A. C., Eastcott, E., Simpson, S. D., & Radford, A. N. (2019). Causes and consequences of intraspecific variation in animal responses to anthropogenic noise. In Behavioral Ecology (Vol. 30, Issue 6, pp. 1501–1511). Oxford Academic. 10.1093/beheco/arz114

Hartig, F. (2024). DHARMa - Residual Diagnostics for HierARchical Models. In R package version 0.4.7 (pp. 1–65). Comprehensive R Archive Network (CRAN). 10.32614/CRAN.PACKAGE.DHARMA

Isbell, L. A., Cheney, D. L., & Seyfarth, R. M. (1993). Are immigrant vervet monkeys, Cercopithecus aethiops, at greater risk of mortality than residents? Animal Behaviour, 45(4), 729–734. 10.1006/anbe.1993.1087

Kaisin, O., Fuzessy, L., Poncin, P., Brotcorne, F., & Culot, L. (2021). A meta-analysis of anthropogenic impacts on physiological stress in wild primates. Conservation Biology, 35(1), 101–114. 10.1111/COBI.13656

Katzner, T. E., & Arlettaz, R. (2020). Evaluating Contributions of Recent Tracking-Based Animal Movement Ecology to Conservation Management. Frontiers in Ecology and Evolution, 7. 10.3389/FEVO.2019.00519/FULL

Kohl, M. T., Stahler, D. R., Metz, M. C., Forester, J. D., Kauffman, M. J., Varley, N., White, P. J., Smith, D. W., & MacNulty, D. R. (2018). Diel predator activity drives a dynamic landscape of fear. Ecological Monographs, 88(4), 638–652. 10.1002/ECM.1313

Laundré, J. W., Hernández, L., & Ripple, W. J. (2010). The Landscape of Fear: Ecological Implications of Being Afraid. The Open Ecology Journal, 3, 1–7.

Lenth, R. V, & Piaskowski, J. (2025). emmeans: Estimated Marginal Means, aka Least-Squares Means.

León, J., Thiriau, C., Bodin, C., Crockford, C., & Zuberbühler, K. (2022). Acquisition of predator knowledge from alarm calls via one-trial social learning in monkeys. IScience, 25(9), 104853. 10.1016/j.isci.2022.104853

Lima, S. L., & Bednekoff, P. A. (1999). Temporal variation in danger drives antipredator behavior: The predation risk allocation hypothesis. American Naturalist, 153(6), 649–659. 10.1086/303202

Marchowski, D., Jurszo, R., Stańczak, P., Jasiński, M., & Guentzel, S. (2025). Non-selective waterbird hunting in a Natura 2000 site results in killing of protected species: A case study from western Poland. Avian Research, 16(3), 100276. 10.1016/j.avrs.2025.100276

McFarland, R., Barrett, L., Boner, R., Freeman, N. J., & Henzi, S. P. (2014). Behavioral flexibility of vervet monkeys in response to climatic and social variability. American Journal of Physical Anthropology, 154(3), 357–364. 10.1002/AJPA.22518

Mucina, L., & Rutherford, M. C. (2006). The vegetation of South Africa, Lesotho and Swaziland. The Vegetation of South Africa, Lesotho and Swaziland.

Mysterud, A. (2026). Collateral damage: hunting effects on non-target species and groups. In European Journal of Wildlife Research (Vol. 72, Issue 2, pp. 30-). Springer Science and Business Media Deutschland GmbH. 10.1007/s10344-026-02071-1

Nathan, R., Monk, C. T., Arlinghaus, R., Adam, T., Alós, J., Assaf, M., Baktoft, H., Beardsworth, C. E., Bertram, M. G., Bijleveld, A. I., Brodin, T., Brooks, J. L., Campos-Candela, A., Cooke, S. J., Gjelland, K., Gupte, P. R., Harel, R., Hellström, G., Jeltsch, F., … Jarić, I. (2022). Big-data approaches lead to an increased understanding of the ecology of animal movement. In Science (Vol. 375, Issue 6582). American Association for the Advancement of Science. 10.1126/science.abg1780

Nathan, R., Spiegel, O., Fortmann-Roe, S., Harel, R., Wikelski, M., & Getz, W. M. (2012). Using tri-axial acceleration data to identify behavioral modes of free-ranging animals: general concepts and tools illustrated for griffon vultures. Journal of Experimental Biology, 215(6), 986–996. 10.1242/JEB.058602

Noë, R., & Laporte, M. (2014). Socio-spatial cognition in vervet monkeys. Animal Cognition, 17(3), 597–607. 10.1007/s10071-013-0690-3

Ozsanlav-Harris, L., McIntosh, A. L. S., Griffin, L. R., Hilton, G. M., Cao, L., Shaw, J. M., & Bearhop, S. (2024). Contrasting effects of shooting disturbance on the movement and behavior of sympatric wildfowl species. Ecological Applications, 34(8), e3032. 10.1002/eap.3032

Pecorella, I., Ferretti, F., Sforzi, A., & Macchi, E. (2016). Effects of culling on vigilance behaviour and endogenous stress response of female fallow deer. Wildlife Research, 43(3), 189–196. 10.1071/WR15118

Peters, A., Smith, A. F., Henrich, M., Dormann, C. F., & Heurich, M. (2025). Temporal displacement of the mammal community in a protected area due to hunting and recreational activities. Ecological ApplicationsÖ: A Publication of the Ecological Society of America, 35(7), e70118. 10.1002/eap.70118

Pirotta, E., Booth, C. G., Costa, D. P., Fleishman, E., Kraus, S. D., Lusseau, D., Moretti, D., New, L. F., Schick, R. S., Schwarz, L. K., Simmons, S. E., Thomas, L., Tyack, P. L., Weise, M. J., Wells, R. S., & Harwood, J. (2018). Understanding the population consequences of disturbance. In Ecology and Evolution (Vol. 8, Issue 19, pp. 9934–9946). John Wiley & Sons, Ltd. 10.1002/ece3.4458

Rhine, R. J., Forthman, D. L., Stillwellß Barnes, R., Westlund, B. J., & Westlund, H. D. (1981). Movement patterns of yellow baboons (Papio cynocephalus): Sex differences in juvenile development toward adult patterns. American Journal of Physical Anthropology, 55(4), 473–484. 10.1002/ajpa.1330550408

Sanchez, M. M., Mccormack, K. M., & Howell, B. R. (2015). Social Neuroscience Social buffering of stress responses in nonhuman primates: Maternal regulation of the development of emotional regulatory brain circuits. 10.1080/17470919.2015.1087426

Semple, S., Harrison, C., & Lehmann, J. (2013). Grooming and Anxiety in Barbary Macaques. Ethology, 119(9), 779–785. 10.1111/ETH.12119

Seyfarth, R. M., Cheney, D. L., & Marler, P. (1980). Vervet monkey alarm calls: Semantic communication in a free-ranging primate. Animal Behaviour, 28(4), 1070–1094. 10.1016/S0003-3472(80)80097-2

Shepard, E. L. C., Wilson, R. P., Quintana, F., Laich, A. G., Liebsch, N., Albareda, D. A., Halsey, L. G., Gleiss, A., Morgan, D. T., Myers, A. E., Newman, C., & Macdonald, D. W. (2008). Identification of animal movement patterns using tri-axial accelerometry. Endangered Species Research, 10(1), 47–60. 10.3354/esr00084

Signer, J., Fieberg, J., & Avgar, T. (2019). Animal movement tools (amt): R package for managing tracking data and conducting habitat selection analyses. Ecology and Evolution, 9(2), 880–890. 10.1002/ece3.4823

Smith, J. A., Suraci, J. P., Clinchy, M., Crawford, A., Roberts, D., Zanette, L. Y., & Wilmers, C. C. (2017). Fear of the human ‘super predator’ reduces feeding time in large carnivores. *Proceedings*. Biological Sciences, 284(1857). 10.1098/RSPB.2017.0433

Struhsaker, T. T. (1967). Social Structure Among Vervet Monkeys (Cercopithecus Aethiops). Behaviour, 29(2–4), 83–121. 10.1163/156853967X00073

Tablado, Z., & Jenni, L. (2017). Determinants of uncertainty in wildlife responses to human disturbance. Biological Reviews, 92(1), 216–233. 10.1111/brv.12224

Tankink, J. A., van de Waal, E., Bshary, R., & van Schaik, C. P. (2025). Leadership under risk: male vervet monkey’s roles in group progression across high-risk terrain. In bioRxiv (p. 2025.11.21.689472). Cold Spring Harbor Laboratory. 10.1101/2025.11.21.689472

Testard, C., Larson, S. M., Watowich, M. M., Snyder-Mackler, N., Platt, M. L., Correspondence, L. J. N. B., Kaplinsky, C. H., Bernau, A., Faulder, M., Marshall, H. H., Lehmann, J., Ruiz-Lambides, A., Higham, J. P., Montague, M. J., & Brent, L. J. N. (2021). Rhesus macaques build new social connections after a natural disaster. Current Biology, 31. 10.1016/j.cub.2021.03.029

Tuomainen, U., & Candolin, U. (2011). Behavioural responses to human-induced environmental change. Biological Reviews, 86(3), 640–657. 10.1111/J.1469-185X.2010.00164.X

Venter, O., Sanderson, E. W., Magrach, A., Allan, J. R., Beher, J., Jones, K. R., Possingham, H. P., Laurance, W. F., Wood, P., Fekete, B. M., Levy, M. A., & Watson, J. E. M. (2016). Sixteen years of change in the global terrestrial human footprint and implications for biodiversity conservation. Nature Communications, 7(1), 12558-. 10.1038/ncomms12558

Wascher, C. A. F. (2021). Heart rate as a measure of emotional arousal in evolutionary biology. In Philosophical Transactions of the Royal Society B: Biological Sciences (Vol. 376, Issue 1831). Royal Society Publishing. 10.1098/rstb.2020.0479

Willems, E. P., & Hill, R. A. (2009). Predator-specific landscapes of fear and resource distribution: effects on spatial range use. Ecology, 90(2), 546–555. 10.1890/08-0765.1

Wong, B. B. M., & Candolin, U. (2015). Behavioral responses to changing environments. In Behavioral Ecology (Vol. 26, Issue 3, pp. 665–673). Oxford Academic. 10.1093/beheco/aru183

Wood, S. N. (2017). Generalized additive models: An introduction with R, second edition. Generalized Additive Models: An Introduction with R, Second Edition, 1–476. 10.1201/9781315370279/GENERALIZED-ADDITIVE-MODELS-SIMON-WOOD/RIGHTS-AND-PERMISSIONS

Woodroffe, R., Donnelly, C. A., Cox, D. R., Bourne, F. J., Cheeseman, C. L., Delahay, R. J., Gettinby, G., McInerney, J. P., & Morrison, W. I. (2006). Effects of culling on badger Meles meles spatial organization: Implications for the control of bovine tuberculosis. Journal of Applied Ecology, 43(1), 1–10. 10.1111/j.1365-2664.2005.01144.x

Young, C., Majolo, B., Heistermann, M., Schülke, O., & Ostner, J. (2014). Responses to social and environmental stress are attenuated by strong male bonds in wild macaques. Proceedings of the National Academy of Sciences of the United States of America, 111(51), 18195–18200. 10.1073/PNAS.1411450111/SUPPL_FILE/PNAS.201411450SI.PDF

Zanette, L. Y., & Clinchy, M. (2019). Ecology of fear. Current Biology, 29(9), R309–R313. 10.1016/j.cub.2019.02.042

