## Supplementary Material for "Collateral effects of lethal management operations: biologging reveals multiscale responses in non-target vervet monkeys"

### Supplementary Methods:

#### SM1. GPS preprocessing

GPS data were obtained for 21 collared individuals and downloaded from Movebank (Kays et al., 2022). Prior to analysis, locations were filtered to retain reliable fixes and avoid pseudo-replication. First, locations with manufacturer-estimated horizontal error >60 m were excluded. Second, because each scheduled GPS fix consisted of three positions recorded one second apart, each burst was reduced to a single representative location. Third, isolated positional artefacts were removed when they corresponded to implausible spikes away from the movement path followed by a rapid return. Candidate artefacts were identified using extreme step lengths (>3000 m) and turning angles (>100°), then confirmed by visual inspection of individual trajectories before removal. Finally, time lags between successive fixes were examined to identify irregular sampling intervals and data gaps.

#### SM2. Influence of sampling effort on movement and space-use metrics

GPS sampling frequency differed among collared individuals because females were programmed at 2-h intervals whereas males were programmed at 4-h intervals. Consequently, both trajectory resolution and the number of daily fixes varied across individuals, potentially affecting estimates of daily path length and daily minimum convex polygon (MCP) area. We therefore assessed the sensitivity of these spatial metrics to sampling effort.

Daily path length increased with the number of GPS fixes per day, but the relationship was sublinear on log-transformed axes (slope = 0.48, 95% CI: 0.40–0.57; Fig. SM2a). Because path length did not scale proportionally with sampling effort, it could not be corrected using a simple offset term. We therefore based the primary daily path length analysis on trajectories resampled to a common 4-h schedule, yielding a standardised sampling effort of 5–6 fixes per individual per day while maximising sample size.

Daily MCP area was also sensitive to sampling effort. In the female 2-h dataset, MCP estimates were highly variable at low fix numbers, particularly below seven fixes per day, but became more stable above this threshold (Fig. SM2b). MCP-based analyses were therefore restricted to females and to days with 7–12 fixes.

Sensitivity analyses showed that group-level during–before estimates for daily path length were broadly consistent across alternative sampling decisions (Fig. SM2c). The primary dataset, including all individuals resampled to 4-h intervals, produced broadly consistent effect directions and magnitudes to two alternative datasets: one including all individuals with 5–13 fixes per day and fix number included as a covariate, and one restricted to females with 7–12 fixes per day and fix number included as a covariate.

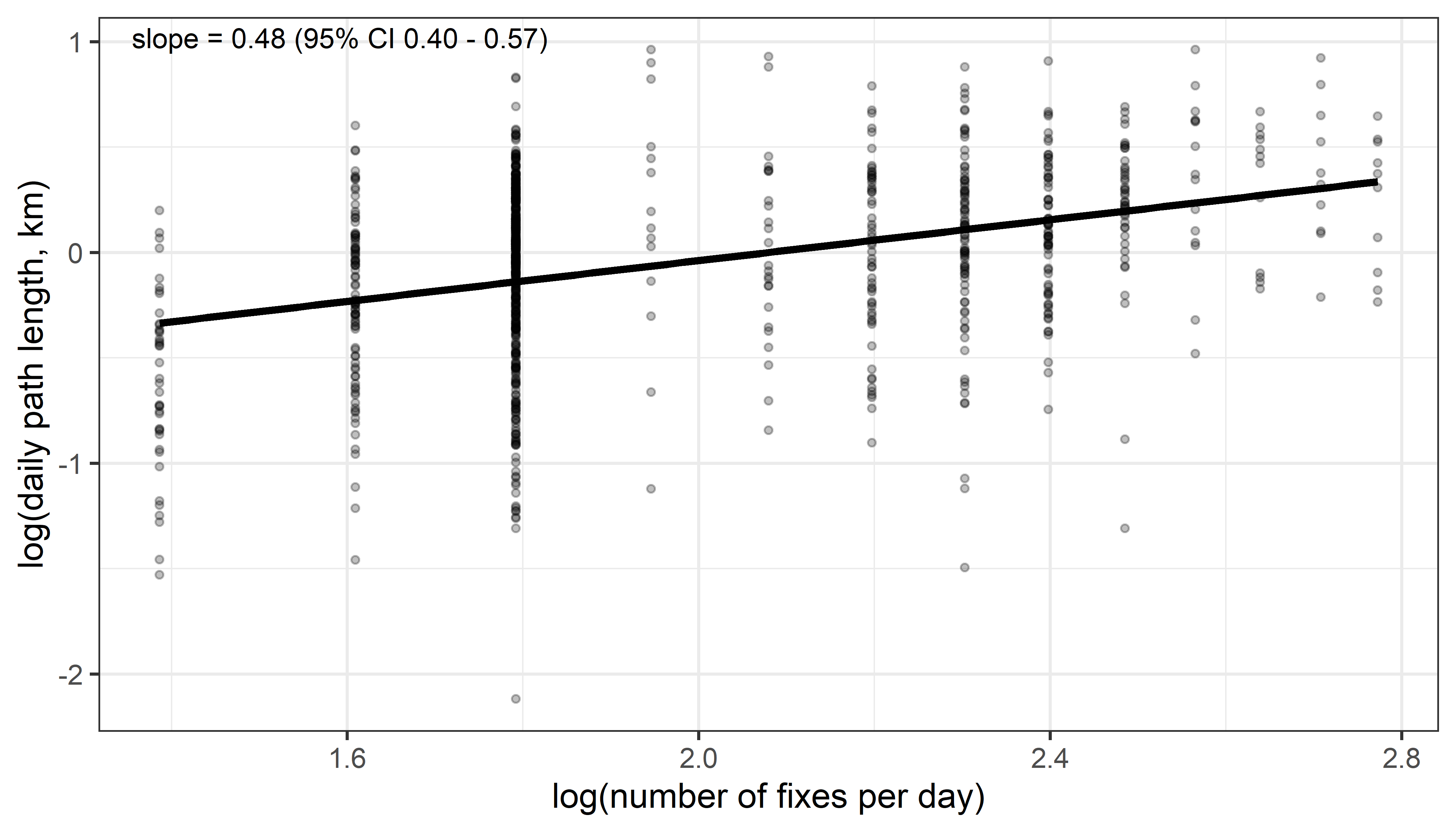

**Figure SM2a:** Relationship between daily path length and number of GPS fixes per day, shown on log-transformed axes and pooled across individuals and phases. The sublinear slope indicates that path length did not scale proportionally with fix number.

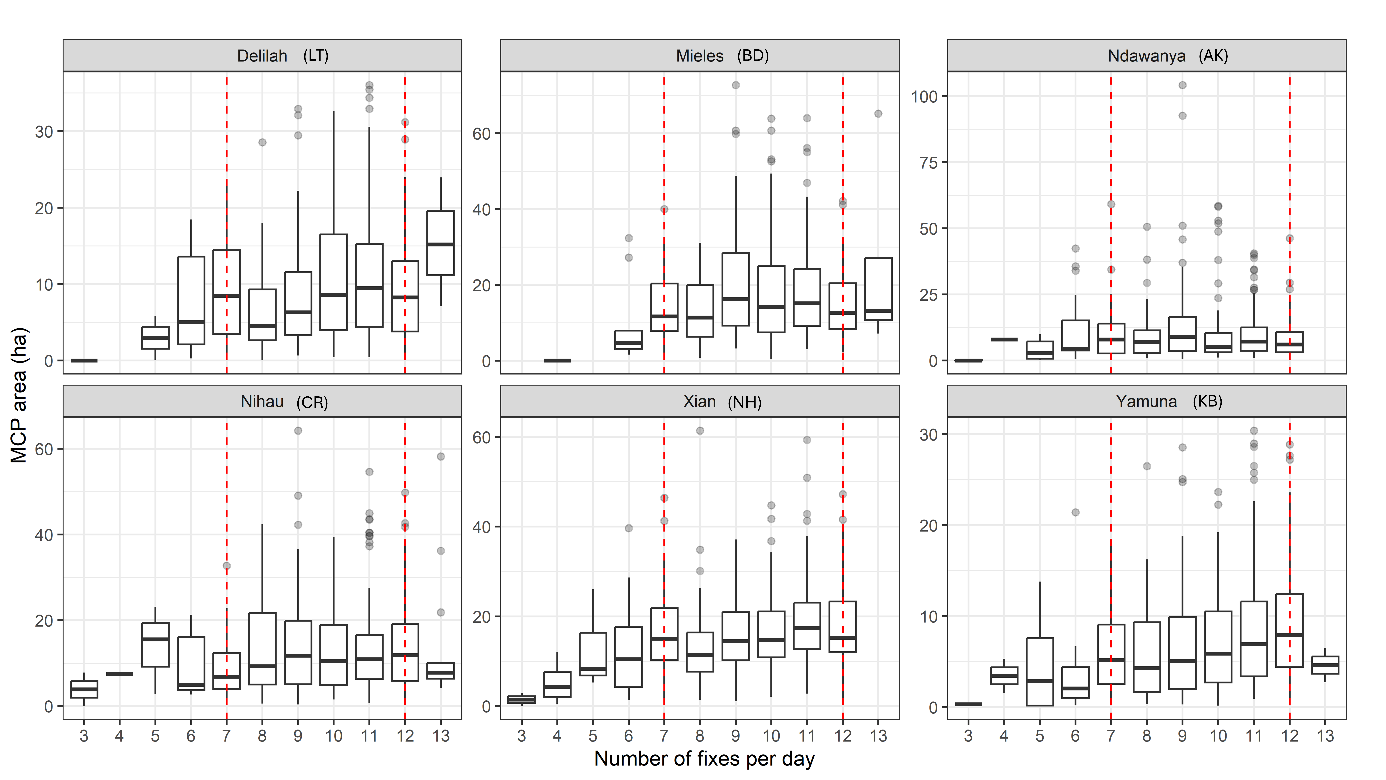

**Figure SM2b:** Distribution of daily MCP area across numbers of fixes per day for females sampled at 2-h resolution. Red dashed lines indicate the minimum and maximum fix thresholds retained for MCP analyses.

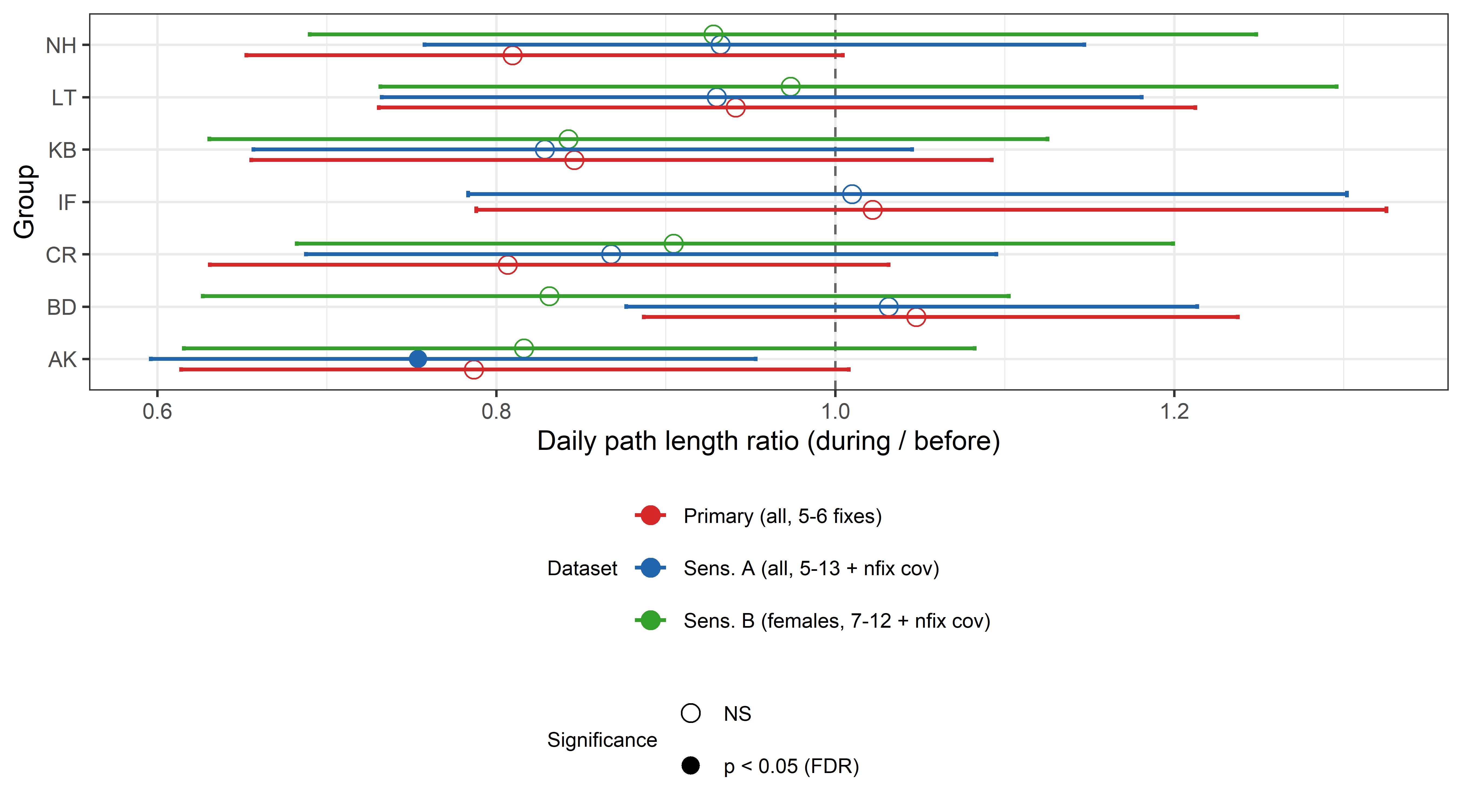

**Figure SM2c:** Sensitivity analysis of daily path length estimates across alternative datasets. Points show model-estimated group-level ratios for the during–before contrast, with horizontal lines indicating 95% confidence intervals. The vertical dashed line indicates no change. Filled symbols denote contrasts significant after FDR correction; open symbols indicate non-significant contrasts.

#### SM3. Rationale for daily space-use estimator selection

To identify a daily space-use metric that was robust and interpretable under our sampling regime, we compared 100% minimum convex polygons (MCPs), individual autocorrelated kernel density estimates (AKDEs), and group-level population kernel density estimates (PKDEs) using the ctmm workflow in R. Kernel-based estimators are attractive because they account for temporal autocorrelation and provide probabilistic utilisation distributions, but their performance was limited at the daily scale by short observation windows and low effective sample sizes. Daily AKDEs were frequently weakly supported after accounting for autocorrelation, resulting in poorly constrained and sometimes spatially inflated estimates. PKDEs were generally more stable because they pooled locations across individuals, but they often emphasised areas of repeated or shared use rather than the full spatial extent covered during a day. Because our objective was to compare daily group-level extent across disturbance phases, rather than estimate core-use distributions, MCPs provided the most transparent and comparable descriptor of daily space use. We therefore retained daily MCP area as the primary space-use metric.

To assess the reliability of daily AKDE and PKDE estimates, we examined the relationship between effective degrees of freedom for area (DOF) and relative confidence interval width on area (RCI = $A_{high}/A_{low}$). Across groups, low DOF values were consistently associated with large and highly variable RCI values, indicating weakly constrained area estimates. Estimates with DOF ≥ 5 generally showed smaller and more stable RCI values, so we used DOF ≥ 5 as the primary threshold for minimal reliability and RCI ≤ 5 as an additional conservative stability check (Fig. SM3a).

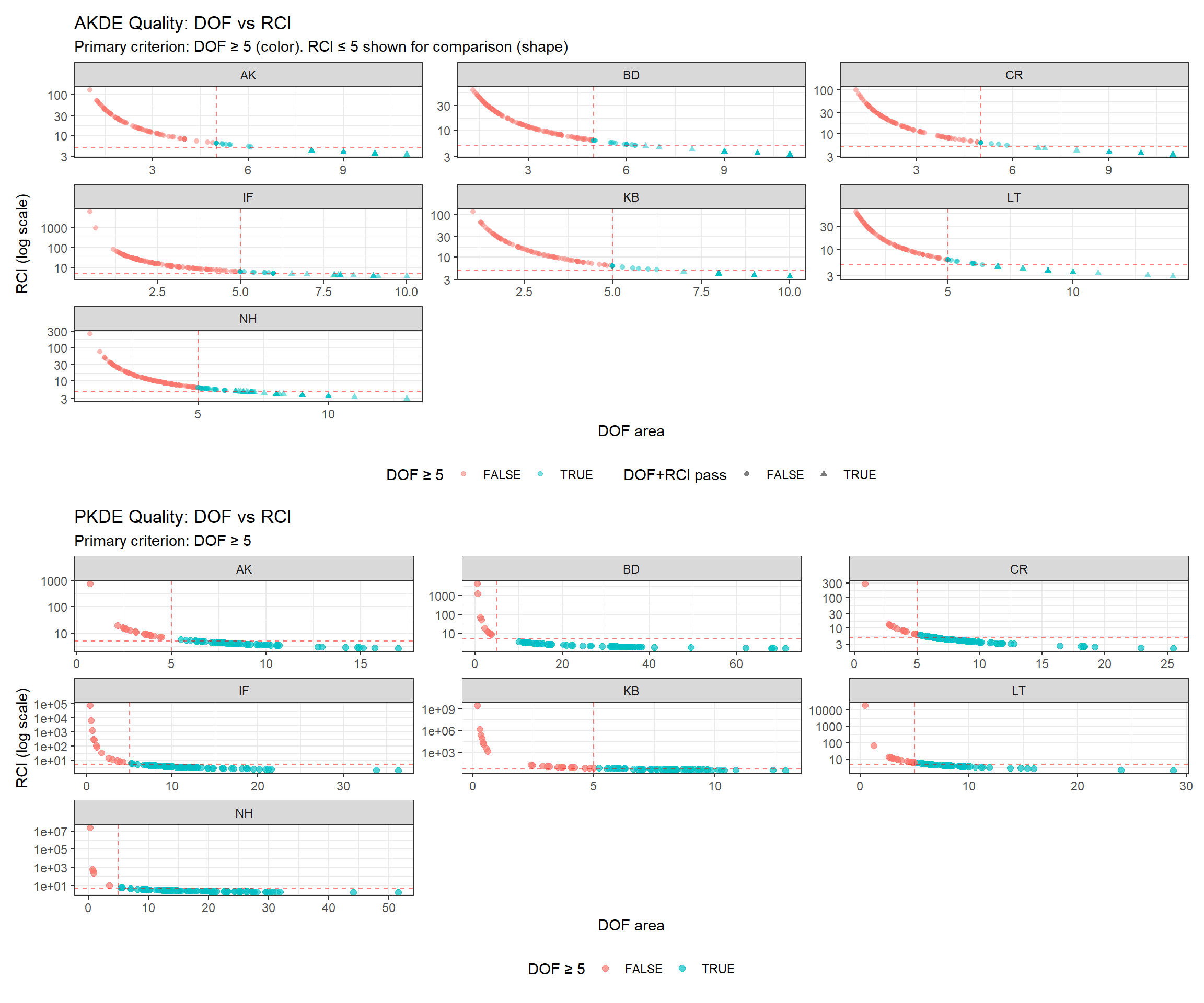

**Figure SM3a:** Quality control of daily AKDE and PKDE estimates based on effective degrees of freedom and uncertainty. Points show daily AKDEs and PKDEs, separated by group. The x-axis shows effective degrees of freedom for area (DOF), and the y-axis shows relative confidence interval width on area (RCI = upper/lower confidence limit; log scale). Red dashed lines indicate the thresholds DOF = 5 and RCI = 5. Low DOF values were associated with high uncertainty, while estimates above DOF ≈ 5 were generally more constrained.

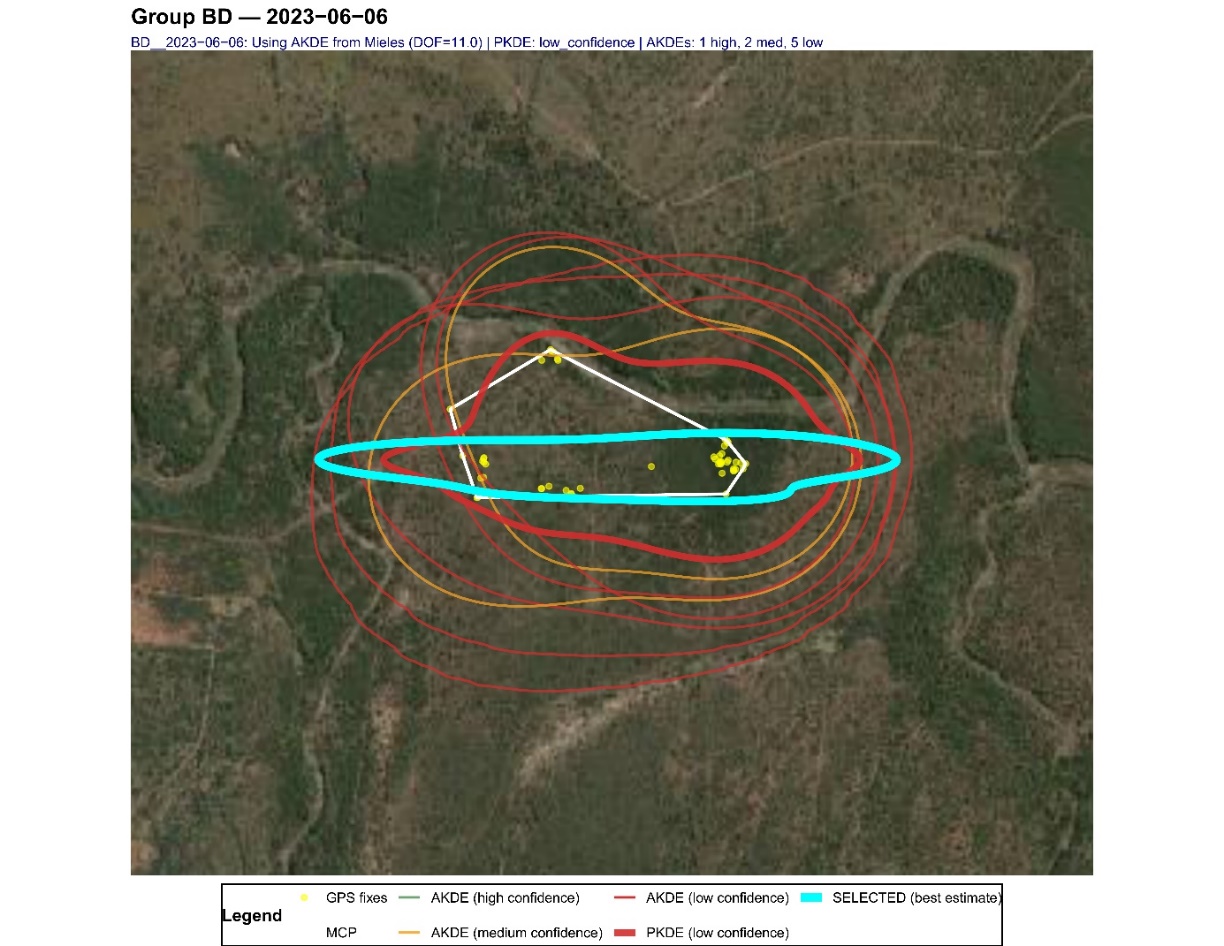

**b)**

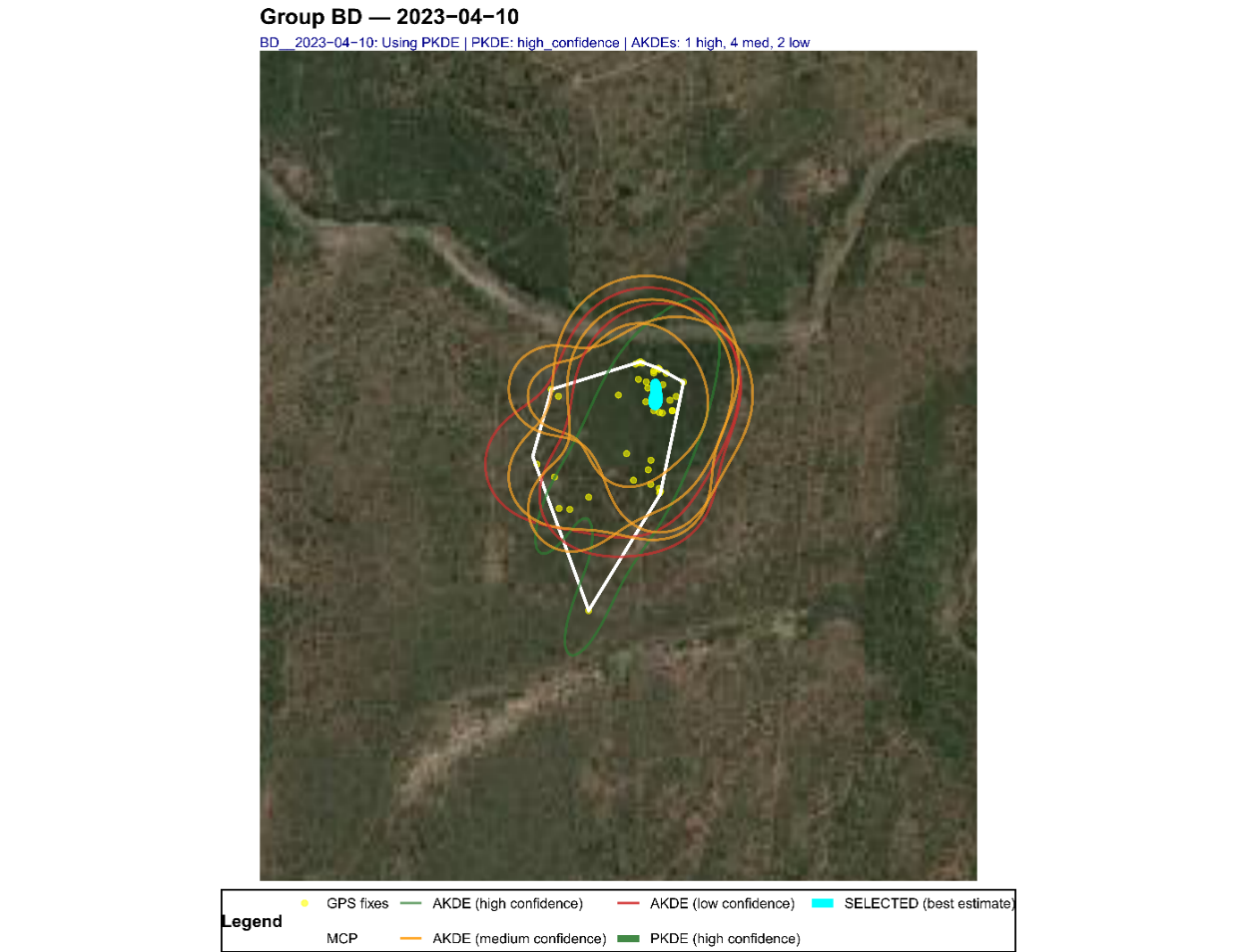

**c)**

**Figure SM3b–c:** Representative examples of daily space-use estimates for group BD on two days around the culling event. Yellow points indicate GPS fixes from collared individuals, white polygons indicate daily 100% MCPs, thin contours show individual AKDEs, and thick contours show group-level PKDEs. Contour colours indicate confidence classes for kernel estimates, with green, orange and red corresponding to high, medium and low confidence, respectively. Blue contours indicate the kernel estimate selected by the diagnostic AKDE/PKDE workflow as the best-supported kernel estimate for that group-day; this selection did not include MCPs and was used only to evaluate whether kernel-based estimators were suitable for daily inference. In the upper panel, AKDEs were often unstable or spatially inflated. In the lower panel, the selected PKDE captured repeated or shared use but did not represent the full daily extent indicated by the observed fixes and MCP. These examples illustrate why kernel-based estimators were not retained for final inference on daily spatial extent, and why daily MCP area was used as the primary space-use metric.

#### SM4. Habitat metrics derived from NDVI

Vegetation structure within each daily MCP was characterised using an NDVI-derived habitat classification. The original raster had a 10 × 10 m resolution (Fig. SM4a), and pixels were assigned to three categories: open (NDVI < 0.25), semi-open (0.25 ≤ NDVI < 0.33), and dense (NDVI ≥ 0.33; Fig. SM4b). To reduce isolated pixels and retain broader vegetation structure, we applied a 80 × 80 m focal smoothing filter to produce the final vegetation layer (Fig. SM4c). For each daily MCP, we calculated the proportional cover of each habitat class.

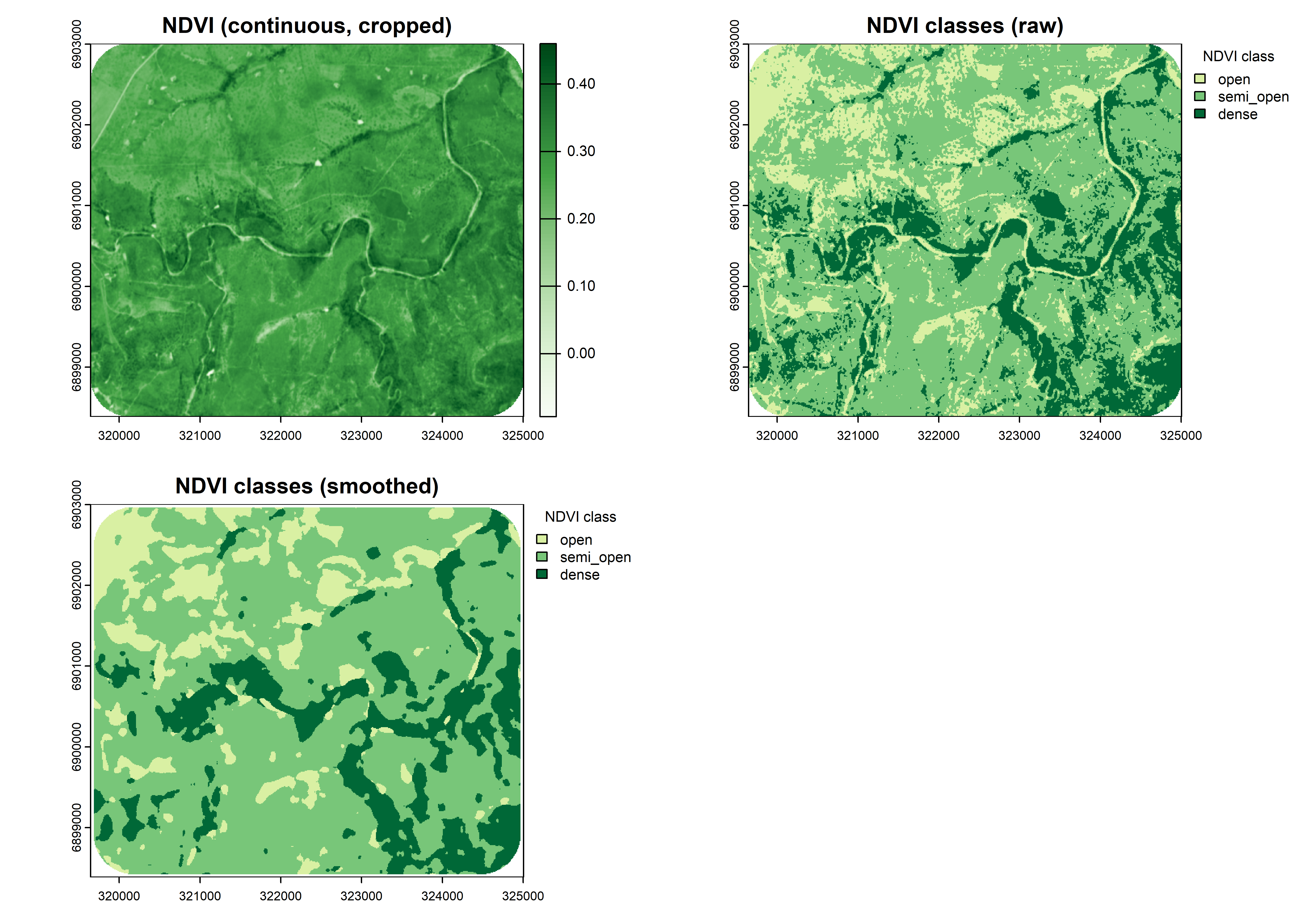

b)

c)

a)

**Figure SM4a-c.** NDVI-based habitat classification used to quantify vegetation structure within daily MCPs.
The upper-left panel shows the cropped continuous NDVI raster for the study area at 10 × 10 m resolution. The upper-right panel shows the raw classification of pixels into three habitat categories based on NDVI thresholds: open, NDVI < 0.25; semi-open, 0.25 ≤ NDVI < 0.33; and dense, NDVI ≥ 0.33. The lower panel shows the final smoothed classification after applying a 80 × 80 m focal filter to reduce isolated pixels and retain broader vegetation structure.

#### SM5. Accelerometer classification and derivation of final analytical behaviours

##### SM5.1 Sub-burst correction of isolated TabPFN predictions

Because TabPFN operates at the sub-burst scale (~3.4 s), it can capture within-burst behavioural variation, including short transitions and postural changes. However, this finer temporal resolution can also introduce isolated misclassifications, particularly among behaviours with similar low-movement signatures. To reduce this noise while preserving biologically meaningful within-burst variation, we applied a limited correction targeting clear 3+1 patterns, where three sub-bursts were assigned to the same behaviour and the remaining sub-burst to a posturally similar behaviour.

Overrides were applied only when three conditions were met: (i) three sub-bursts agreed on a dominant state, (ii) the minority sub-burst belonged to a predefined ambiguous low-movement pair, and (iii) HydraROCKET supported the dominant assignment at the whole-burst scale. Corrections were restricted to resting–sleeping, resting–grooming receiver, sleeping–grooming receiver, and eating–grooming actor pairs. Mixtures involving locomotor behaviours, such as walking or running, were not corrected because these were considered more likely to reflect genuine within-burst transitions. This procedure altered only a small fraction of sub-burst labels and had minimal effect on the overall behavioural composition (Fig. SM5a).

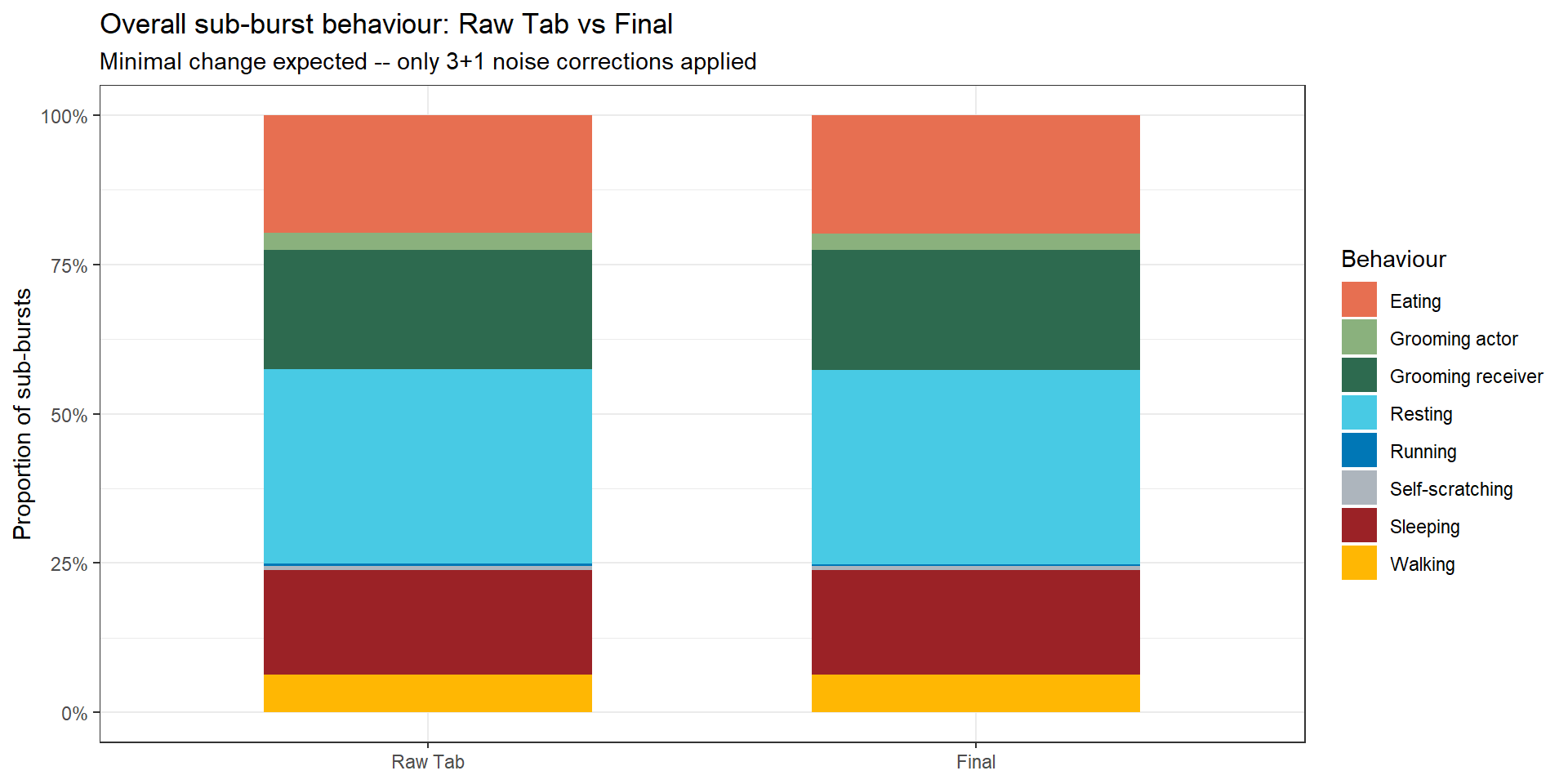

**Figure SM5a:** **Effect of sub-burst correction on behavioural composition.** Proportional distribution of TabPFN sub-burst predictions before and after the limited 3+1 correction procedure. Corrections were restricted to predefined ambiguous postural pairs and applied only when supported by the whole-burst HydraROCKET prediction. The near-identical composition of the two stacked bars indicates that the correction modified only a small fraction of sub-burst labels and did not materially alter the overall behavioural distribution.

##### SM5.2 Derivation of analytical behavioural categories

Burst-level behavioural categories used in the ecological analyses were derived by combining the corrected TabPFN sub-burst composition with the HydraROCKET whole-burst prediction. We first identified high-agreement bursts, defined as bursts for which all four corrected TabPFN sub-bursts were assigned to the same behaviour and this behaviour matched the HydraROCKET whole-burst prediction. These bursts were directly assigned to the corresponding behaviour and constituted the majority of the dataset (~63%).

We then grouped related classifier outputs into the six final analytical categories used in the analyses. A foraging category was derived for bursts in which HydraROCKET predicted either eating or walking and TabPFN identified a mixture of feeding- and locomotion-related sub-bursts. These bursts were interpreted as active food-searching, distinct from stationary eating. Resting and sleeping were merged into a single inactive category because they were difficult to distinguish reliably from accelerometry alone and had similar relevance for the disturbance analyses. Grooming actor and grooming receiver were merged into a single social category because both represented affiliative social interactions. Self-scratching was flagged separately during classification but excluded from the final analytical categories because it showed the weakest classification performance, varied strongly among individuals, including within the same social group, and showed no coherent response centred on the culling period. Because the aim of the present study was to assess culling-related behavioural reorganisation at the group and population levels, self-scratching was therefore not retained for the main ecological analyses.

Bursts for which classifier predictions diverged were then resolved using hierarchical rules that prioritised classifier agreement and the structure of the TabPFN sub-burst composition. Bursts with a clear sub-burst majority, most commonly 3+1 compositions (~13% of bursts), were assigned to the majority behaviour when it was also supported by HydraROCKET. For example, a burst with three TabPFN sub-bursts classified as walking and one as resting was assigned to walking if HydraROCKET also predicted walking. Evenly mixed 2+2 compositions (~8% of bursts) were resolved using the HydraROCKET prediction. When the two classifiers disagreed and no clear rule applied (~4% of bursts), the final assignment was based on empirical reference distributions derived from high-agreement bursts. Bursts that remained fragmented or unresolved after this procedure were excluded from downstream analyses.

The final analytical categories were walking, running, foraging, eating, inactive behaviour, and social behaviour (Fig.SM5b). Analyses were restricted to daytime active-period bursts because the culling operation occurred during the day and including night-time periods would have strongly inflated inactive states.

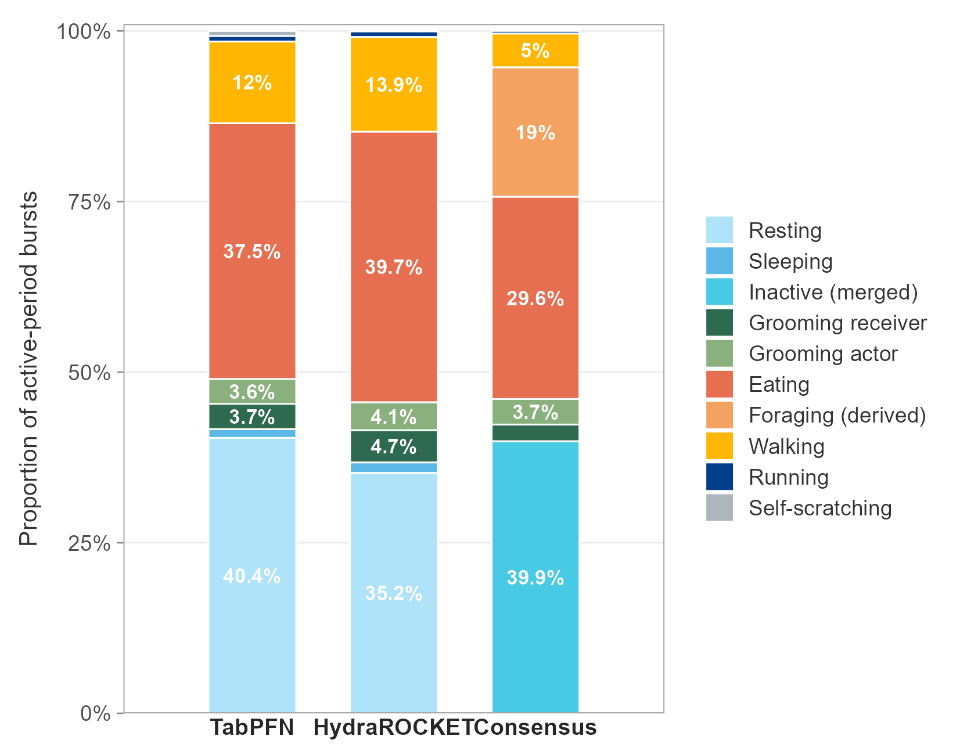

**Figure SM5b: Behavioural classification pipeline from classifiers predictions to final analytical categories.** Stacked bars show the proportional distribution of behavioural categories across active-period bursts in habituated groups after exclusion of fragmented and unresolved bursts. The first bar shows corrected TabPFN sub-burst labels, the second shows HydraROCKET whole-burst labels, and the third shows final consensus burst states used in downstream analyses. The consensus stage includes derivation of a foraging state from mixed feeding–locomotion patterns, merging of resting and sleeping into inactive behaviour, and merging of grooming actor and grooming receiver into social behaviour for the main ecological analyses.

##### SM5.3 Confidence scoring and sensitivity analyses

Each resolved burst was assigned a confidence level (i.e. very high, high, medium, or low) reflecting the strength of support for its final behavioural assignment. Confidence scoring was based on three elements: agreement between HydraROCKET and TabPFN, the structure of the corrected TabPFN sub-burst composition, and empirical thresholds derived from high-agreement bursts. These high-agreement bursts were used to derive behaviour-specific reference distributions, which were then used to arbitrate conflicting predictions and to downgrade weakly supported assignments.

Very-high-confidence bursts corresponded to the clearest cases, mainly bursts with homogeneous TabPFN sub-burst composition and a concordant HydraROCKET prediction. This category also included a small number of predefined cases in which one classifier was considered more reliable for a specific ambiguity, such as homogeneous grooming-receiver bursts for which TabPFN was prioritised over HydraROCKET. High-confidence bursts were mainly assigned to mixed compositions that could be resolved clearly by the rule set, particularly 3+1 or 2+2 patterns in which HydraROCKET supported the corresponding TabPFN majority or composition. Medium-confidence bursts were resolved cases with weaker support, including cases in which TabPFN showed a majority pattern but HydraROCKET supported the minority prediction. Low-confidence bursts were resolved cases with the weakest support. This included cases where the two classifiers disagreed and the final assignment was chosen by comparing each candidate behaviour against behaviour-specific reference distributions derived from high-agreement bursts. It also included cases initially assigned at higher confidence but downgraded because their support score fell below the 25th percentile of the corresponding high-agreement reference distribution (Fig. SM5c).

Confidence levels were used primarily as a diagnostic tool to evaluate the robustness of the classification pipeline. To assess whether the main biological patterns depended on lower-confidence assignments, we repeated the descriptive trajectory analyses using only high- and very-high-confidence bursts (Fig. SM5d). Activity budgets and temporal trajectories were highly similar to those obtained from the full set of resolved bursts, indicating that lower-confidence classifications did not materially affect the main patterns. However, this filtering disproportionately reduced the representation of some low-frequency behaviours, particularly grooming actor and other social states (Fig. SM5c). We therefore retained all resolved bursts in the final analyses and excluded only fragmented and unresolved cases.

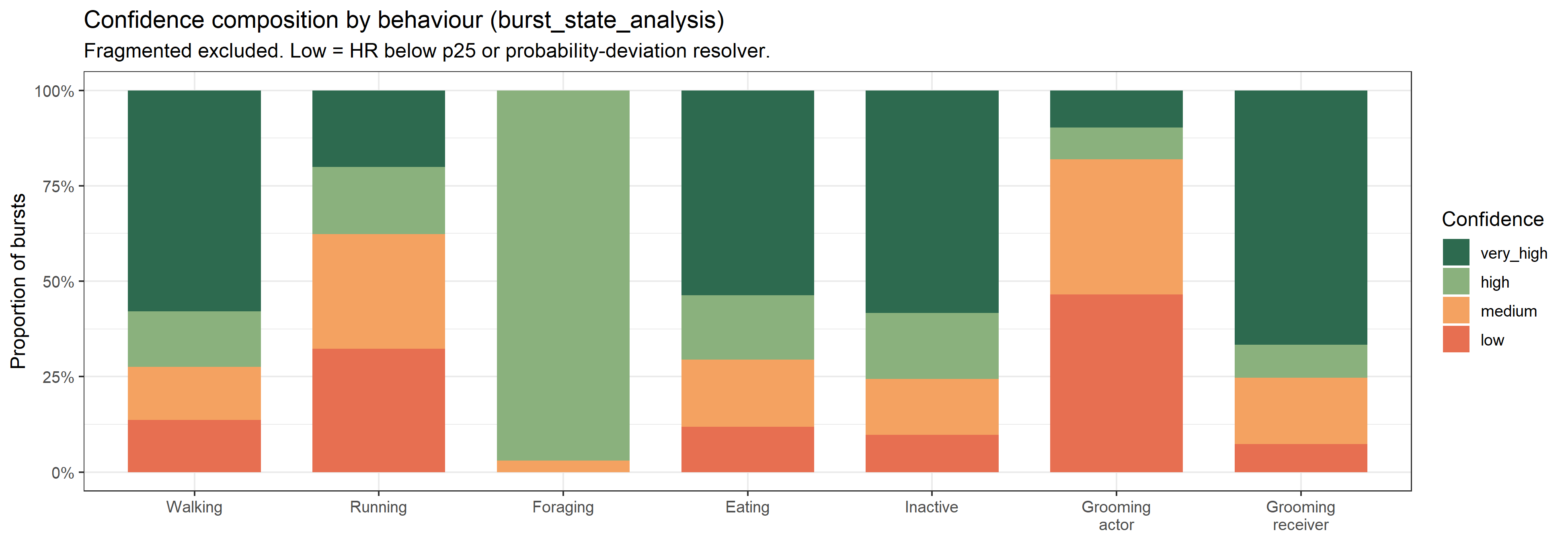

**Figure SM5c: Confidence level composition across burst states.** Stacked bars show the proportion of bursts assigned to each confidence level within each behavioural category. Confidence levels reflect the degree of support for the final behavioural assignment from the hierarchical consensus procedure. Foraging, inactive behaviour, and grooming receiver were dominated by high- or very-high-confidence assignments, whereas walking, running, and grooming actor contained larger proportions of medium- and low-confidence bursts. Self-scratching was composed almost entirely of medium-confidence assignments, reflecting its frequent detection by TabPFN without corresponding support from HydraROCKET.

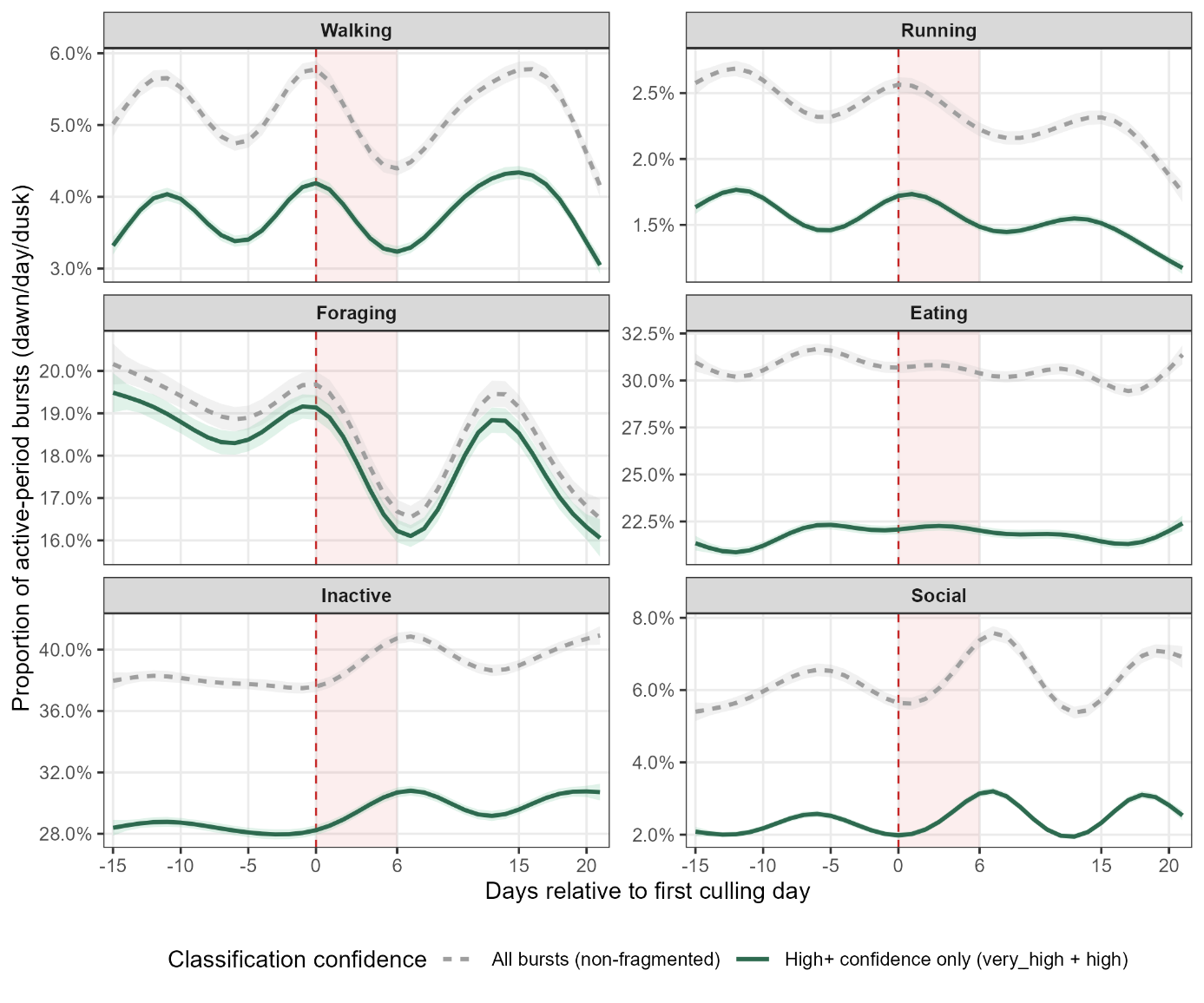

**Figure SM5d: Sensitivity of behavioural trajectories to confidence-based filtering.** Dashed grey lines show trajectories derived from all resolved bursts, whereas solid green lines show trajectories derived only from high- and very-high-confidence bursts. Trajectories were fitted over the same temporal window used for the main population-level behavioural GAMs. Shaded ribbons indicate 95% confidence intervals from weighted beta GAMs. Because confidence filtering changes the behavioural composition of the dataset, absolute proportions differ between the full and high-confidence datasets, particularly for behaviours with larger medium- or low-confidence components. The broadly similar temporal structure within behaviours indicates that the main temporal patterns were not driven by lower-confidence assignments.

### Supplementary Tables:

##### Table S1: Collared individuals included in the analyses

**Table S1.** Number of collared individuals studied across the seven groups during the culling event. All listed individuals stayed residents in one group throughout the whole focal study period. Each group included one collared female, except I-Family (IF), in which only males were collared.

| **Groups** | **Males** | **Females** | **Total** |
| --- | --- | --- | --- |
| Ankhase (AK) | 1 | 1 | **2** |
| Baie-Dankie (BD) | 6 | 1 | **7** |
| Crossing (CR) | 1 | 1 | **2** |
| I-Family (IF) | 2 |  | **2** |
| Kubu (KB) | 1 | 1 | **2** |
| Lemon-Tree (LT) | 1 | 1 | **2** |
| Noha (NH) | 3 | 1 | **4** |
| **Total** | **15** | **6** | **21** |

##### Table S2: Model selection for movement, space use, and habitat composition metrics.

Table S2: GLMMs were fitted using a Gamma distribution (log link) for continuous responses (daily path length, MCP area) and a Beta distribution (logit link) for proportional responses (dense and semi-open vegetation). Candidate models included (i) interaction models (event phase × group), (ii) additive models, (iii) population-level models, (iv) null models, and (v) models excluding temporal random effects. For daily path length, both individual identity (ID) and date were included as random effects. For MCP area and vegetation metrics, data were aggregated at the group-day level (ID = group), and only day was included as a random effect. Model selection was based on AIC (ΔAIC within each response). While a simpler structure was supported for daily path length, interaction models were retained for MCP and vegetation metrics due to the lower replication (one observation per group per day) and the partial dependence of MCP area on daily movement.

| **Metric** | **Family** | **Model** | **Formula** | **df** | **AIC** | **ΔAIC** | **Selected** |
| --- | --- | --- | --- | --- | --- | --- | --- |
| Daily path length | Gamma (log) | **no_ID** | **path_km ~ event_phase * group + (1 \| date_f)** | **23** | **366.8** | **0** | **Yes** |
|  |  | interaction | path_km ~ event_phase * group + (1 \| date_f) + (1 \| ID) | 24 | 368.8 | 2 |  |
|  |  | additive | path_km ~ event_phase + group + (1 \| date_f) + (1 \| ID) | 12 | 378.3 | 11.5 |  |
|  |  | pop_null | path_km ~ 1 + (1 \| group) + (1 \| date_f) + (1 \| ID) | 5 | 394.1 | 27.3 |  |
|  |  | population | path_km ~ event_phase + (1 \| group) + (1 \| date_f) + (1 \| ID) | 7 | 396.8 | 29.9 |  |
|  |  | no_date | path_km ~ event_phase * group + (1 \| ID) | 23 | 434.7 | 67.9 |  |
| MCP area (ha) | Gamma (log) | additive | area_ha ~ event_phase + group + (1 \| day) | 10 | 1306.4 | 0 |  |
|  |  | add_no_day | area_ha ~ event_phase + group | 9 | 1309.4 | 3.1 |  |
|  |  | **interaction** | **area_ha ~ event_phase * group + (1 \| day)** | **20** | **1313.3** | **6.9** | **Yes** |
|  |  | pop_null | area_ha ~ 1 + (1 \| group) + (1 \| day) | 4 | 1313.9 | 7.5 |  |
|  |  | no_day | area_ha ~ event_phase * group | 19 | 1315.1 | 8.8 |  |
|  |  | population | area_ha ~ event_phase + (1 \| group) + (1 \| day) | 6 | 1316.1 | 9.8 |  |
| % Dense vegetation | Beta (logit) | add_no_day | dense_beta ~ event_phase + group | 9 | -20.2 | 0 |  |
|  |  | additive | dense_beta ~ event_phase + group + (1 \| day) | 10 | -18.7 | 1.5 |  |
|  |  | pop_null | dense_beta ~ 1 + (1 \| group) + (1 \| day) | 4 | -12.7 | 7.5 |  |
|  |  | no_day | dense_beta ~ event_phase * group | 19 | -11.1 | 9.1 |  |
|  |  | population | dense_beta ~ event_phase + (1 \| group) + (1 \| day) | 6 | -10.1 | 10.1 |  |
|  |  | **interaction** | **dense_beta ~ event_phase * group + (1 \| day)** | **20** | **-9.9** | **10.3** | **Yes** |
| % Semi-open vegetation | Beta (logit) | add_no_day | semi_beta ~ event_phase + group | 9 | -23.7 | 0 |  |
|  |  | additive | semi_beta ~ event_phase + group + (1 \| day) | 10 | -22 | 1.7 |  |
|  |  | no_day | semi_beta ~ event_phase * group | 19 | -19.7 | 4 |  |
|  |  | **interaction** | **semi_beta ~ event_phase * group + (1 \| day)** | **20** | **-18.6** | **5.1** | **Yes** |
|  |  | pop_null | semi_beta ~ 1 + (1 \| group) + (1 \| day) | 4 | -15 | 8.7 |  |
|  |  | population | semi_beta ~ event_phase + (1 \| group) + (1 \| day) | 6 | -12.4 | 11.2 |  |

##### Table S3: Model selection for acceleration-based behavioural categories

**Table S3: Model selection for acceleration-based behavioural categories.** Daily behavioural proportions were analysed using GLMMs with a beta distribution (logit link). For each behaviour, three model structures were compared: population (phase), additive (phase + group), and interaction (phase × group). All models included random intercepts for individual (ID_key) and date (date_f). Model selection was based on AIC; ΔAIC (dAIC) values are relative to the best model. Models with ΔAIC < 2 were considered equivalent, and the more parsimonious model was retained. Selected models are indicated.

| **Behaviour** | **Family** | **Model** | **Formula** | **df** | **AIC** | **dAIC** | **Selected** |
| --- | --- | --- | --- | --- | --- | --- | --- |
| **Walking** | **Beta (logit)** | **interaction** | **prop_beta ~ phase * group + (1 \| ID_key) + (1 \| date_f)** | **24** | **-3965.6** | **0.0** | **Yes** |
|  |  | additive | prop_beta ~ phase + group + (1 \| ID_key) + (1 \| date_f) | 12 | -3963.6 | 2.0 |  |
|  |  | population | prop_beta ~ phase + (1 \| group) + (1 \| ID_key) + (1 \| date_f) | 7 | -3958.7 | 6.9 |  |
| **Foraging** | **Beta (logit)** | **population** | **prop_beta ~ phase + (1 \| group) + (1 \| ID_key) + (1 \| date_f)** | **7** | **-2952.7** | **0.0** | **Yes** |
|  |  | additive | prop_beta ~ phase + group + (1 \| ID_key) + (1 \| date_f) | 12 | -2951.7 | 1.0 |  |
|  |  | interaction | prop_beta ~ phase * group + (1 \| ID_key) + (1 \| date_f) | 24 | -2944.1 | 8.6 |  |
| **Eating** | **Beta (logit)** | **interaction** | **prop_beta ~ phase * group + (1 \| ID_key) + (1 \| date_f)** | **24** | **-2567.9** | **0.0** | **Yes** |
|  |  | additive | prop_beta ~ phase + group + (1 \| ID_key) + (1 \| date_f) | 12 | -2532.6 | 35.3 |  |
|  |  | population | prop_beta ~ phase + (1 \| group) + (1 \| ID_key) + (1 \| date_f) | 7 | -2531.8 | 36.1 |  |
| **Inactive** | **Beta (logit)** | **interaction** | **prop_beta ~ phase * group + (1 \| ID_key) + (1 \| date_f)** | **24** | **-2336.2** | **0.0** | **Yes** |
|  |  | population | prop_beta ~ phase + (1 \| group) + (1 \| ID_key) + (1 \| date_f) | 7 | -2320.2 | 16.1 |  |
|  |  | additive | prop_beta ~ phase + group + (1 \| ID_key) + (1 \| date_f) | 12 | -2317.6 | 18.6 |  |
| **Social** | **Beta (logit)** | **population** | **prop_beta ~ phase + (1 \| group) + (1 \| ID_key) + (1 \| date_f)** | **7** | **-3378.1** | **0.0** | **Yes** |
|  |  | additive | prop_beta ~ phase + group + (1 \| ID_key) + (1 \| date_f) | 12 | -3373.3 | 4.7 |  |
|  |  | interaction | prop_beta ~ phase * group + (1 \| ID_key) + (1 \| date_f) | 24 | -3366.3 | 11.8 |  |
| **Running** | **Beta (logit)** | **interaction** | **prop_beta ~ phase * group + (1 \| ID_key) + (1 \| date_f)** | **24** | **-4824.1** | **0.0** | **Yes** |
|  |  | additive | prop_beta ~ phase + group + (1 \| ID_key) + (1 \| date_f) | 12 | -4817.1 | 7.0 |  |
|  |  | population | prop_beta ~ phase + (1 \| group) + (1 \| ID_key) + (1 \| date_f) | 7 | -4813.7 | 10.4 |  |

##### Table S4: Space use metrics and their contrasts

**Table S4:** Estimated marginal means (EMM) and pairwise contrasts between phases for daily path length, MCP area, and proportional cover of dense and semi-open vegetation. *EMM ref* and *EMM comp* denote model-estimated means for the reference (i.e. before/ before/ during) and comparison phases (i.e. during/ after/after) respectively. *Delta* represents the absolute difference between phases (metres for daily path length, hectares for MCP area, and percentage points for vegetation). Effects are expressed as ratios (comparison/reference) for daily path length and MCP area, and as odds ratios (OR) for vegetation models. Values >1 indicate increases and <1 decreases; percentage change is shown in parentheses for ratios. Confidence intervals are given on the ratio/OR scale. *p (FDR)* indicates Benjamini–Hochberg false discovery rate–adjusted p-values within each response.

| **Response** | **Contrast** | **Group** | **EMM ref** | **EMM comp** | **Delta** | **Ratio / OR** | **95% CI** | **p (FDR)** |
| --- | --- | --- | --- | --- | --- | --- | --- | --- |
| **Daily path length** | During - Before | AK | 0.675 km | 0.531 km | -144 m | 0.787 (-21.3%) | [0.614, 1.008] | 0.0578 |
|  |  | BD | 1.081 km | 1.132 km | 51.6 m | 1.048 (+4.8%) | [0.887, 1.238] | 0.5831 |
|  |  | CR | 0.87 km | 0.702 km | -168.1 m | 0.807 (-19.3%) | [0.631, 1.031] | 0.0867 |
|  |  | IF | 0.84 km | 0.859 km | 18.5 m | 1.022 (+2.2%) | [0.788, 1.325] | 0.8693 |
|  |  | KB | 0.722 km | 0.61 km | -111.1 m | 0.846 (-15.4%) | [0.655, 1.092] | 0.1996 |
|  |  | LT | 0.691 km | 0.65 km | -40.6 m | 0.941 (-5.9%) | [0.731, 1.212] | 0.6391 |
|  |  | NH | 1.08 km | 0.874 km | -205.6 m | 0.810 (-19.0%) | [0.653, 1.004] | 0.0549 |
|  | After - Before | AK | 0.675 km | 0.608 km | -67.1 m | 0.901 (-9.9%) | [0.737, 1.101] | 0.3082 |
|  |  | BD | 1.081 km | 1.047 km | -33.2 m | 0.969 (-3.1%) | [0.849, 1.106] | 0.6439 |
|  |  | CR | 0.87 km | 0.768 km | -101.7 m | 0.883 (-11.7%) | [0.725, 1.076] | 0.2175 |
|  |  | IF | 0.84 km | 1.022 km | 181.6 m | 1.216 (+21.6%) | [0.994, 1.488] | 0.0571 |
|  |  | KB | 0.722 km | 0.78 km | 58.1 m | 1.081 (+8.1%) | [0.888, 1.314] | 0.4378 |
|  |  | LT | 0.691 km | 0.91 km | 218.9 m | 1.317 (+31.7%) | [1.077, 1.610] | 0.0073 |
|  |  | NH | 1.08 km | 1.011 km | -69.2 m | 0.936 (-6.4%) | [0.791, 1.108] | 0.4420 |
|  | After - During | AK | 0.531 km | 0.608 km | 76.9 m | 1.145 (+14.5%) | [0.893, 1.467] | 0.2854 |
|  |  | BD | 1.132 km | 1.047 km | -84.8 m | 0.925 (-7.5%) | [0.783, 1.093] | 0.3606 |
|  |  | CR | 0.702 km | 0.768 km | 66.3 m | 1.095 (+9.5%) | [0.856, 1.400] | 0.4722 |
|  |  | IF | 0.859 km | 1.022 km | 163.1 m | 1.190 (+19.0%) | [0.922, 1.536] | 0.1824 |
|  |  | KB | 0.61 km | 0.78 km | 169.2 m | 1.277 (+27.7%) | [0.989, 1.649] | 0.0604 |
|  |  | LT | 0.65 km | 0.91 km | 259.5 m | 1.399 (+39.9%) | [1.088, 1.798] | 0.0087 |
|  |  | NH | 0.874 km | 1.011 km | 136.5 m | 1.156 (+15.6%) | [0.928, 1.440] | 0.1959 |
| **MCP area (ha)** | During - Before | AK | 5.46 ha | 4.31 ha | -1.15 ha | 0.789 (-21.1%) | [0.391, 1.594] | 0.5089 |
|  |  | BD | 13.91 ha | 14.43 ha | 0.52 ha | 1.037 (+3.7%) | [0.509, 2.113] | 0.9202 |
|  |  | CR | 7.41 ha | 8.3 ha | 0.88 ha | 1.119 (+11.9%) | [0.556, 2.254] | 0.7523 |
|  |  | KB | 5.29 ha | 2.98 ha | -2.31 ha | 0.563 (-43.7%) | [0.279, 1.133] | 0.1075 |
|  |  | LT | 6.88 ha | 4.96 ha | -1.92 ha | 0.721 (-27.9%) | [0.325, 1.599] | 0.4207 |
|  |  | NH | 15.58 ha | 6.9 ha | -8.68 ha | 0.443 (-55.7%) | [0.213, 0.922] | 0.0294 |
|  | After - Before | AK | 5.46 ha | 4.99 ha | -0.47 ha | 0.914 (-8.6%) | [0.516, 1.618] | 0.7576 |
|  |  | BD | 13.91 ha | 9.06 ha | -4.86 ha | 0.651 (-34.9%) | [0.373, 1.137] | 0.1312 |
|  |  | CR | 7.41 ha | 7.58 ha | 0.17 ha | 1.023 (+2.3%) | [0.576, 1.814] | 0.9392 |
|  |  | KB | 5.29 ha | 5.97 ha | 0.68 ha | 1.128 (+12.8%) | [0.644, 1.976] | 0.6724 |
|  |  | LT | 6.88 ha | 9.97 ha | 3.09 ha | 1.450 (+45.0%) | [0.809, 2.597] | 0.2118 |
|  |  | NH | 15.58 ha | 10.62 ha | -4.96 ha | 0.682 (-31.8%) | [0.387, 1.201] | 0.1845 |
|  | After - During | AK | 4.31 ha | 4.99 ha | 0.68 ha | 1.158 (+15.8%) | [0.569, 2.357] | 0.6851 |
|  |  | BD | 14.43 ha | 9.06 ha | -5.37 ha | 0.628 (-37.2%) | [0.309, 1.276] | 0.1981 |
|  |  | CR | 8.3 ha | 7.58 ha | -0.72 ha | 0.914 (-8.6%) | [0.451, 1.850] | 0.8017 |
|  |  | KB | 2.98 ha | 5.97 ha | 2.99 ha | 2.006 (+100.6%) | [0.997, 4.036] | 0.0511 |
|  |  | LT | 4.96 ha | 9.97 ha | 5.01 ha | 2.011 (+101.1%) | [0.911, 4.439] | 0.0836 |
|  |  | NH | 6.9 ha | 10.62 ha | 3.72 ha | 1.539 (+53.9%) | [0.735, 3.223] | 0.2529 |
| **% Dense veg.** | During - Before | AK | 33.5% | 33.4% | -0.15 pp | 0.993 | [0.368, 2.683] | 0.9891 |
|  |  | BD | 47.8% | 55.1% | 7.34 pp | 1.343 | [0.502, 3.590] | 0.5573 |
|  |  | CR | 50.8% | 61.8% | 10.95 pp | 1.564 | [0.581, 4.214] | 0.3765 |
|  |  | KB | 50.5% | 75.8% | 25.26 pp | 3.064 | [1.126, 8.335] | 0.0283 |
|  |  | LT | 69.4% | 62.5% | -6.83 pp | 0.737 | [0.241, 2.255] | 0.5930 |
|  |  | NH | 46.8% | 32.6% | -14.16 pp | 0.551 | [0.191, 1.586] | 0.2690 |
|  | After - Before | AK | 33.5% | 25.5% | -8.04 pp | 0.678 | [0.304, 1.515] | 0.3436 |
|  |  | BD | 47.8% | 59.4% | 11.6 pp | 1.597 | [0.728, 3.506] | 0.2429 |
|  |  | CR | 50.8% | 46.7% | -4.17 pp | 0.846 | [0.379, 1.888] | 0.6832 |
|  |  | KB | 50.5% | 60.3% | 9.85 pp | 1.492 | [0.678, 3.282] | 0.3202 |
|  |  | LT | 69.4% | 53.8% | -15.58 pp | 0.514 | [0.229, 1.155] | 0.1072 |
|  |  | NH | 46.8% | 40.7% | -6.08 pp | 0.781 | [0.349, 1.747] | 0.5469 |
|  | After - During | AK | 33.4% | 25.5% | -7.88 pp | 0.683 | [0.249, 1.870] | 0.4582 |
|  |  | BD | 55.1% | 59.4% | 4.25 pp | 1.190 | [0.447, 3.170] | 0.7281 |
|  |  | CR | 61.8% | 46.7% | -15.13 pp | 0.541 | [0.199, 1.473] | 0.2294 |
|  |  | KB | 75.8% | 60.3% | -15.42 pp | 0.487 | [0.180, 1.317] | 0.1563 |
|  |  | LT | 62.5% | 53.8% | -8.75 pp | 0.697 | [0.228, 2.128] | 0.5266 |
|  |  | NH | 32.6% | 40.7% | 8.07 pp | 1.417 | [0.486, 4.128] | 0.5228 |
| **% Semi-open veg.** | During - Before | AK | 57.8% | 67.5% | 9.72 pp | 1.518 | [0.591, 3.897] | 0.3857 |
|  |  | BD | 49.8% | 38.3% | -11.46 pp | 0.627 | [0.246, 1.596] | 0.3272 |
|  |  | CR | 43.6% | 36.1% | -7.49 pp | 0.731 | [0.286, 1.871] | 0.5137 |
|  |  | KB | 46.8% | 22.7% | -24.13 pp | 0.333 | [0.128, 0.868] | 0.0244 |
|  |  | LT | 22.6% | 30.3% | 7.65 pp | 1.484 | [0.506, 4.359] | 0.4723 |
|  |  | NH | 53.2% | 68.5% | 15.33 pp | 1.916 | [0.703, 5.221] | 0.2039 |
|  | After - Before | AK | 57.8% | 69.8% | 11.99 pp | 1.687 | [0.788, 3.609] | 0.1778 |
|  |  | BD | 49.8% | 36.9% | -12.88 pp | 0.590 | [0.280, 1.242] | 0.1645 |
|  |  | CR | 43.6% | 47.2% | 3.55 pp | 1.154 | [0.539, 2.472] | 0.7123 |
|  |  | KB | 46.8% | 36.6% | -10.17 pp | 0.657 | [0.312, 1.386] | 0.2702 |
|  |  | LT | 22.6% | 44.4% | 21.73 pp | 2.725 | [1.252, 5.932] | 0.0115 |
|  |  | NH | 53.2% | 59.9% | 6.78 pp | 1.319 | [0.616, 2.824] | 0.4766 |
|  | After - During | AK | 67.5% | 69.8% | 2.27 pp | 1.111 | [0.426, 2.898] | 0.8292 |
|  |  | BD | 38.3% | 36.9% | -1.42 pp | 0.941 | [0.370, 2.393] | 0.8988 |
|  |  | CR | 36.1% | 47.2% | 11.04 pp | 1.579 | [0.610, 4.086] | 0.3468 |
|  |  | KB | 22.7% | 36.6% | 13.97 pp | 1.972 | [0.759, 5.124] | 0.1633 |
|  |  | LT | 30.3% | 44.4% | 14.08 pp | 1.836 | [0.632, 5.336] | 0.2645 |
|  |  | NH | 68.5% | 59.9% | -8.55 pp | 0.688 | [0.249, 1.899] | 0.4709 |

##### Table S5: Acceleration-based behavioural categories and estimated marginal means

**Table S5:** Estimated marginal means (EMM) and pairwise contrasts between phases for behavioural proportions across groups. Behaviours include **walking, foraging, eating, inactive, social, and running**. *EMM ref* and *EMM comp* denote model-estimated means (%) for the reference and comparison phases defined by each contrast (e.g. before for “during–before” and “after–before”; during for “after–during”). *Δ pp* represents the absolute difference between phases in percentage points. Effects are expressed as odds ratios (OR) derived from models fitted on the logit scale. Values >1 indicate increases and <1 decreases relative to the reference phase. Confidence intervals are given on the OR scale. *p (FDR)* indicates Benjamini–Hochberg false discovery rate–adjusted p-values within each behaviour.

| **Behaviour** | **Best model** | **Level** | **Contrast** | **Group** | **EMM ref (%)** | **EMM comp (%)** | **Δ pp** | **Odds Ratio** | **95% CI** | **p (FDR)** |  |
| --- | --- | --- | --- | --- | --- | --- | --- | --- | --- | --- | --- |
| Walking | M_int | group | During - Before | AK | 3.10 | 3.24 | 0.14 | 1.047 | [0.797, 1.375] | 0.9700 | NS |
|  |  |  | During - Before | BD | 6.30 | 5.35 | -0.95 | 0.840 | [0.717, 0.984] | 0.0585 | NS |
|  |  |  | During - Before | CR | 4.14 | 3.30 | -0.84 | 0.790 | [0.554, 1.126] | 0.4248 | NS |
|  |  |  | During - Before | IF | 5.21 | 5.34 | 0.13 | 1.027 | [0.817, 1.291] | 0.8198 | NS |
|  |  |  | During - Before | KB | 3.91 | 3.28 | -0.63 | 0.833 | [0.638, 1.087] | 0.2678 | NS |
|  |  |  | During - Before | LT | 3.98 | 3.85 | -0.14 | 0.964 | [0.749, 1.243] | 0.7793 | NS |
|  |  |  | During - Before | NH | 6.04 | 6.05 | 0.01 | 1.003 | [0.839, 1.198] | 0.9778 | NS |
|  |  |  | After - Before | AK | 3.10 | 3.22 | 0.12 | 1.042 | [0.837, 1.296] | 0.9700 | NS |
|  |  |  | After - Before | BD | 6.30 | 6.25 | -0.05 | 0.992 | [0.877, 1.123] | 0.9027 | NS |
|  |  |  | After - Before | CR | 4.14 | 3.98 | -0.16 | 0.960 | [0.734, 1.256] | 0.7670 | NS |
|  |  |  | After - Before | IF | 5.21 | 4.77 | -0.44 | 0.912 | [0.756, 1.099] | 0.4968 | NS |
|  |  |  | After - Before | KB | 3.91 | 3.96 | 0.05 | 1.015 | [0.828, 1.244] | 0.8889 | NS |
|  |  |  | After - Before | LT | 3.98 | 4.85 | 0.86 | 1.228 | [1.011, 1.492] | 0.0846 | NS |
|  |  |  | After - Before | NH | 6.04 | 5.35 | -0.68 | 0.880 | [0.761, 1.018] | 0.2345 | NS |
|  |  |  | After - During | AK | 3.24 | 3.22 | -0.02 | 0.995 | [0.758, 1.305] | 0.9700 | NS |
|  |  |  | After - During | BD | 5.35 | 6.25 | 0.91 | 1.181 | [1.008, 1.383] | 0.0585 | NS |
|  |  |  | After - During | CR | 3.30 | 3.98 | 0.68 | 1.215 | [0.851, 1.734] | 0.4248 | NS |
|  |  |  | After - During | IF | 5.34 | 4.77 | -0.57 | 0.888 | [0.704, 1.119] | 0.4968 | NS |
|  |  |  | After - During | KB | 3.28 | 3.96 | 0.69 | 1.218 | [0.934, 1.589] | 0.2678 | NS |
|  |  |  | After - During | LT | 3.85 | 4.85 | 1.00 | 1.274 | [0.993, 1.633] | 0.0846 | NS |
|  |  |  | After - During | NH | 6.05 | 5.35 | -0.70 | 0.878 | [0.733, 1.051] | 0.2345 | NS |
| Foraging | M_pop | population | During - Before | Population | 19.76 | 17.70 | -2.06 | 0.873 | [0.785, 0.97] | 0.0178 |  |
|  |  |  | After - Before | Population | 19.76 | 18.03 | -1.73 | 0.893 | [0.821, 0.972] | 0.0178 |  |
|  |  |  | After - During | Population | 17.70 | 18.03 | 0.34 | 1.023 | [0.92, 1.138] | 0.6698 | NS |
| Eating | M_int | group | During - Before | AK | 32.91 | 33.12 | 0.21 | 1.010 | [0.885, 1.151] | 0.8878 | NS |
|  |  |  | During - Before | BD | 32.43 | 31.57 | -0.86 | 0.961 | [0.878, 1.052] | 0.4308 | NS |
|  |  |  | During - Before | CR | 26.34 | 26.29 | -0.05 | 0.997 | [0.83, 1.198] | 0.9761 | NS |
|  |  |  | During - Before | IF | 28.44 | 32.06 | 3.62 | 1.188 | [1.039, 1.357] | 0.0171 |  |
|  |  |  | During - Before | KB | 31.81 | 30.76 | -1.04 | 0.953 | [0.834, 1.088] | 0.4740 | NS |
|  |  |  | During - Before | LT | 31.61 | 34.04 | 2.43 | 1.117 | [0.979, 1.273] | 0.1992 | NS |
|  |  |  | During - Before | NH | 31.41 | 29.95 | -1.46 | 0.933 | [0.84, 1.037] | 0.5072 | NS |
|  |  |  | After - Before | AK | 32.91 | 29.87 | -3.04 | 0.868 | [0.781, 0.965] | 0.0267 |  |
|  |  |  | After - Before | BD | 32.43 | 30.79 | -1.64 | 0.927 | [0.863, 0.996] | 0.1152 | NS |
|  |  |  | After - Before | CR | 26.34 | 23.44 | -2.90 | 0.856 | [0.738, 0.994] | 0.1238 | NS |
|  |  |  | After - Before | IF | 28.44 | 33.26 | 4.82 | 1.254 | [1.127, 1.395] | 0.0001 | *** |
|  |  |  | After - Before | KB | 31.81 | 27.73 | -4.08 | 0.823 | [0.739, 0.916] | 0.0011 | ** |
|  |  |  | After - Before | LT | 31.61 | 31.82 | 0.21 | 1.010 | [0.909, 1.122] | 0.8561 | NS |
|  |  |  | After - Before | NH | 31.41 | 30.70 | -0.71 | 0.967 | [0.89, 1.052] | 0.5072 | NS |
|  |  |  | After - During | AK | 33.12 | 29.87 | -3.25 | 0.860 | [0.753, 0.982] | 0.0381 |  |
|  |  |  | After - During | BD | 31.57 | 30.79 | -0.78 | 0.964 | [0.881, 1.055] | 0.4308 | NS |
|  |  |  | After - During | CR | 26.29 | 23.44 | -2.84 | 0.859 | [0.713, 1.034] | 0.1616 | NS |
|  |  |  | After - During | IF | 32.06 | 33.26 | 1.20 | 1.056 | [0.926, 1.205] | 0.4177 | NS |
|  |  |  | After - During | KB | 30.76 | 27.73 | -3.03 | 0.864 | [0.755, 0.988] | 0.0484 |  |
|  |  |  | After - During | LT | 34.04 | 31.82 | -2.22 | 0.904 | [0.793, 1.031] | 0.1992 | NS |
|  |  |  | After - During | NH | 29.95 | 30.70 | 0.75 | 1.036 | [0.933, 1.152] | 0.5072 | NS |
| Inactive |  |  | During - Before | AK | 38.71 | 40.55 | 1.84 | 1.080 | [0.94, 1.241] | 0.2776 | NS |
|  |  |  | During - Before | BD | 34.55 | 36.66 | 2.10 | 1.096 | [0.996, 1.207] | 0.0916 | NS |
|  |  |  | During - Before | CR | 42.72 | 47.76 | 5.04 | 1.226 | [1.023, 1.469] | 0.0416 |  |
|  |  |  | During - Before | IF | 39.34 | 35.82 | -3.52 | 0.861 | [0.747, 0.991] | 0.0952 | NS |
|  |  |  | During - Before | KB | 37.32 | 41.46 | 4.14 | 1.189 | [1.035, 1.368] | 0.0222 |  |
|  |  |  | During - Before | LT | 35.78 | 36.46 | 0.68 | 1.030 | [0.894, 1.186] | 0.8286 | NS |
|  |  |  | During - Before | NH | 36.68 | 38.16 | 1.48 | 1.065 | [0.953, 1.19] | 0.3980 | NS |
|  |  |  | After - Before | AK | 38.71 | 43.19 | 4.48 | 1.204 | [1.077, 1.345] | 0.0031 | ** |
|  |  |  | After - Before | BD | 34.55 | 37.82 | 3.27 | 1.152 | [1.067, 1.244] | 0.0009 | *** |
|  |  |  | After - Before | CR | 42.72 | 48.90 | 6.18 | 1.283 | [1.11, 1.483] | 0.0022 | ** |
|  |  |  | After - Before | IF | 39.34 | 36.83 | -2.51 | 0.899 | [0.803, 1.006] | 0.0952 | NS |
|  |  |  | After - Before | KB | 37.32 | 42.84 | 5.52 | 1.259 | [1.126, 1.407] | 0.0002 | *** |
|  |  |  | After - Before | LT | 35.78 | 36.06 | 0.29 | 1.013 | [0.905, 1.133] | 0.8286 | NS |
|  |  |  | After - Before | NH | 36.68 | 38.19 | 1.51 | 1.067 | [0.976, 1.165] | 0.3980 | NS |
|  |  |  | After - During | AK | 40.55 | 43.19 | 2.63 | 1.114 | [0.97, 1.28] | 0.1888 | NS |
|  |  |  | After - During | BD | 36.66 | 37.82 | 1.16 | 1.051 | [0.955, 1.157] | 0.3095 | NS |
|  |  |  | After - During | CR | 47.76 | 48.90 | 1.14 | 1.047 | [0.874, 1.254] | 0.6194 | NS |
|  |  |  | After - During | IF | 35.82 | 36.83 | 1.01 | 1.045 | [0.907, 1.204] | 0.5463 | NS |
|  |  |  | After - During | KB | 41.46 | 42.84 | 1.38 | 1.058 | [0.921, 1.216] | 0.4247 | NS |
|  |  |  | After - During | LT | 36.46 | 36.06 | -0.39 | 0.983 | [0.854, 1.132] | 0.8286 | NS |
|  |  |  | After - During | NH | 38.16 | 38.19 | 0.03 | 1.001 | [0.896, 1.119] | 0.9800 | NS |
| Social | M_pop | population | During - Before | Population | 5.41 | 6.21 | 0.80 | 1.158 | [0.982, 1.365] | 0.2464 | NS |
|  |  |  | After - Before | Population | 5.41 | 5.89 | 0.47 | 1.093 | [0.957, 1.248] | 0.2833 | NS |
|  |  |  | After - During | Population | 6.21 | 5.89 | -0.33 | 0.944 | [0.801, 1.113] | 0.4927 | NS |
| Running | M_int | group | During - Before | AK | 1.86 | 1.89 | 0.03 | 1.018 | [0.759, 1.366] | 0.9043 | NS |
|  |  |  | During - Before | BD | 2.76 | 2.43 | -0.33 | 0.877 | [0.751, 1.025] | 0.1490 | NS |
|  |  |  | During - Before | CR | 2.42 | 1.97 | -0.45 | 0.810 | [0.547, 1.199] | 0.4371 | NS |
|  |  |  | During - Before | IF | 3.78 | 3.39 | -0.38 | 0.895 | [0.712, 1.124] | 0.3406 | NS |
|  |  |  | During - Before | KB | 1.41 | 0.75 | -0.66 | 0.527 | [0.355, 0.781] | 0.0043 | ** |
|  |  |  | During - Before | LT | 1.59 | 1.18 | -0.41 | 0.740 | [0.527, 1.038] | 0.1232 | NS |
|  |  |  | During - Before | NH | 2.34 | 2.37 | 0.03 | 1.014 | [0.835, 1.231] | 0.8870 | NS |
|  |  |  | After - Before | AK | 1.86 | 1.31 | -0.55 | 0.701 | [0.544, 0.903] | 0.0181 |  |
|  |  |  | After - Before | BD | 2.76 | 2.48 | -0.28 | 0.895 | [0.791, 1.012] | 0.1490 | NS |
|  |  |  | After - Before | CR | 2.42 | 2.42 | 0.00 | 1.001 | [0.745, 1.343] | 0.9967 | NS |
|  |  |  | After - Before | IF | 3.78 | 2.41 | -1.36 | 0.630 | [0.519, 0.766] | 0.0000 | *** |
|  |  |  | After - Before | KB | 1.41 | 0.98 | -0.43 | 0.692 | [0.52, 0.92] | 0.0169 |  |
|  |  |  | After - Before | LT | 1.59 | 1.27 | -0.32 | 0.794 | [0.612, 1.03] | 0.1232 | NS |
|  |  |  | After - Before | NH | 2.34 | 2.10 | -0.24 | 0.896 | [0.765, 1.049] | 0.3257 | NS |
|  |  |  | After - During | AK | 1.89 | 1.31 | -0.58 | 0.689 | [0.505, 0.939] | 0.0275 |  |
|  |  |  | After - During | BD | 2.43 | 2.48 | 0.05 | 1.020 | [0.871, 1.193] | 0.8067 | NS |
|  |  |  | After - During | CR | 1.97 | 2.42 | 0.45 | 1.236 | [0.835, 1.83] | 0.4371 | NS |
|  |  |  | After - During | IF | 3.39 | 2.41 | -0.98 | 0.704 | [0.553, 0.897] | 0.0067 | ** |
|  |  |  | After - During | KB | 0.75 | 0.98 | 0.23 | 1.313 | [0.873, 1.974] | 0.1916 | NS |
|  |  |  | After - During | LT | 1.18 | 1.27 | 0.09 | 1.073 | [0.758, 1.52] | 0.6919 | NS |
|  |  |  | After - During | NH | 2.37 | 2.10 | -0.27 | 0.884 | [0.726, 1.075] | 0.3257 | NS |

### Supplementary Figures:

##### Figure S1: **DHARMa residual diagnostics for GPS-based models**

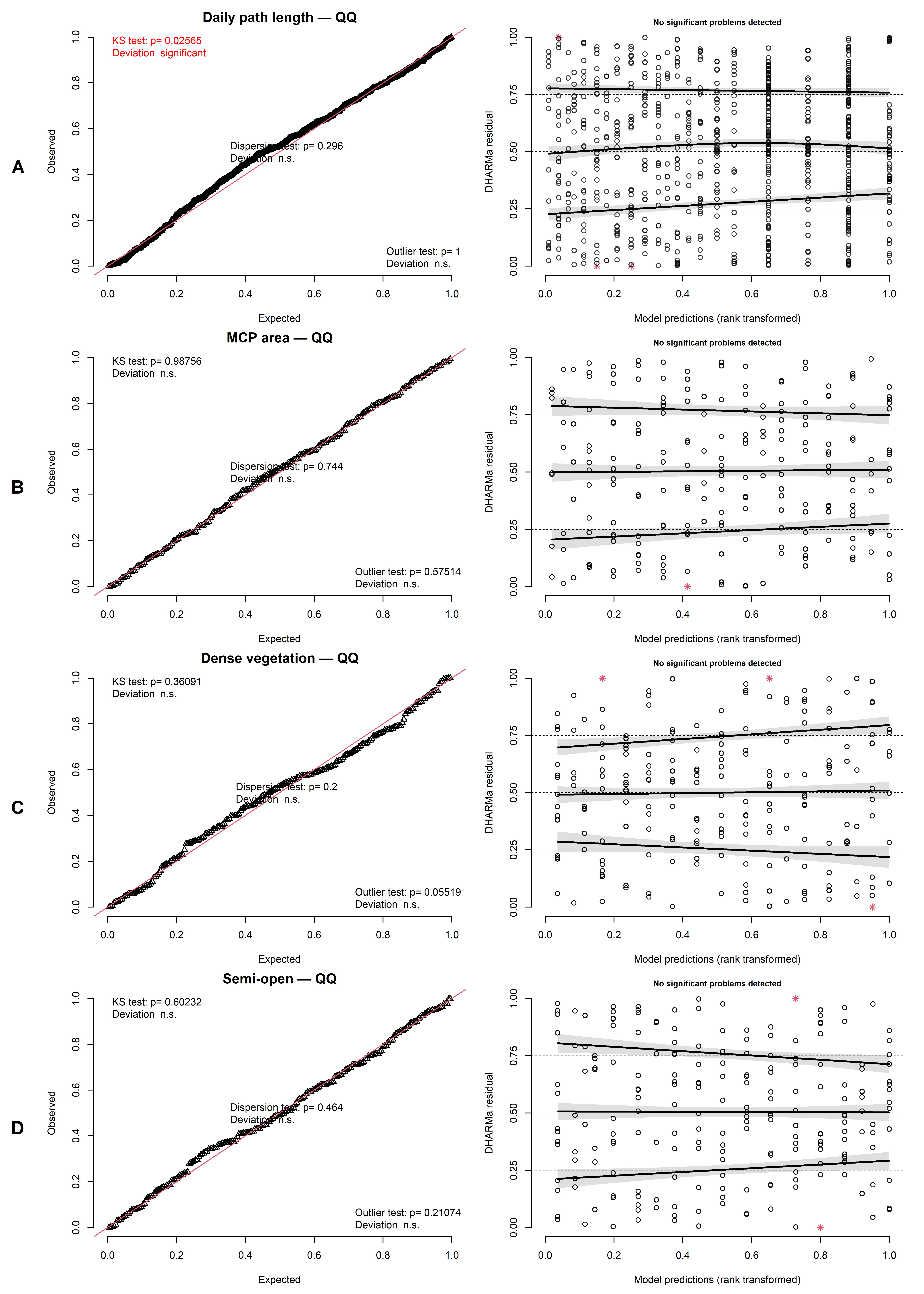

**Figure S1:** DHARMa residual diagnostics for GLMMs fitted to GPS-derived metrics. Panels A–D correspond to the different response variables (e.g. daily path length, MCP area, and habitat composition metrics). For each panel, the left plot shows the quantile–quantile (QQ) comparison of simulated residuals against the expected uniform distribution, while the right plot shows residuals versus fitted values. Model assumptions were assessed using uniformity (Kolmogorov–Smirnov test), dispersion, and outlier tests. Across models, residuals were generally well distributed with no strong systematic deviations, indicating an overall adequate model fit.

##### Figure S2: DHARMa residual diagnostics for accelerometry-based behavioural models

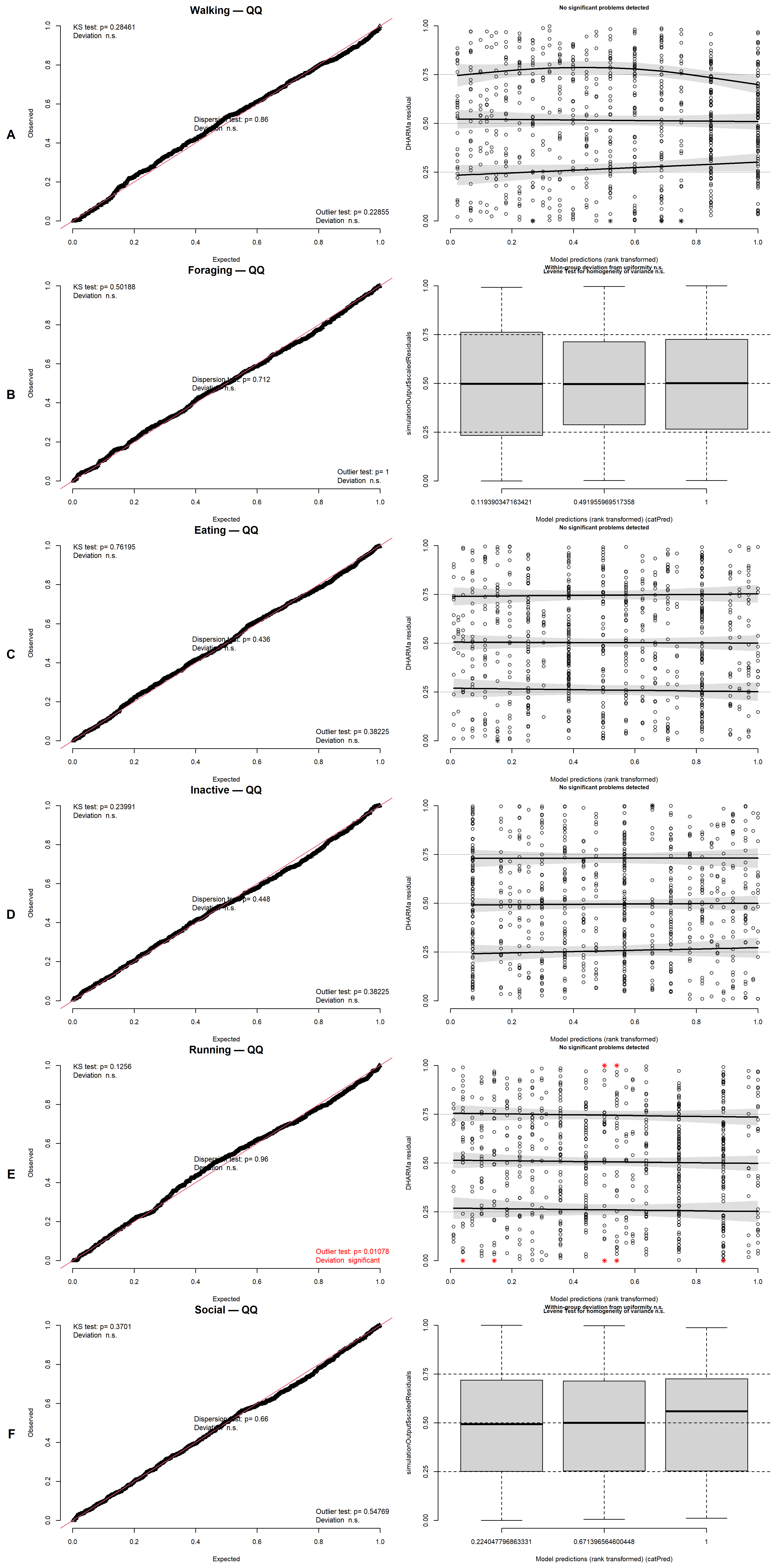

Figure S2: DHARMa residual diagnostics for GLMMs fitted to daily behavioural proportions derived from accelerometry. Panels **A–F** correspond to behavioural categories. For each behaviour, the **left plot** shows the QQ plot of simulated residuals (uniformity check), and the **right plot** shows residuals versus fitted values to assess potential structure or heteroscedasticity. Statistical tests include **uniformity (Kolmogorov–Smirnov test)**, **dispersion**, and **outlier detection**, with additional checks for **variance homogeneity** where indicated. Overall, residuals showed no major deviations from model assumptions, supporting the validity of the fitted models. Some structure was nonetheless observed in a few cases (notably *Foraging*, *Running*, and *Social*), with a **significant Levene test for Foraging**, indicating mild heteroscedasticity that did not substantially affect model interpretation.

##### Figure S3: Daily temporal trajectories of GPS-derived metrics around the May 2023 culling event.

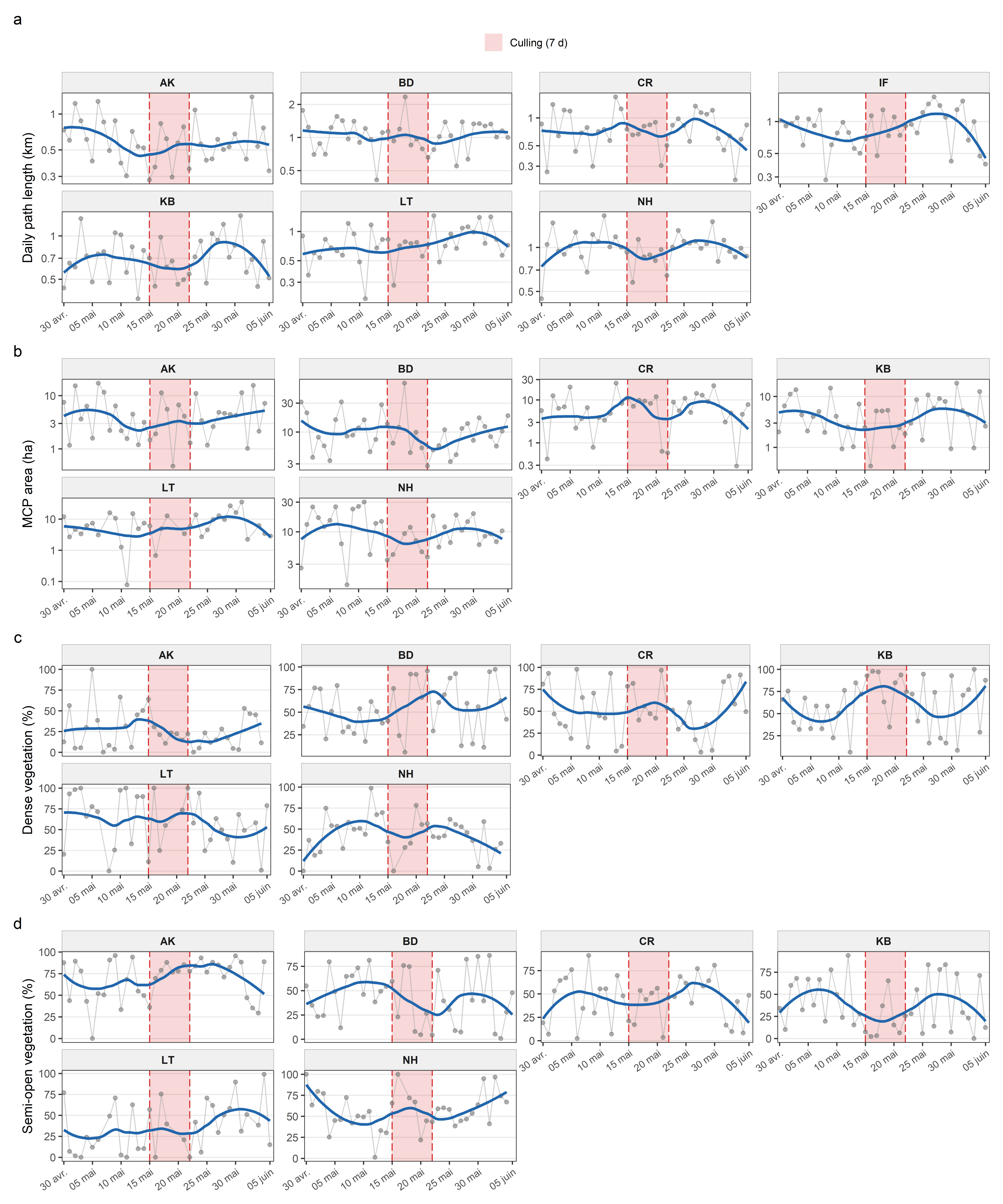

**Figure S3: Daily temporal trajectories of GPS-derived metrics around the May 2023 culling event.** Grey points and lines show observed daily values averaged at the group level; blue lines show LOESS-smoothed trends. Panels show (a) daily path length (km), (b) daily MCP area (ha), (c) dense vegetation within daily MCPs (%), and (d) semi-open vegetation within daily MCPs (%), separately for each group. Red shading indicates the culling period (days 0–7); dashed red lines mark its start and end. Daily path length is available for all groups, whereas MCP and habitat metrics are restricted to groups with valid daily female MCP estimates. These descriptive trajectories illustrate temporal heterogeneity in movement, space use, and habitat use around the culling event.

##### Figure S4: Daily temporal trajectories of accelerometry-derived behavioural proportions.

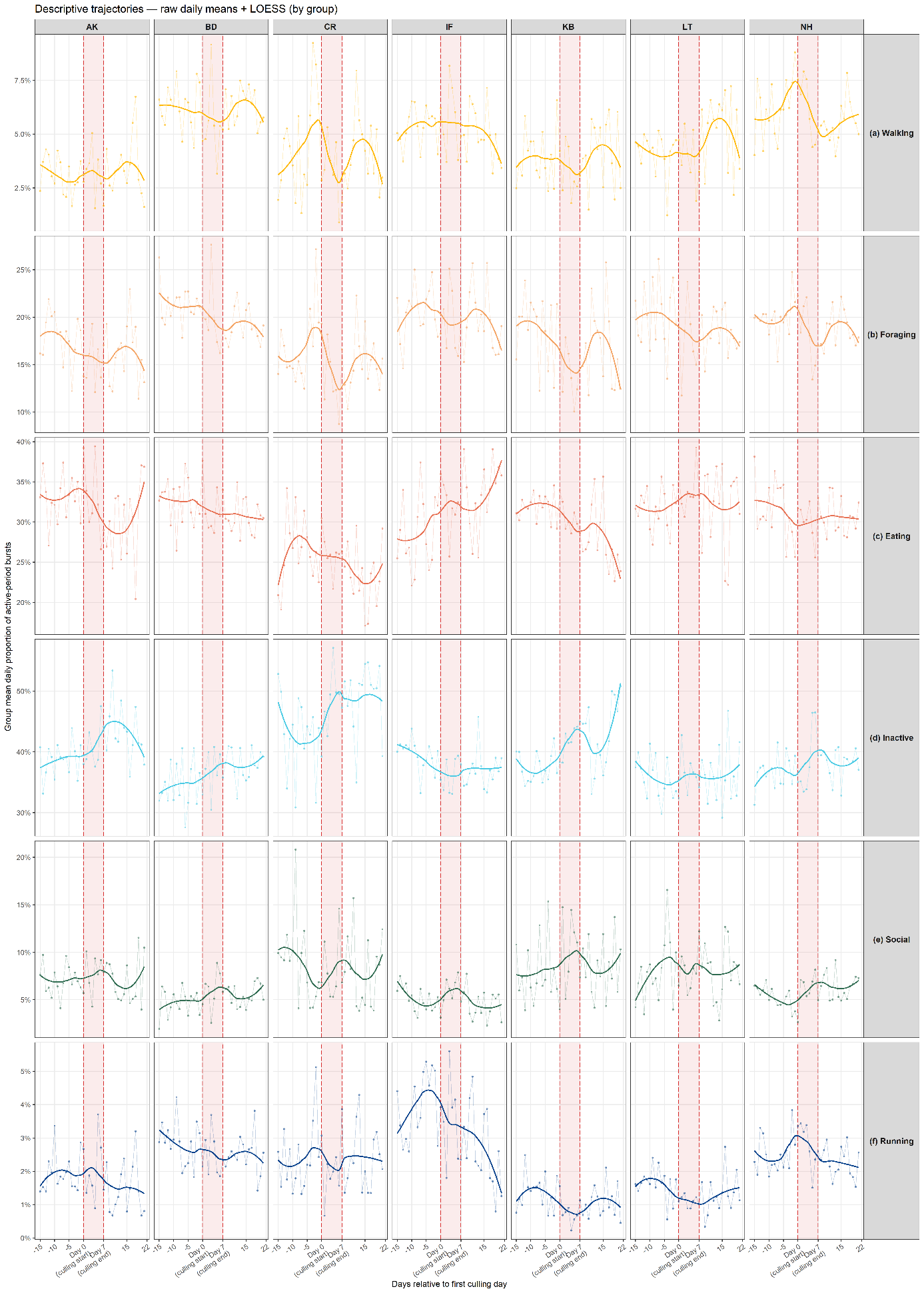

**Figure S4. Daily temporal trajectories of accelerometry-based behavioural proportions around the culling event.** Observed daily group-level mean proportions (points and thin lines) and LOESS-smoothed trends (solid coloured lines) are shown separately for each group (columns) and behavioural category (rows: (a) walking, (b) foraging, (c) eating, (d) inactive, (e) social, and (f) running). Dashed red vertical lines mark the start and end of the culling period (days 0–7) and red shading indicates the culling period. The x-axis shows days relative to the first culling day. These trajectories provide a descriptive complement to the model-based analyses by illustrating short-term temporal dynamics and heterogeneity in behavioural responses across groups.

##### Figure S5: Group-level Circadian Organisation

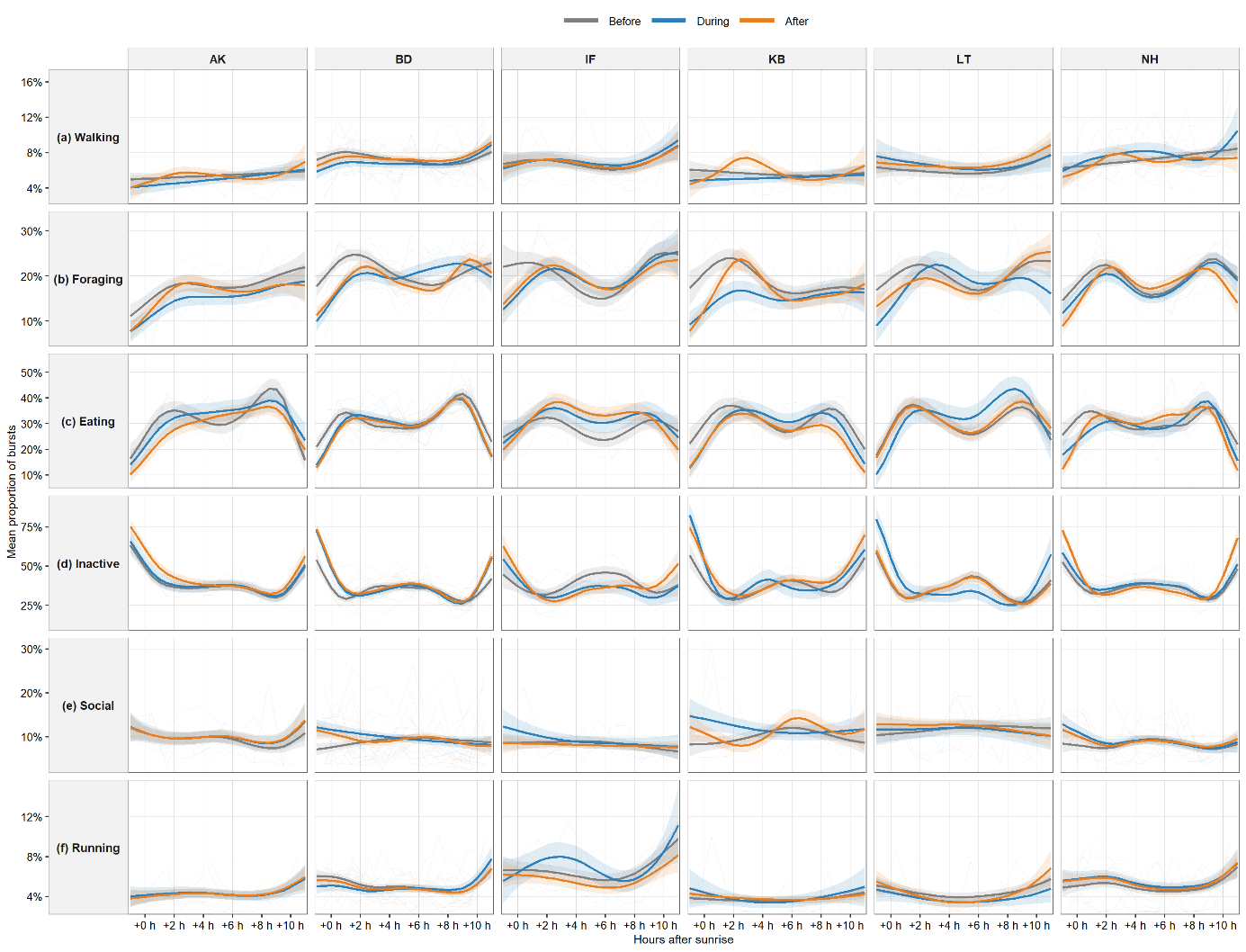

**Figure S5. Group-level circadian organisation of behaviour across disturbance phases.** Behavioural proportions were aligned relative to sunrise and plotted separately for each group and phase to compare within-day activity patterns before, during, and after the culling event. Solid lines show phase-specific smooth trajectories estimated with beta generalised additive models (GAMs), and shaded ribbons indicate 95% confidence intervals. Faint background lines show individual trajectories. These group-level curves highlight heterogeneity in the timing and magnitude of circadian responses among groups, while showing that several groups exhibited reduced foraging and eating together with increased inactivity during the early active period of the culling phase.

##### Figure S6: Standardised residual summaries of acute locomotor deviations during the culling period

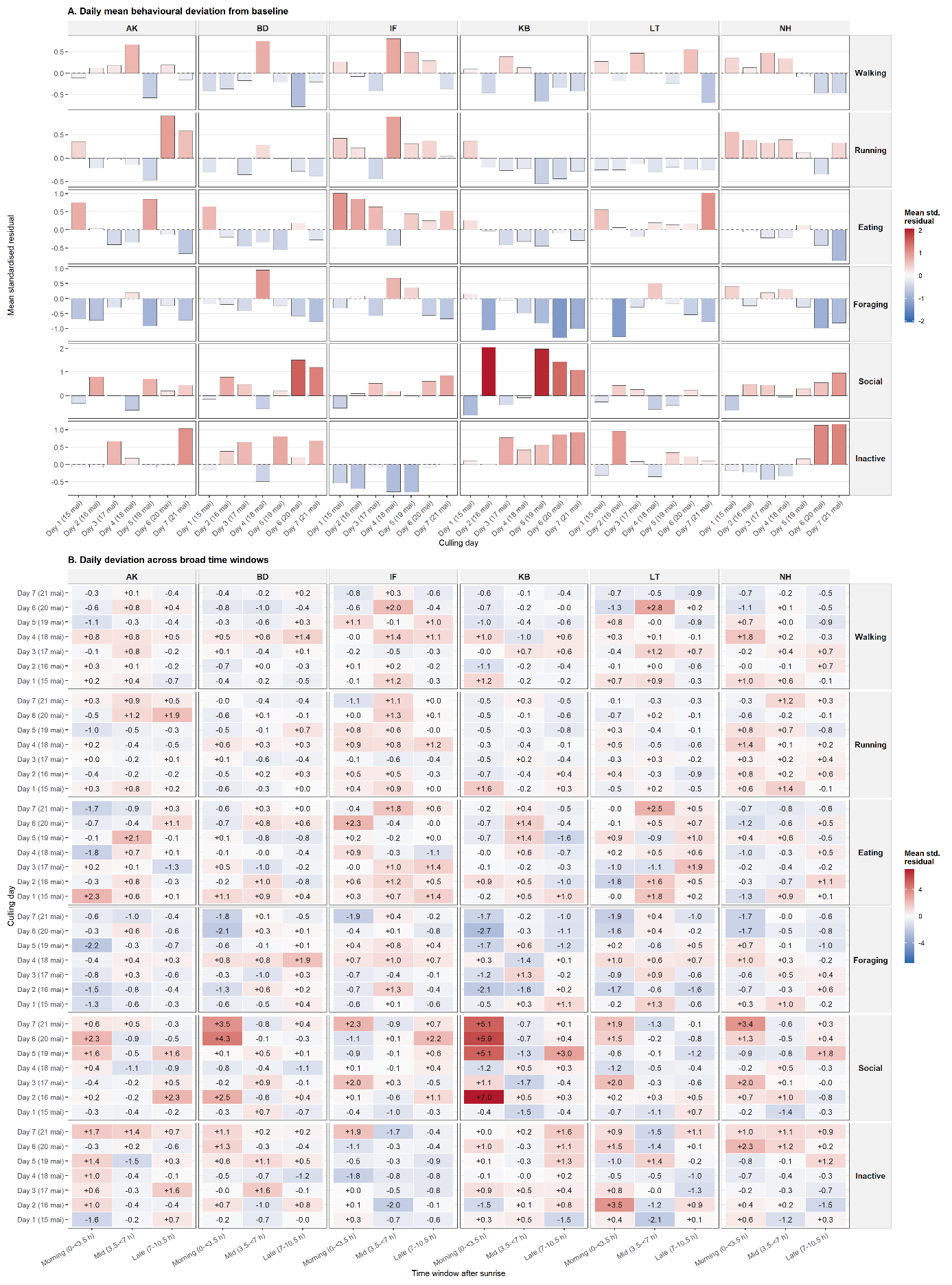

**Figure S6. Standardised residual summaries of acute within-day behavioural deviations during the culling period for all six behavioural categories.** (A) Daily mean standardised residuals across culling days for each group, summarising daily departures from the group-specific baseline expectation. (B) Mean standardised residuals summarised within three time windows after sunrise: morning (0–3.5 h), midday (3.5–7 h), and late afternoon (7–10.5 h). Positive values indicate higher-than-expected activity relative to baseline and negative values indicate lower-than-expected activity. Comparing panels A and B illustrates that some departures were weak at the whole-day scale but pronounced within specific time windows, indicating that acute responses were often temporally concentrated rather than sustained across the full active period. Walking and running are highlighted in the main text; the remaining behaviours are shown here to confirm that their fine-scale deviations were broadly consistent with patterns already captured by the daily and circadian analyses.

##### Figure S7: Within group variability

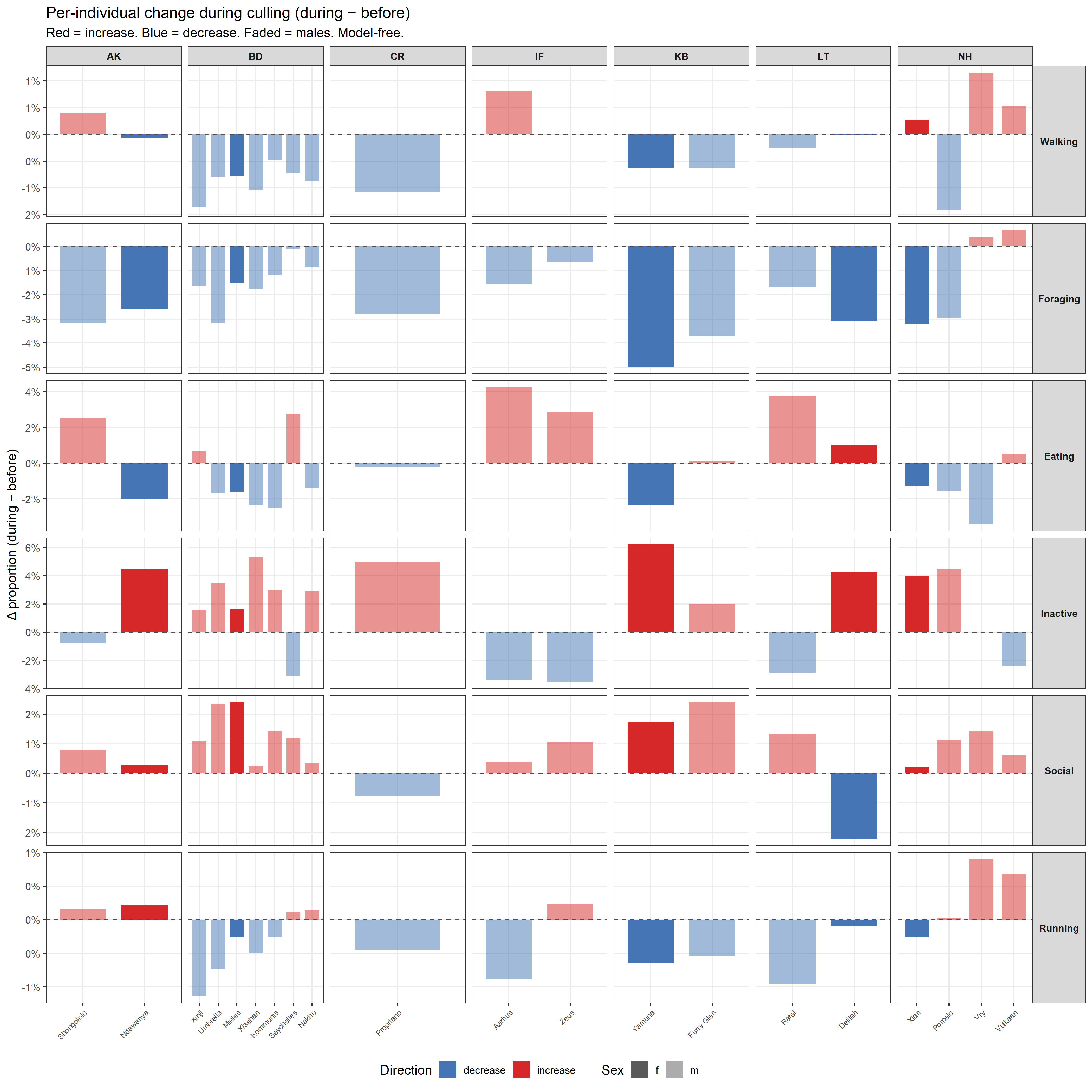

**Figure S7: Individual-level changes in daily behavioural budgets during culling relative to baseline.** Bars represent, for each collared individual, the change in mean daily behavioural proportion during culling relative to the before period (during − before), expressed in percentage points. Positive values indicate increases and negative values decreases during culling. Panels are organised by behaviour and group, with bar colour showing the direction of change and shading indicating sex. Despite some inter-individual variation in response magnitude, individuals within the same group overall showed concordant directional shifts, indicating that the group-level effects were not driven by isolated outliers.

##### Figure S8: Environmental conditions during the study window

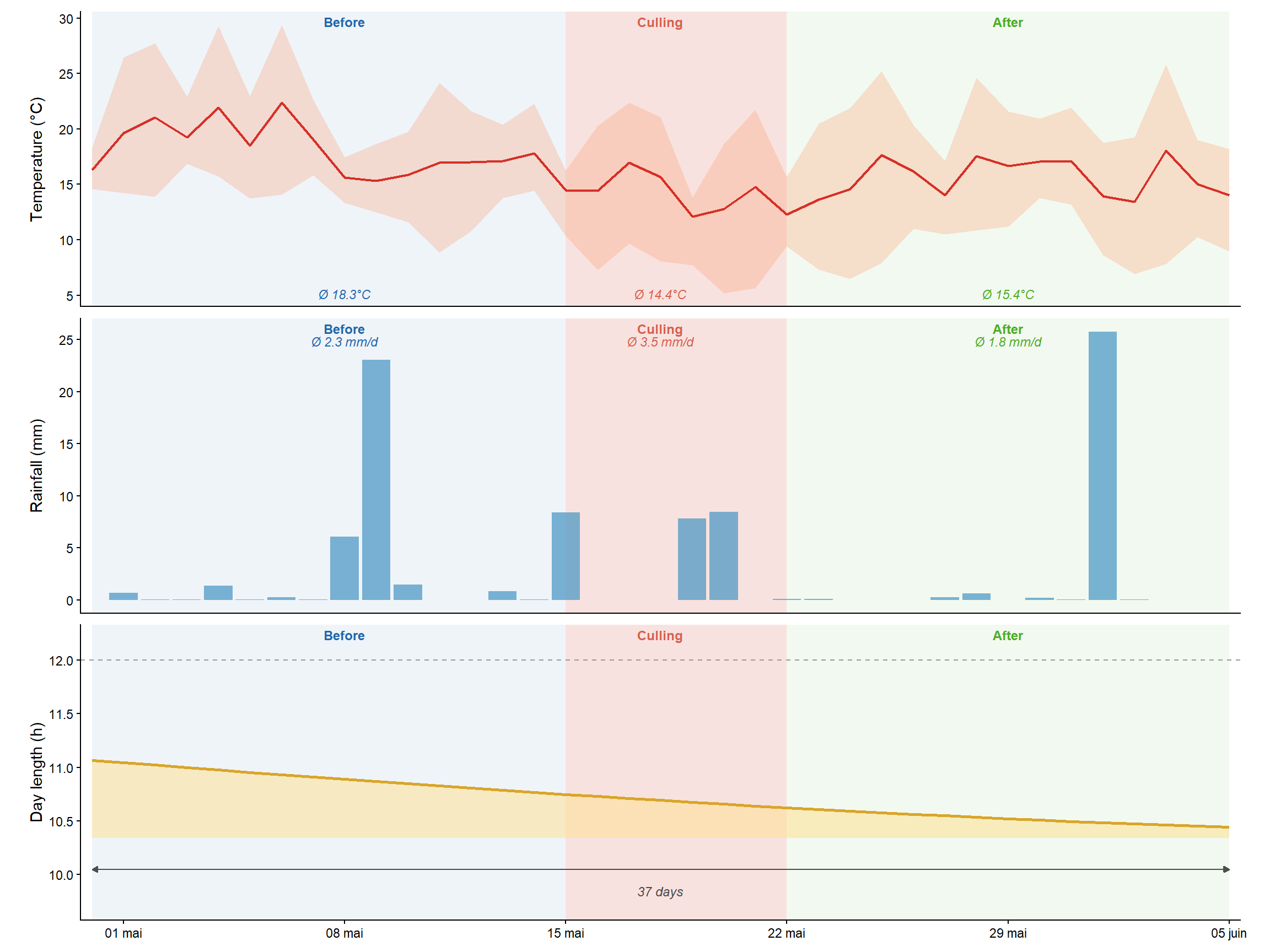

**Figure S8. Environmental conditions during the study window (±15 days around the culling event).** Daily environmental variables are shown for the 37-day period spanning 30 April to 5 June 2023, centred on the culling event (15–21 May 2023). Background shading indicates the three analysis phases: blue = before culling (30 April – 14 May), red = culling period (15–21 May), green = after culling (22 May – 5 June). Temperature and rainfall data were obtained from the ERA5-Land reanalysis dataset (ECMWF; 9 km spatial resolution with hourly temporal resolution), extracted for each GPS collar locations via Google Earth Engine and aggregated to daily values. **(A)** Daily temperature (°C). The solid line shows the daily mean and the ribbon the within-day range across all hourly ERA5-Land estimates. Mean daily temperatures summarized across phases are annotated on the figure. **(B)** Daily rainfall (mm), computed as the sum of hourly ERA5-Land precipitation estimates. Mean daily rainfall (mm/day) is indicated for each phase. **(C)** Day length (hours of daylight) at the study site, computed from astronomical sunrise and sunset times using the *suncalc* R package. The filled area shows daylight hours and the dashed line marks the 12-hour equinox reference. The double-headed arrow spans the full 37-day study window.

##### Figure S9: Mean NDVI within daily MCPs across groups

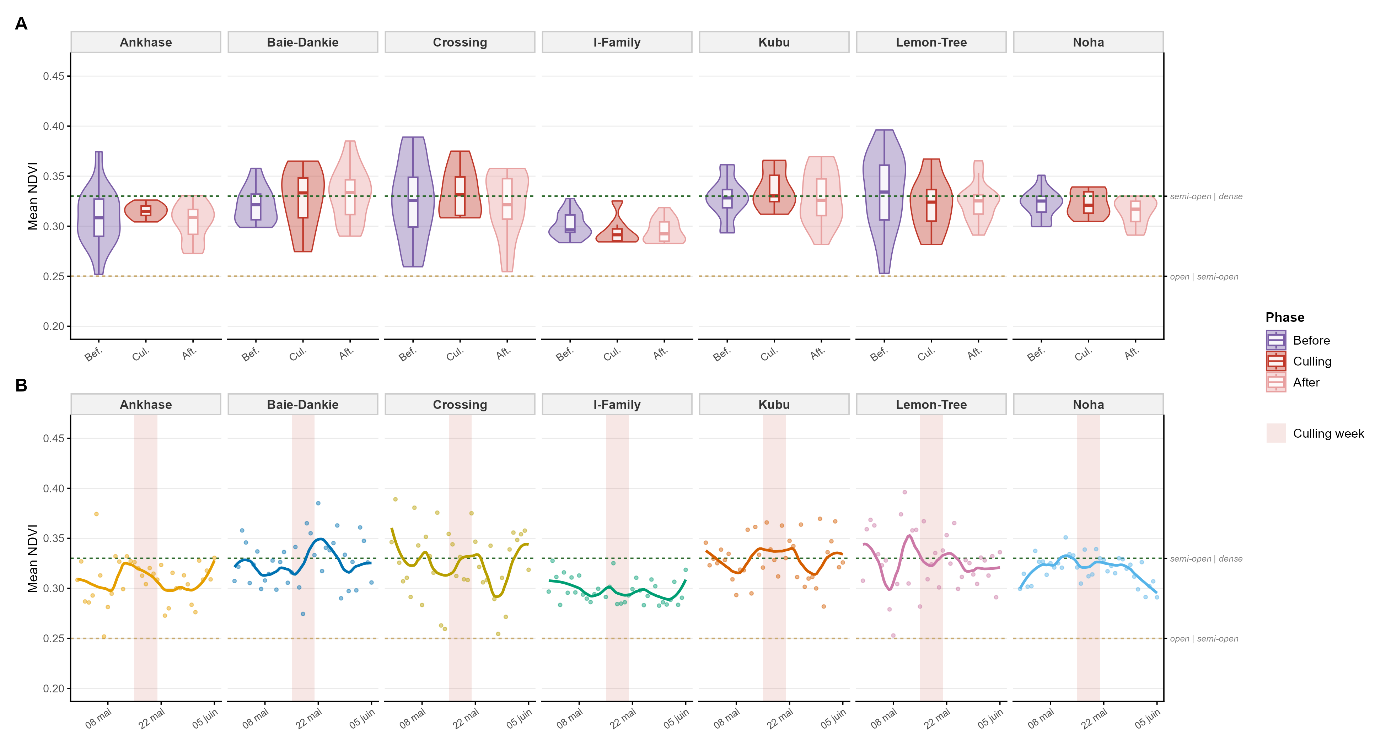

**Figure S9. Daily mean NDVI within 100% MCPs per social group, summarised by phase (A) and across the study period (B).** In both panels, each data point represents one group-day: the mean NDVI across all pixels falling within that day's 100% MCP, computed from a single focal female per group (except I-Family, for which a male was used). (A) Distribution of daily mean NDVI values grouped by phase (Before, Culling, After), shown as violin plots scaled proportionally to density with embedded boxplots (median, interquartile range). (B) Daily mean NDVI values (points) and LOESS trend (line) across the 30-day study window (±15 days around culling). Red shading indicates the culling week (15–21 May 2023). Dashed lines in both panels indicate NDVI classification thresholds: open (< 0.25) and semi-open | dense (≥ 0.33), derived from a 10 × 10 m Sentinel-2 raster smoothed with a 60 m majority filter.
